# Expanding Macrocyclic Topology through Cysteine-to-N-Terminal Cyclisation Enables Covalent Peptide Inhibitor Discovery

**DOI:** 10.64898/2026.09.04.749435

**Authors:** Ella Sawtell, Marina Barrueco, Jonathan Whiteside, Christopher Williams, Maisem Laabei, Scott Lovell

## Abstract

Macrocyclic peptides are an attractive therapeutic modality capable of engaging challenging protein targets while retaining many favourable drug-like properties. Their high-affinity binding also provides an ideal framework for proximity-driven covalent inhibition through incorporation of latent electrophiles. Phage display enables the high-throughput screening of billion-member macrocyclic peptide libraries; however, existing libraries rely predominantly on cysteine-mediated cyclisation, restricting the range of macrocyclic topologies available for ligand discovery.

Here, we report a mild and efficient cyclisation strategy based on a bromomethyl picolinaldehyde (BMP) linker that reacts with a cysteine side chain and the peptide N-terminus to generate a previously unexplored macrocyclic topology incorporating neighbouring pyridine and imidazolidinone rings. The chemistry is compatible with phage display and enabled screening of BMP-cyclised peptide libraries against plasma kallikrein, yielding a potent macrocyclic inhibitor. The BMP-cyclised peptide displayed substantially greater potency than analogous peptides cyclised through either a disulfide bond or the widely used linker 1,4-bis(bromomethyl)benzene (DBMB). Furthermore, comparison with an equivalent DBMB-cyclised library demonstrated that BMP-mediated cyclisation enabled access to binding motifs not identified by conventional cysteine-to-cysteine cyclisation.

Finally, positional sulfur(VI) fluoride exchange (SuFEx) electrophile scanning converted the BMP-derived hit into a selective covalent macrocyclic activity-based probe capable of labelling plasma kallikrein in human plasma. Together, these findings establish BMP-mediated cyclisation as a versatile strategy for expanding the topological diversity of phage-displayed macrocycles and accelerating the discovery of both reversible and covalent macrocyclic peptide ligands.

**TOC Caption:** A strategy for the discovery of covalent BMP-macrocyclised peptide inhibitors.

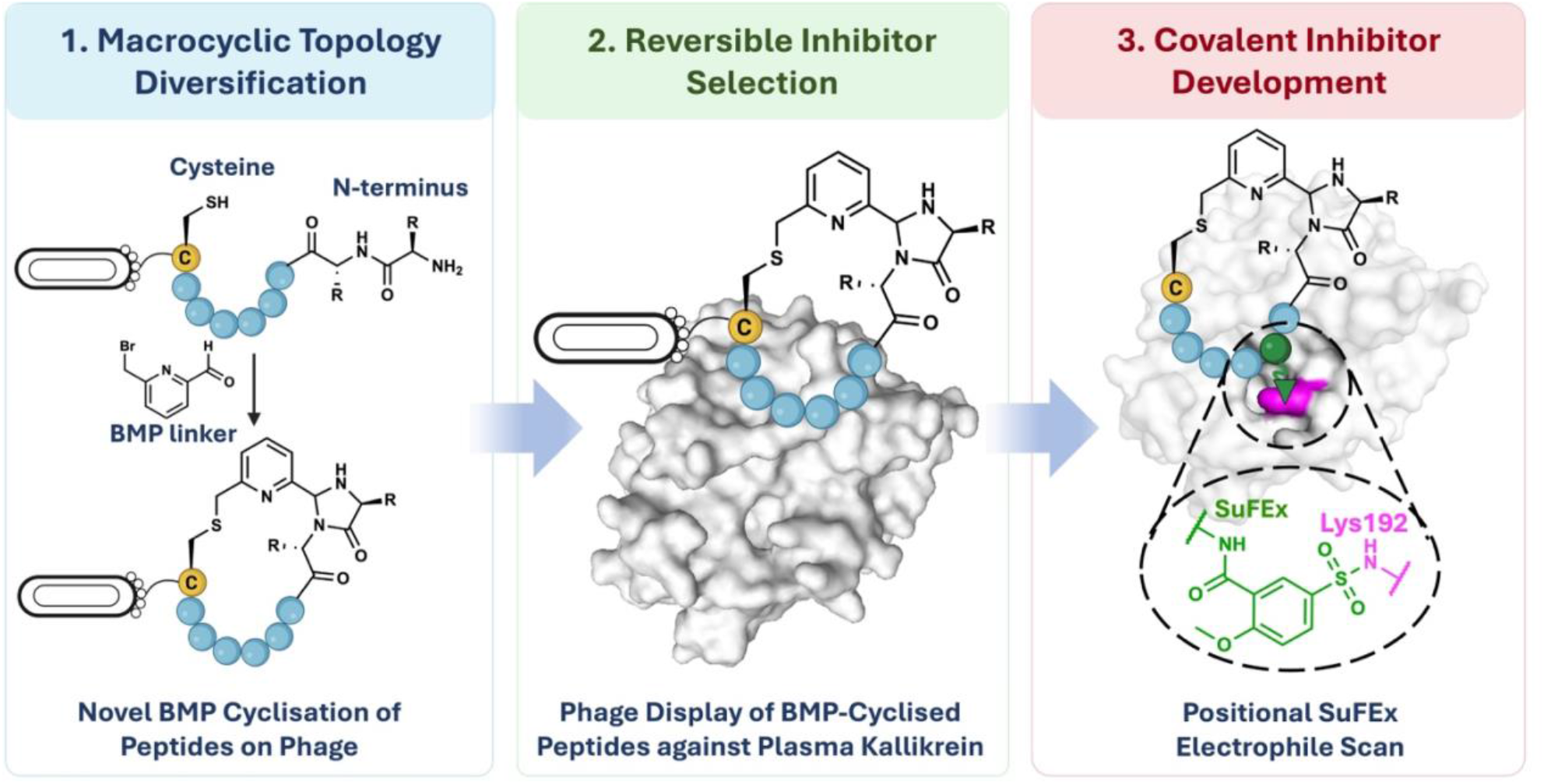

## Introduction

Macrocyclic peptides have emerged as a powerful class of therapeutic agents that sit in the chemical space between small molecules and large biologics. This affords them sufficient size and complexity to bind challenging protein targets for which small-molecule drugs are unsuited, whilst avoiding the poor tissue penetration and unfavourable immune responses associated with large biologics.^1, 2^ The conformational pre-organisation generated by cyclisation of the peptide backbone is particularly important for improving binding affinity, maximising specificity, and reducing susceptibility to proteolytic degradation compared to linear counterparts.^3^

The high binding affinity achieved by macrocyclic peptides makes them an ideal framework for incorporation of proximity-driven covalent inhibition. The strong non-covalent binding of the macrocyclic peptide scaffold precisely positions a weakly reactive electrophile for selective and irreversible target modification.^4-8^ The resultant permanent target engagement then gives enhanced potency, improved selectivity and reduced dosing intervals.^9^ As such, covalent macrocycles are emerging as a powerful modality for selectively engaging disease-relevant protein targets across mammalian, bacterial, and viral systems.

Phage display is a powerful technique that enables the generation and high-throughput screening of billion-member peptide libraries. Genetically encoded peptides, varying in amino acid composition, are displayed on the surface of bacteriophages and, for macrocyclic peptide discovery, are cyclised prior to iterative rounds of selection against protein targets of interest. The genotype–phenotype link between phage DNA and their displayed peptides enables rapid amplification and subsequent identification of enriched peptide binders.^10,11^ To date, phage-displayed peptides are predominantly cyclised at cysteine side chains. This is mostly achieved through spontaneous disulfide bond formation during phage assembly or by post-translational modification with cysteine-reactive chemical linkers (**Fig. 1a**).^12^ While these approaches are robust and have yielded numerous high-affinity ligands, they rely on libraries containing multiple fixed positions cysteines and thus generate macrocycles with broadly similar topologies.^13-16^

**Figure 1:**
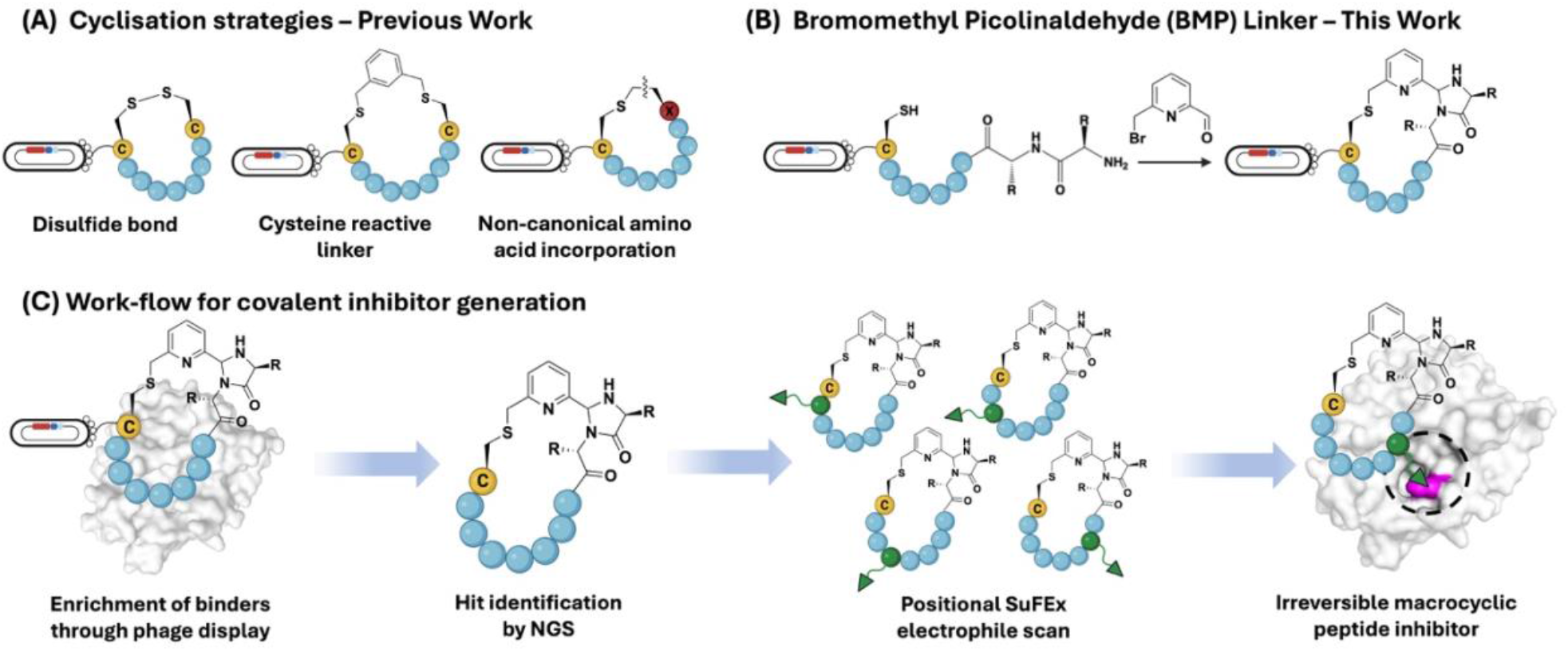
BMP-mediated cyclisation strategy expands macrocyclic topology in phage display and enables covalent inhibitor development. (A) Existing cyclisation strategies for phage-displayed peptides, including disulfide bond formation, cysteine-reactive chemical linkers and incorporation of non-canonical amino acids. (B) Bromomethyl picolinaldehyde (BMP) linker developed in this work enables N-terminus-to-cysteine cyclisation. (C) Workflow for covalent inhibitor development: BMP-cyclised peptide phage display libraries are screened to identify macrocyclic peptide binders, after which electrophile scanning within the hit scaffold enables identification of positions capable of covalent target engagement.

It is well understood that overall macrocyclic topology, including connectivity, rigidity, and three-dimensional shape, can play a decisive role in determining binding affinity and selectivity.^3, 17, 18^ Expanding the repertoire of cyclisation chemistries compatible with phage display offers a powerful route to increasing structural diversity in terms of macrocyclic topology. This understanding has driven a recent development of novel cyclisation strategies for phage-displayed peptides utilising non-cysteine reactivity.^12^

Broadly, these strategies can be divided into those utilising genetically encoded non-canonical amino acids and those based on post-translational modification with chemical linkers. Although non-canonical amino acid approaches have produced high-affinity macrocyclic peptides, the complexity of library generation and associated loss of diversity remain key limitations.^19-22^ Consequently, several post-translational cyclisation strategies have recently emerged, including chemistries targeting the peptide N-terminus, lysine side chains, and N-terminal serine residues.^23-27^ Many of these strategies require harsh reaction conditions, lengthy or multistep procedures, or generate structures unstable at room temperature. These limitations can compromise phage viability and cyclisation efficiency, ultimately reducing the diversity of macrocyclic peptides screened. Therefore, expanding the repertoire of cyclisation strategies compatible with phage display remains an important goal.

In this work, we report a novel bromomethyl picolinaldehyde (BMP) linker that cyclises phage-displayed peptides through reaction with a single fixed-position cysteine residue and the peptide N-terminus, generating a previously unexplored macrocyclic topology that incorporates a five-membered imidazolidinone ring (**Fig. 1b**). Phage panning with BMP-cyclised libraries enabled the identification of potent macrocyclic peptide inhibitors of plasma kallikrein, highlighting the value of accessing new macrocyclic topologies. Building on hit peptide scaffolds, we incorporated sulfur(VI) fluoride exchange (SuFEx) electrophiles to generate covalent macrocycle analogues (**Fig. 1c**).^28, 29^ Electrophile scanning identified a covalent active-site inhibitor, which was subsequently deployed as an activity-based probe for monitoring plasma kallikrein activity in blood.

## Results and discussion

### Linker Design and Validation on Synthetic Peptides

Building on chemistry developed by MacDonald et al. for site-specific protein modification, we synthesised a chemical-linker centred around a picolinaldehyde moiety.^30^ We anticipated that rapid reaction of the bromomethyl group with a single cysteine side chain would position the aldehyde in proximity to the peptide N-terminus, promoting formation of an imidazolidinone ring (**Fig. 2A**). This generates a distinct macrocyclic topology, incorporating neighbouring pyridine and imidazolidinone rings while simultaneously masking the peptide N-terminus, a common strategy for improving peptide stability. Formation of the imidazolidinone proceeds through a planar imine intermediate, enabling generation of two diastereomeric macrocycles from a single peptide sequence (**Fig. S1**). Although such diastereomer formation can complicate hit purification and characterisation, we were interested to explore the increased structural diversity this generates within phage display libraries.

**Figure 2:**
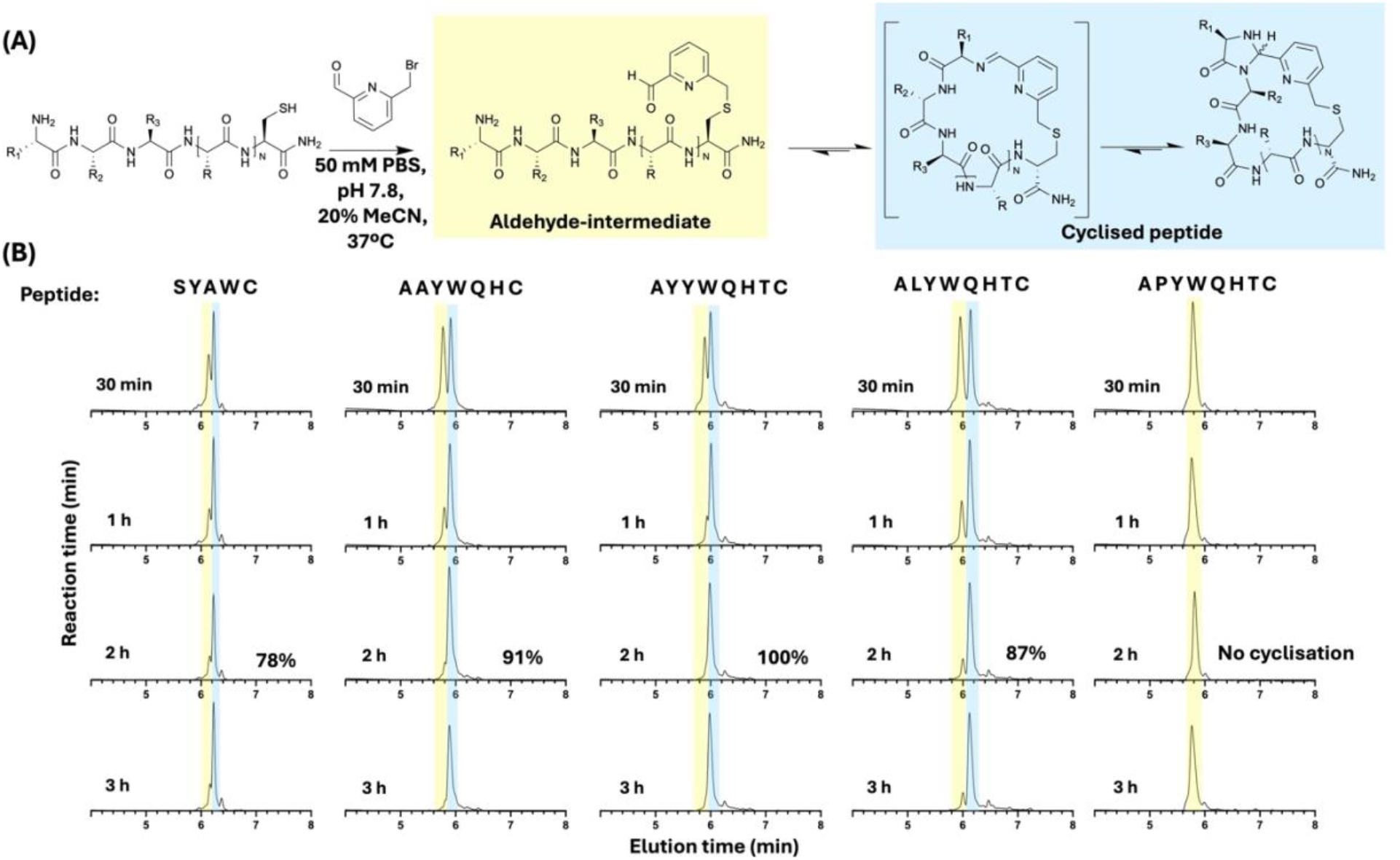
Cyclisation of model peptides with the bromomethyl picolinaldehyde linker. (A) Mechanism and selected conditions (PBS (pH 7.8), 20% MeCN, 37 oC) for cyclisation. (B) UV traces (280 nm) from monitoring the cyclisation of model synthetic peptides under the selected conditions. The alkylated aldehyde-intermediate and cyclised peptide species are highlighted for each trace (yellow and blue, respectively). The percentage of cyclised peptide relative to the total peptide signal after 2 hours is displayed on each trace. This was calculated by integrating the area under the curve of the corresponding UV peaks.

Cyclisation was first evaluated on synthetic peptides varying in length and amino acid composition. Peptides were of the form X_n_C, where X represents any amino acid other than cysteine and n = 4-7. Cyclisation was observed under all the conditions tested, including a range of aqueous buffers containing 20% acetonitrile for linker solubilisation, at pH 7.8-9 and at both room temperature and 37°C. However, cyclising in PBS at pH 7.8 and 37°C gave the most rapid cyclisation and was therefore selected for subsequent studies. To minimise sequence-dependent variation in cyclisation efficiency, the N-terminal residue was fixed as alanine, which we and others have previously found to robustly react with picolinaldehyde moieties.^30^ Cyclisation reactions were monitored by LC-MS and the extent of cyclisation at fixed time points quantified by UV peak integration (**Fig. 2B**). No attempt was made to separate diastereomers at this stage. Reaction monitoring confirmed rapid alkylation of the cysteine side chain, with complete modification observed for all peptides at the first time point analysed (30 min). This was followed by slower reaction of the picolinaldehyde with the N-terminal amine to yield fully cyclised products. For peptides in which the alkylated and cyclised species could be resolved chromatographically, cyclisation efficiencies of ≥78% were observed after 2 h (**Fig. 2B**). Although picolinaldehyde moieties are susceptible to hydrate formation, this was only minimally observed for the alkylated intermediates.

Reaction of the picolinaldehyde with the peptide N-terminus can be monitored by LC-MS due to a mass decrease of 18 Da. However, the cyclised imine intermediate and final imidazolidine product are isomeric and therefore cannot be distinguished by mass spectrometry alone. As an initial negative control, the peptide APYWQHTC was modified and monitored by LC-MS. While the free N-terminus should permit formation of the cyclised imine intermediate, incorporation of proline in the N-terminal-1 position removes the backbone amide nitrogen required for imidazolidinone formation (**Fig. S1**). Complete alkylation of the cysteine residue was observed, but no cyclised imine species was detected by LC-MS over 24 hours. This suggests the transience of the imine intermediate under the cyclisation tracking conditions employed (**Fig. 2B**). To further investigate the nature of the cyclised products, the model peptide ALYWQHTC was cyclised for 2 hours, and a species with the expected mass for the cyclised peptide was purified for structural characterisation. The structure was analysed by 1D and 2D NMR spectroscopy and key correlations confirmed connectivity from the cysteine side chain, through the pyridine linker, to the imidazolidinone formed by reaction with the N-terminal alanine (**Fig. 3A** and **Fig. S2-20)**. Notably, NOESY correlations between the cysteine β-protons and the linker CH_2_ group confirmed alkylation of the cysteine side chain (**Fig. 3B**). In addition, a singlet peak at 5.39 ppm, consistent with the characteristic CH proton of the imidazolidinone ring^31^, showed through-space interactions with pyridine protons (**Fig. 3C**) and through-bond correlations with carbons of the N-terminal alanine (**Fig. 3D**). No signals consistent with the imine intermediate were observed. Together, these data support formation of the imidazolidinone structure proposed from the LC-MS studies above.

**Figure 3:**
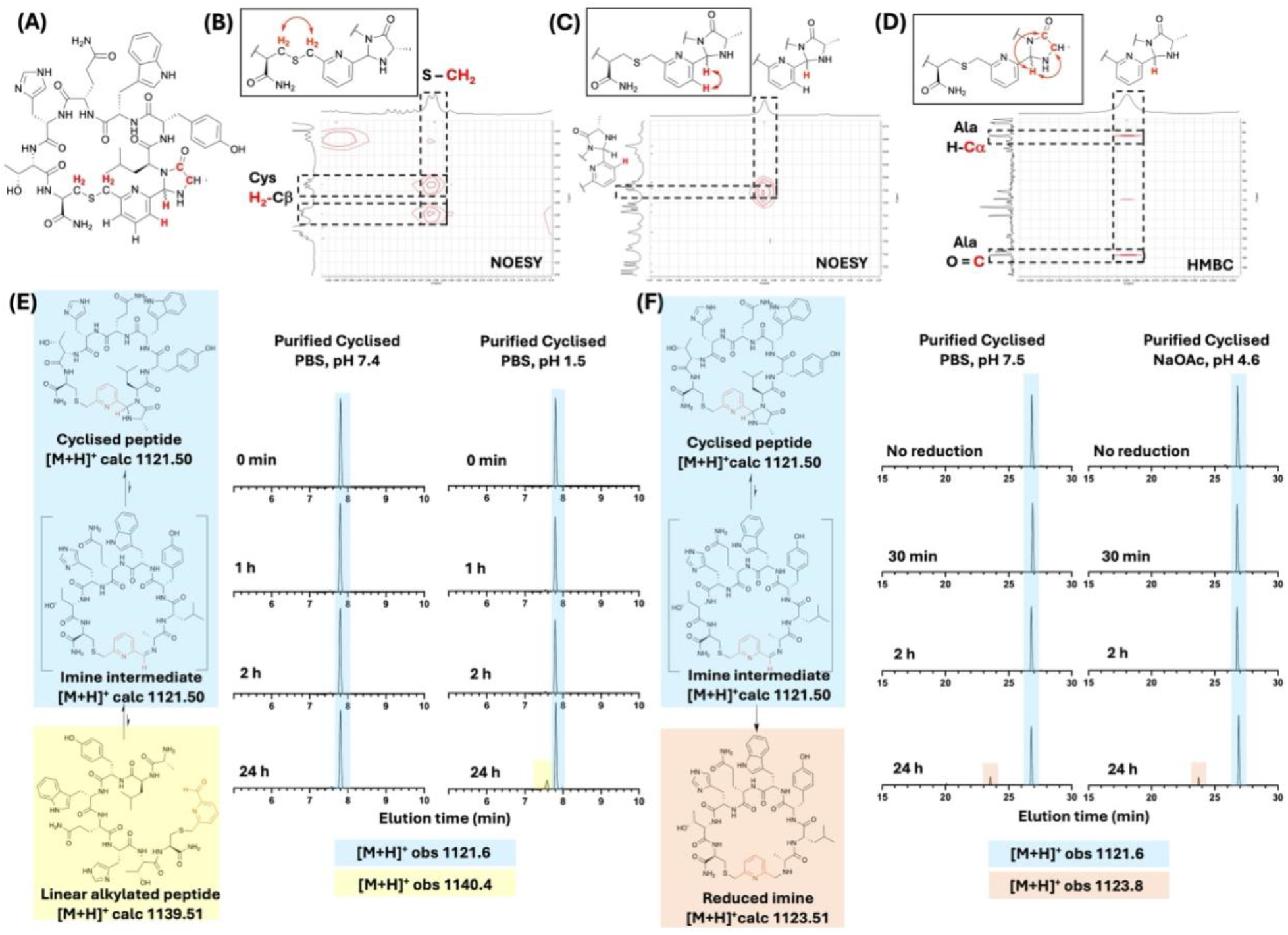
Characterisation of cyclised model peptide ALYWQHTC. (A) Structure of peptide with key protons for confirming linker connectivity highlighted in red. (B) Excerpt from the NOESY spectrum of cyclised ALYWQHTC showing interaction between the β-protons of the cysteine and the CH_2_ group of the linker. (C) Excerpt from the NOESY spectrum of cyclised ALYWQHTC showing interaction between a pyridine proton of the linker and the CHproton of the imidazolidinone. (D) Excerpt from HMBC spectrum of cyclised ALYWQHTC showing interaction between the imidazolidinone CH and the carbons of the N-terminal alanine, within the imidazolidinone. (E) UV traces (280 nm) from monitoring reversion of the cyclised peptide to the linear alkylated intermediate (pH 7.4 or pH 1.5, RT). (F) UV traces (280 nm) from monitoring reversion of the imidazolidinone structure to the imine intermediate by irreversible reduction to the secondary amine (25 equiv. NaCNBH_3_, RT, pH 7.5 or pH 4.6, RT).

To assess the stability of the imidazolidinone structure formed, the cyclised peptide ALYWQHTC was monitored for reversion to the linear alkylated intermediate under neutral and acidic conditions at room temperature (**Fig. 3E** and **Fig. S21-22)**. Minimal reversion to the aldehyde intermediate was observed, even under highly acidic conditions, demonstrating that the macrocycles were sufficiently stable for the timescale of a phage display experiment. This finding supports previous reports that the pyridine nitrogen promotes imidazolidinone formation and enhances its stability relative to analogous benzaldehyde-derived systems.^25,30^ We next investigated potential reversion of the imidazolidinone to the imine intermediate by incubating the cyclised ALYWQHTC peptide with sodium cyanoborohydride under both neutral and acidic conditions. This should enable any imine formed to be irreversibly trapped as the reduced secondary amine, causing a +2 Da mass change.^23, 25, 30^ Only minimal trapping of the reduced imine species was observed after 24 hours incubation, consistent with slow reversion to the imine intermediate (**Fig. 3F** and **Fig. S23**). To confirm that the reduction conditions were suitable for trapping formed imine, sodium cyanoborohydride was added to a crude cyclisation reaction after 5 min at 37 °C, a time point at which alkylation had occurred, but cyclisation remained incomplete. Under these conditions, reduced imine and reduced alkylated species were readily observed with minimal further cyclisation detected following addition of the reductant (**Fig. S24**). Together, these data support the stability of the imidazolidinone macrocycles under conditions relevant to downstream phage display experiments.

### Validation of BMP Cyclisation on pIII-Displayed Peptides

Having established efficient cyclisation of model synthetic peptides and the stability of the resulting macrocycles, we next evaluated the chemistry in a system more representative of phage-displayed peptides. In phage display, peptides are displayed on the five copies of the phage coat protein III (pIII protein). Therefore, we expressed the model peptide AAYWQHTC fused to the N1-N2 domains of a disulfide-free pIII protein, with a C-terminal His-tag for purification.^32^ Expression and purification of the fusion protein were confirmed by SDS-PAGE and mass spectrometry (**Fig. S25**).

Given the rapid cysteine alkylation observed for the synthetic peptides, the fusion protein (20 µM) was first incubated with BMP-linker (1 mM) in PBS with acetonitrile (20%) for just 1 h at 30 °C (**Fig. 4A**). Complete alkylation of the displayed peptide was confirmed by mass spectrometry (**Fig. 4B**). Excess linker and acetonitrile could then be removed by buffer exchange into PBS before incubation at 37 °C to promote cyclisation under conditions expected to minimise phage toxicity. Mass spectrometry analysis revealed 73% and 77% cyclisation after one and two hours at 37 °C, respectively (**Fig. 4B**). As only a modest increase in cyclisation was observed during the second hour, subsequent experiments utilised a protocol comprising 1 h alkylation at 30 °C, buffer exchange, and a further 1 h incubation at 37 °C.

**Figure 4:**
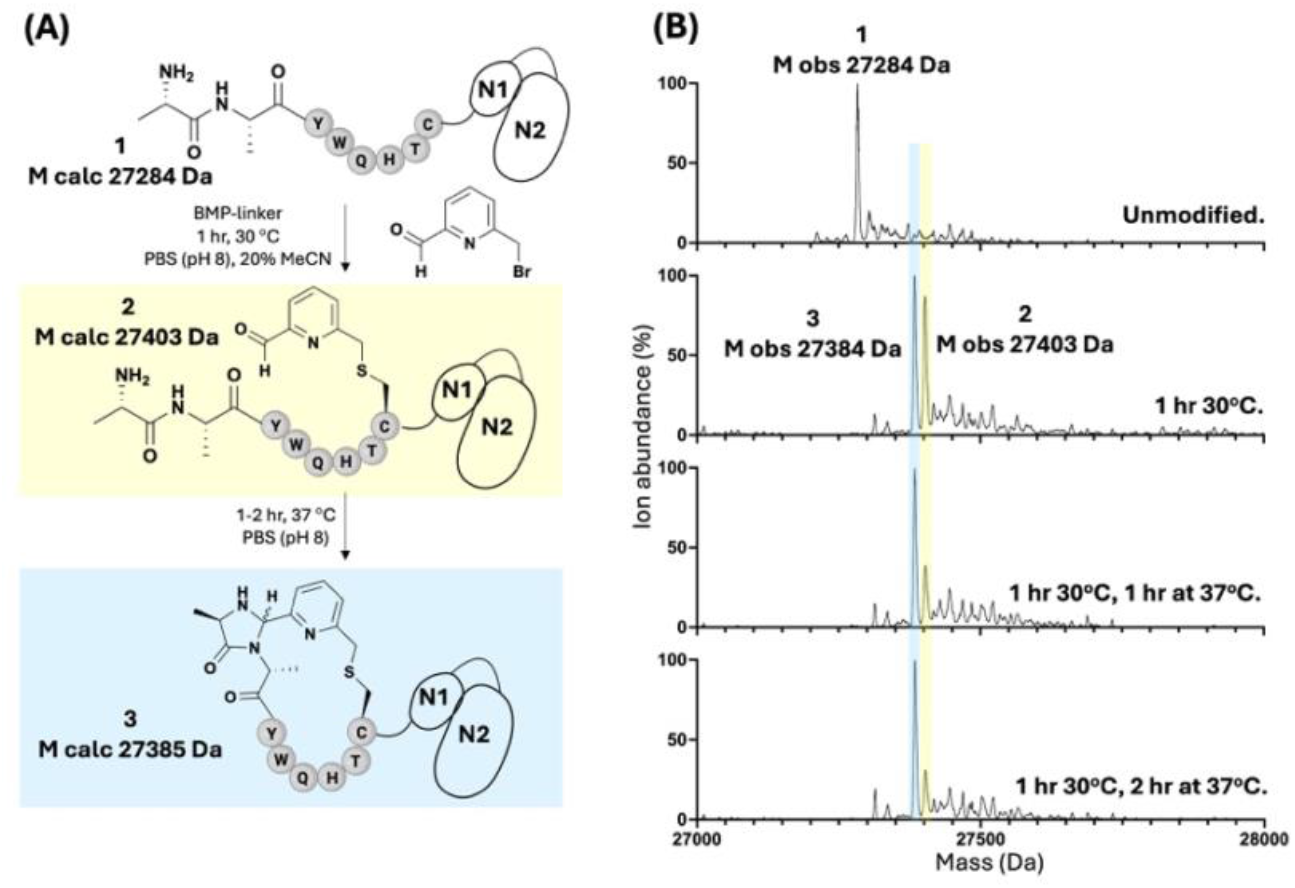
Confirming cyclisation of the test peptide AAYWQHTC with the bromomethyl picolinaldehyde linker when displayed on the phage pIII protein. (A) Scheme showing cyclisation of the displayed peptide. The BMP linker was added with 20% MeCN to allow for linker solubilisation. Full peptide alkylation was observed after 1 h at 30°C allowing excess linker and MeCN to be removed. The reaction was then heated at 37 °C for a further 2 h, pushing the cyclisation towards completion with minimised toxicity risk. (B) Mass spectra from LC-MS analysis of the cyclisation reaction. The linear alkylated and cyclised peptide species are highlighted (yellow and blue, respectively). Spectra are shown for the AAYWQHTC-fusion protein when: unmodified, after linker addition at 30 °C for 1 h (with 20% MeCN and excess linker), after a further 1 and 2 h at 37 °C (following MeCN and excess linker removal).

We next evaluated these conditions on a proof-of-concept fd phage library displaying peptides of the form AX_7_C (where X=any canonical amino acid), generated utilising degenerate NNK primers and a whole-plasmid PCR method.^33^ The fd phage plasmid utilised confers chloramphenicol resistance and as such phage viability was assessed throughout the modification procedure by infecting TG1 cells and quantifying chloramphenicol-resistant colonies. No significant loss of phage titre was observed following TCEP treatment, linker alkylation, or the subsequent cyclisation step after removal of excess linker and acetonitrile, demonstrating that the BMP cyclisation protocol is well tolerated by phage (**Fig. S26**).

### Generation of a Phage Library for Screening Against Plasma Kallikrein

Having established that BMP cyclisation is compatible with phage particles, we next sought to determine whether the resulting macrocyclic libraries could be used for ligand discovery. As a proof-of-concept target, we selected plasma kallikrein (PK), a serine protease involved in the kinin-kallikrein, complement and coagulation pathways. Dysregulated PK activity contributes to diseases such as hereditary angioedema and diabetic macular oedema, and several macrocyclic peptide PK inhibitors have previously been identified through phage display selections.^14, 34-36^

As above, a whole-plasmid PCR method was used to generate a phage library displaying peptides of the form AX_7_C (where X=any canonical amino acid), suitable for cyclisation with the BMP linker and comparable in size to previously reported PK-binding peptide inhibitors.^33, 36^ Given the strong preference of PK for arginine at the S1 binding site,^37^ the library was constructed using seven degenerate AX_7_C primers, each encoding arginine at a different variable amino acid position. This ensured that every displayed peptide contained a single arginine residue while allowing its position to vary throughout the sequence (**Fig. 5A**). In addition to biasing the library towards PK recognition, this design reduced the theoretical sequence space, increasing the likelihood of comprehensive library coverage and hence identification of high-affinity binders. In total, 5.55 x10^8^ TG1 cells were transformed during library construction, providing theoretical coverage of the 4.48 x10^8^ possible peptide sequences.

**Figure 5:**
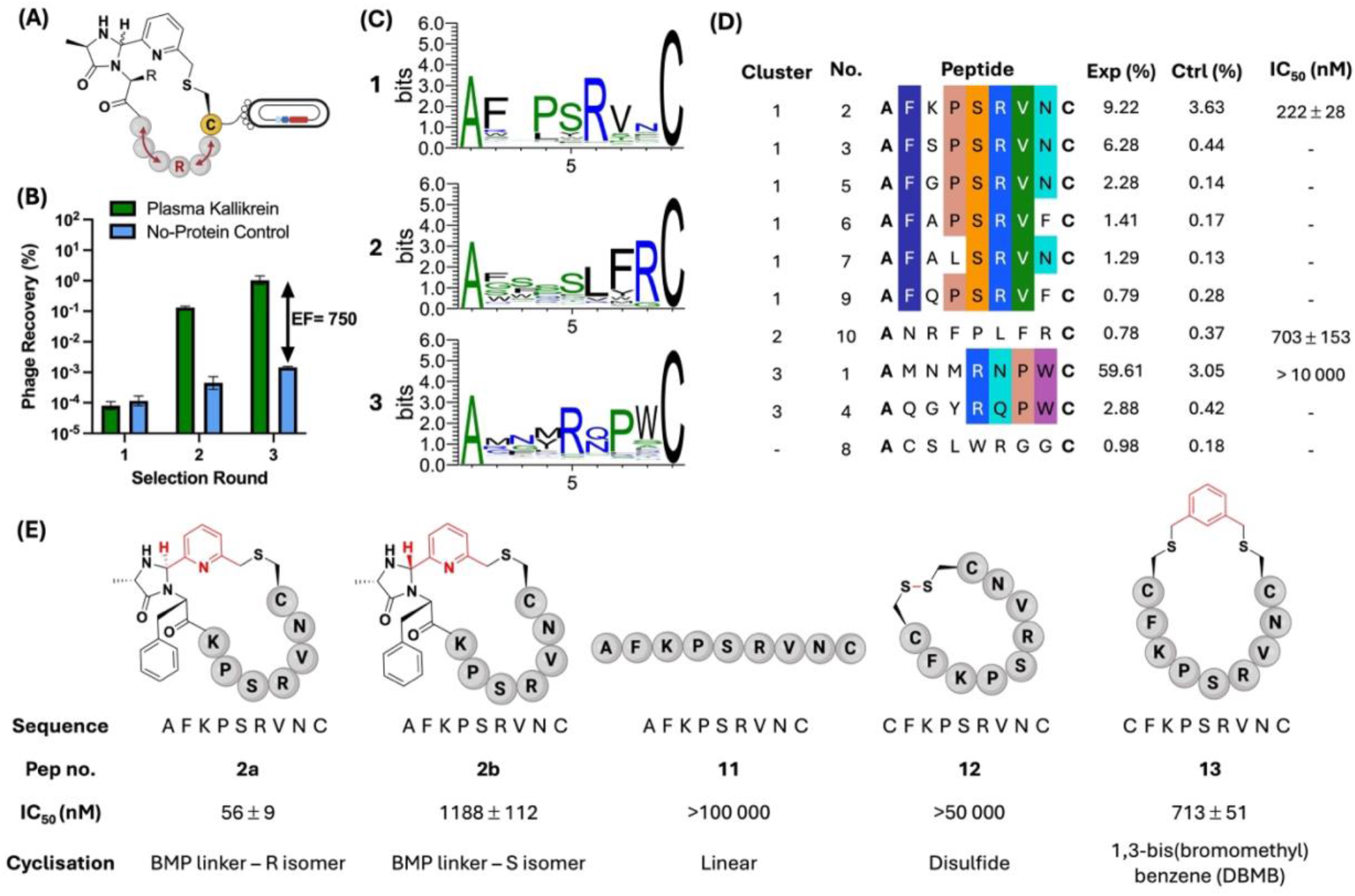
Proof-of-concept screening against plasma kallikrein. (A) Format of the BMP-cyclised AX_7_C library. (B) Percentage recovery of the BMP-cyclised AX_7_C phage over three rounds of selection against plasma kallikrein. An enrichment factor (EF) of 750 was seen after the third round of selection. (C) Logo plots showing the consensus sequences obtained after GibbsCluster analysis of the enriched peptide sequences. (D) The top ten peptide sequences from the NGS data of the third round of selection against plasma kallikrein. The percentage abundance of each peptide sequence from the selection against plasma kallikrein and the no-protein control are given (Exp and Ctrl, respectively). An IC_50_ for the most abundant peptide representative of each of the three consensus sequences was calculated. (E) The most potent peptide, peptide **2**, was synthesised and tested for plasma kallikrein inhibition as the two BMP-cyclised diastereomers, a linear peptide sequence, cyclised with a disulfide bond and cyclised with the commercial linker DBMB.

The BMP-cyclised library was subjected to three rounds of selection against immobilised PK, with a pre-clearing step against the bead matrix used for target immobilisation (streptavidin- or neutravidin-coated magnetic beads) before each round and increasing wash stringency throughout the selection process. For each round, a parallel selection was performed against a no-protein control to monitor non-specific enrichment. Phage retained from both positive and negative selections were quantified, but only the phage captured in the positive selection were amplified and carried through to the next round of selection. After the third round, approximately 750-fold enrichment was observed for PK relative to the control selection (**Fig. 5B**). Next generation sequencing of the final outputs identified 895 sequences enriched against PK, which were subsequently analysed using GibbsCluster to identify consensus binding motifs (**Fig. S27**).^38^

### Validation of Plasma Kallikrein-Enriched Peptides

GibbsCluster analysis of the enriched sequences identified three clear consensus motifs (**Fig. 5C**). The most abundant peptide representative of each motif was synthesised, cyclised with the BMP-linker and evaluated for inhibition of PK (**Fig. 5D** and **Fig. S28**). No attempt was made to separate the cyclisation diastereomers for each peptide at this stage. Of the three peptides tested, peptide **2** was the most potent, exhibiting an IC_50_ of 220 nM. Optimisation of the chromatographic purification of peptide 2 enabled subsequent isolation of the individual R/S diastereomers, which were characterised by NMR and NaCNBH_3_ reduction studies (**Fig. S29-38**). Evaluation of the isolated species revealed more than 20-fold difference in potency, with the R isomer (peptide **2a**) displaying an IC_50_ of 56 nM (**Fig. S39**). Although formation of diastereomers complicates hit purification and characterisation, this result demonstrates the benefit of generating additional diversity through use of a linker able to form two distinct structures from a single peptide sequence.

To investigate the importance of the BMP-derived topology, peptide **2a** was also evaluated as a linear peptide and as analogues cyclised either through a disulfide bond or with the widely used linker 1,4-bisbromomethyl benzene (DBMB) (peptides **11, 12** and **13**, respectively) (**Fig. 5E** and **Fig. S28**). For the disulfide- and DBMB-cyclised peptides, the N-terminal alanine was replaced with cysteine to enable cyclisation. The linear peptide showed no measurable inhibition of PK, consistent with enrichment occurring only after cyclisation of the displayed peptides. Both the disulfide- and DBMB-cyclised analogues were significantly less potent than the BMP-cyclised peptide, demonstrating the importance of the specific macrocyclic topology generated by the BMP-linker for productive target engagement.

The improved potency of the BMP-cyclised peptide led us to ask whether the altered macrocyclic topology also enabled access to different binding motifs during phage display. Comparison with an analogous selection against PK performed using a DBMB-cyclised library of equivalent peptide length revealed that the F-PSRVN motif represented by peptide **2** was not enriched in the DBMB library.^36^ Thus, beyond improving the activity of individual peptide scaffolds, the BMP linker appears to expand the range of ligands accessible by phage display through generation of previously unexplored macrocyclic topologies.

Finally, we assessed whether replacing the commonly used DBMB linker with the BMP scaffold altered blood compatibility. The BMP-cyclised peptide (**2a**), together with the corresponding disulfide- and DBMB-cyclised analogues (peptides **12** and **13**), was incubated with horse erythrocytes for 1 h after which haemolysis was quantified (**Fig. S40**). All three peptides exhibited minimal haemolysis (<10%) at 100 μM. Notably, the BMP-cyclised peptide consistently displayed lower haemolysis than its DBMB-cyclised counterpart, suggesting that the BMP linker maintains, and may modestly improve, the blood compatibility of macrocyclic peptides.^39^

### SuFEx Scan of Peptide 2 for Covalent macrocycle Development

Having identified peptide **2** as a potent BMP-cyclised inhibitor of PK, we next sought to determine whether this scaffold could be developed into a covalent inhibitor. Covalent inhibitors offer prolonged target engagement and can be converted into activity-based probes for selective detection of enzyme activity in complex biological samples.^4, 40^ SuFEx electrophiles are latent covalent warheads capable of reacting with nucleophilic residues including lysine, histidine and tyrosine. Their broad amino acid reactivity and compatibility with cysteine-based cyclisation strategies make them attractive electrophiles for covalent peptide development.^28^

As the binding mode of peptide **2** is unknown, we performed an electrophile scan to identify positions within the macrocycle from which a SuFEx electrophile could engage a proximal nucleophilic residue on PK. A para-methoxy-substituted aryl sulfonyl fluoride^28^ was therefore incorporated at each residue position other than the proline, which is expected to be important for the overall structure of the peptide, and the P1 and P2 positions, which are expected to be critical for active site binding (**Fig. 6A**).

**Figure 6:**
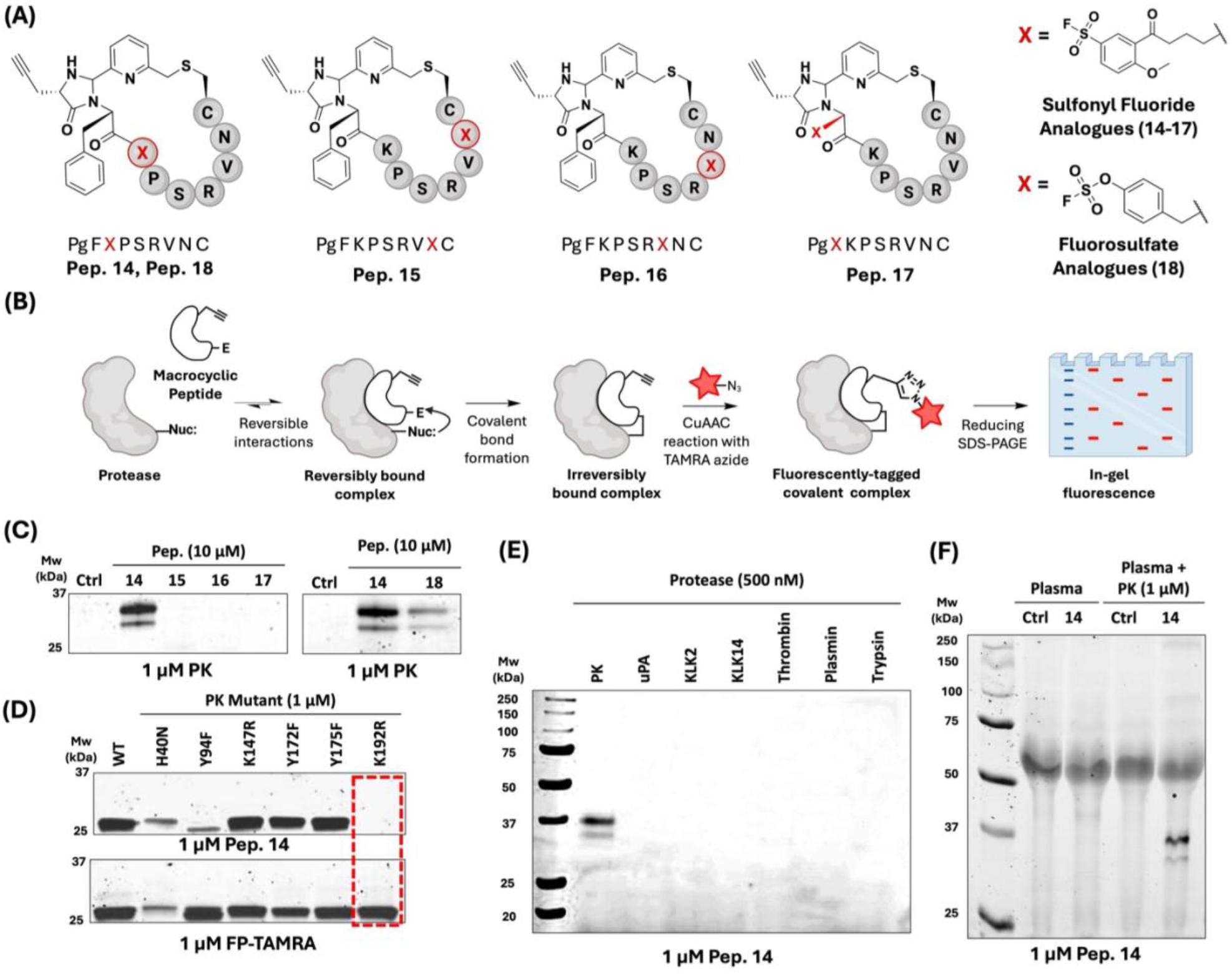
Development of a covalent macrocyclic peptide inhibitor of plasma kallikrein. (A) Structures of the SuFEx analogues of peptide **2** with either a sulfonyl fluoride (**14-17**) or fluorosulfate (**18**) electrophile incorporated. (B) Schematic of the activity-based protein profiling and in-gel fluorescence workflow for visualisation of covalent bond formation. (C) In-gel fluorescence analysis of plasma kallikrein labelling by peptides **14-18**. (D) In-gel fluorescence analysis of plasma kallikrein mutant labelling by either peptide **14** or fluorophosphonate-TAMRA. (E) In-gel fluorescence analysis of serine protease labelling by peptide **14**. PK = plasma kallikrein, uPA = urokinase plasminogen activator, KLK = kallikrein. (F) In-gel fluorescence analysis of plasma kallikrein labelling in blood plasma by peptide **14**. Plasma was incubated with peptide **14** both with and without plasma kallikrein addition.

To generate the analogue series, linear peptides were synthesised by standard Fmoc SPPS with alloc-protected 2,4-diaminobutyric acid (Dab) incorporated at the desired position for electrophile installation (**Scheme S2**). Following selective Pd-catalysed alloc deprotection, 5-(fluorosulfonyl)-2-methylbenzoic acid was then coupled to the Dab side chain. The N-terminal alanine was replaced with propargylglycine to provide an alkyne handle for downstream activity-based protein profiling (ABPP) experiments (**Fig. 6B**).^40^ Following cleavage and global deprotection, the linear peptides were cyclised in-solution with the BMP-linker to afford the final covalent macrocycles. No attempt was made to separate diastereomers for any of the covalent analogues.

The sulfonyl fluoride analogues were assessed for covalent labelling of PK by ABPP and in-gel fluorescence. Following incubation of each macrocycle (10 µM) with PK (1 µM) for 1 h at 37 °C, covalent labelling was visualised by in-gel fluorescence after copper-catalysed azide–alkyne cycloaddition (CuAAC)-mediated conjugation of the incorporated alkyne tag to an azide TAMRA reporter (**Fig. 6B**). Of the analogues tested, only incorporation of the SuFEx electrophile at the Lys-3 position (peptide **14**) yielded a fluorescent band indicative of covalent target engagement (**Fig. 6C**). Two bands were consistently observed by in-gel fluorescence for blood-purified human PK, aligning with previously reported heterogeneous glycosylation of the protease.^41^ Covalent modification by peptide **14** was further confirmed by intact protein mass spectrometry, which revealed 73% modification of PK (5 µM) after incubation with peptide **14** (50 µM) for 4 h at 37 °C (**Fig. S41**). These results identified the Lys-3 position as a productive site for electrophile installation and established peptide 14 as a promising covalent analogue for further characterisation.

Aryl sulfonyl fluorides have been widely employed in the development of covalent inhibitors and chemical probes due to their ability to achieve selective target engagement.^42^ However, alternative SuFEx electrophiles with further reduced intrinsic reactivity have also attracted interest for covalent ligand development.^28^ We therefore investigated whether the covalent macrocycle could be further optimised through incorporation of less reactive aryl fluorosulfate in place of the sulfonyl fluoride warhead. To generate the fluorosulfate analogue, the Dab residue of peptide **14** was replaced with an orthogonally protected tyrosine. This enabled installation of the fluorosulfate electrophile using AISF ([4-(Acetylamino)phenyl]-imidodisulfuryl difluoride) (**Fig. 6A, Scheme S3**).^43, 44^ Following cleavage and cyclisation with the BMP linker, the resulting fluorosulfate analogue (peptide **18**) was evaluated by ABPP and in-gel fluorescence. Peptide **18** retained the ability to covalently label PK but exhibited lower labelling efficiency than peptide **14 (Fig. 6C)**. Consequently, subsequent studies focused on further characterisation of peptide **14**.

To identify the PK nucleophilic residue responsible for covalent bond formation, peptide **14** was evaluated against a panel of active-site mutants using the ABPP workflow described above. To ensure that any loss of labelling reflected disruption of covalent bond formation rather than loss of enzyme activity, the catalytic activity of each mutant (H40N, Y94F, K147R, Y172F, Y175F, and K192R), was first confirmed using the broad-spectrum serine protease probe fluorophosphonate-TAMRA (FP-TAMRA) (**Fig. 6D, bottom**). Of the mutants against which peptide **14** was tested, loss of covalent labelling was observed for only the K192 mutant (**Fig. 6D, top**), implicating Lys-192 as the site of covalent modification. These results demonstrate that covalent bond formation is driven by productive molecular recognition rather than indiscriminate SuFEx reactivity, with the electrophile selectively engaging the nucleophilic residue appropriately positioned within the peptide-protein complex.

To assess the selectivity of peptide **14**, covalent labelling was evaluated against a panel of serine proteases by ABPP and in-gel fluorescence following incubation with 500 nM of each protease for 1 h at 37 °C. The serine proteases tested were urokinase plasminogen activator (uPA), kallikrein (KLK) 2, KLK14, thrombin, plasmin and trypsin. Peptide **14** exhibited complete selectivity for PK, with no detectable labelling of any other serine protease tested (**Fig. 6E**). In contrast, the broad-spectrum serine protease probe FP-TAMRA labelled multiple proteases under the same conditions (**Fig. S42**), highlighting the high selectivity afforded by the BMP-derived macrocyclic scaffold.

Finally, we assessed the stability of peptide **14** and its suitability for application in biologically relevant samples. The peptide displayed a half-life of 28.4 h in PBS (pH 7.4), as determined by HPLC analysis. (**Fig. S43**). Consistent with this stability, peptide **14** retained the ability to covalently label PK in human plasma as observed by ABPP (**Fig. 6F**). Together, these results demonstrate that peptide **14** combines selective covalent target engagement with sufficient stability for activity-based profiling in complex biological environments.

## Conclusion

In conclusion, we have developed a new strategy for cyclising phage-displayed peptides through reaction of a BMP linker with a cysteine side chain and the peptide N-terminus. This chemistry generates a previously unexplored macrocyclic topology incorporating neighbouring pyridine and imidazolidinone rings while simultaneously masking the peptide N-terminus. The rapid reaction kinetics and compatibility with phage display make the BMP linker a valuable addition to the repertoire of cyclisation strategies available for macrocyclic peptide discovery.

Proof-of-concept screening of a BMP-cyclised phage library against plasma kallikrein identified a potent macrocyclic inhibitor and demonstrated the importance of macrocyclic topology in ligand discovery. The BMP-derived scaffold afforded substantially greater potency than analogous disulfide- and DBMB-cyclised peptides, while comparison with an equivalent DBMB-cyclised library revealed binding motifs not identified by conventional cysteine-cysteine cyclisation. Furthermore, the ability of the BMP linker to generate two diastereomeric macrocycles from a single peptide sequence provides an additional source of structural diversity, illustrated by the greater than 20-fold difference in potency observed between the isolated diastereomers of peptide **2**.

Finally, we demonstrate a straightforward strategy for converting phage-derived macrocyclic peptide hits into selective covalent inhibitors through positional SuFEx electrophile scanning. This workflow enabled the development of a selective covalent macrocyclic activity-based probe capable of reporting plasma kallikrein activity in human plasma. Together, these findings establish BMP-mediated cyclisation as a versatile platform for both macrocyclic peptide discovery and the development of covalent chemical probes.

## Acknowledgements

This work was supported by AdvanCell Pty Ltd. This work was supported by the Engineering and Physical Sciences Research Council (UKRI117) and Medical Research Council (UKRI2332).

## Author Contributions

Author contributions are assigned using Contributor Role Taxonomy (CRediT).

E.S: Investigation, Data Curation, Methodology, Visualisation, Formal Analysis, Writing – original draft, Writing – review and editing. M.B: Investigation, Data Curation, Methodology. J.W: Investigation, Data Curation, Methodology. C.W: Investigation, Data Curation, Methodology, Resources. M.LB Investigation, Formal Analysis, Methodology, Resources. S.L: Conceptualisation, Funding Acquisition, Supervision, Formal Analysis, Writing – original draft, Writing – review and editing.

