## Supplementary Information for "Expanding Macrocyclic Topology through Cysteine-to-N-Terminal Cyclisation Enables Covalent Peptide Inhibitor Discovery"

### 1  Materials and Methods

#### 1.1 Chemicals and Reagents

All chemicals and reagents were purchased from commercial suppliers and used without further purification. Rink Amide MBHA resin (100-200 mesh) was obtained from Merck. Natural Fmoc-amino acids were obtained from Fluorochem. Amicon 10 kDa centrifugal filters were obtained from Merck. Zeba desalting columns (7 kDa) were obtained from ThermoFisher Scientific. Purified human plasma kallikrein and purified biotinylated human plasma kallikrein was obtained from Innovative Research. Streptavidin- and neutravidin-coated magnetic beads were purchased from ThermoFisher Scientific. 2xYT media and agar were purchased from Merck. All remaining chemical reagents and biological products were obtained from Fluorochem, VWR, Merck, ThermoFisher Scientific, TCI chemicals or BLD Pharmatech.

#### 1.2 Equipment List

Synthetic reactions were monitored by TLC on silica gel plates (60 Å, F254 indicator) and visualised with UV light (254 nm). Automated peptide synthesis was performed on a Multipep 2 (CEM). Chromatography was performed on either a CombiFlash Rf200 (Teledyne), CombiFlash NextGen 300 (Teledyne ISCO), ACCQ Prep 150 (Teledyne ISCO) or ECS28P01 Compact Preparative System (ECOM). Peptide drying was performed using a Genevac EZ-2 series centrifugal evaporator or an Edwards Modulyo Freeze Dryer. High-resolution mass spectrometry data was collected using an Agilent 1260 Infinity II HPLC coupled to an Agilent 6545Q-TOF (ESI). Analytical low-resolution mass spectrometry was conducted using an Agilent 1260 Infinity II HPLC coupled to an Agilent InfinityLab LC/MSD iQ (ESI). NMR analysis for small molecule product confirmation was conducted on a 400 MHz Bruker Advance Neo or 500 MHz Agilent ProPulse instrument. NMR analysis for peptide characterisation was conducted on a on a Bruker Avance III HD spectrometer operating at 700 MHz equipped with a 1.7 mm TCI microcryocoil probe. NMR spectra were processed using MestReNova 14.2.0.

DNA amplification was performed using a SimpliAmp Thermal Cycler (ThermoFisher Scientific). Gel electrophoresis was performed using an mPAGE Mini Gel Tank (Millipore) and PowerPac™ Basic (BioRad). Fluorescent plate-based enzymatic assays were carried out using a VANTAstar Plate reader (BMG Labtech) in kinetic mode for 30 min. In-gel fluorescence-based assays were imaged on a BioRad ChemiDoc™ Imaging System. Cell lysis was performed using a sonopuls UW 100 ultrasonic transducer (Bandelin). Protein purification was performed using an AKTA start chromatography system (Cytiva).

### 2 Chemical Synthesis

#### 2.1 General Procedures

##### 2.1.1 Automated Peptide Synthesis:

All peptides were prepared by standard Fmoc solid phase peptide synthesis (SPPS) on a 0.1 mmol scale using an automated peptide synthesiser (CEM MultiPep 2). Rink amide resin was manually weighed into 12 mL syringes fitted with polypropylene filters and pre-swelled in DCM prior to synthesis. The automated method used repeated cycles of the following steps for each amino acid coupling:

**Fmoc deprotections** - incubation with 20% 4-methylpiperinde in DMF (1 x 5 min then 1 x 7 min).

**Amino acid double couplings** - incubation with amino acid (0.4 mmol, 4 equiv.), HBTU (0.5 mmol, 5 equiv.) and DIPEA (0.5 mmol, 5 equiv.) in DMF (30 min, RT, shaking at 350 rpm, repeated twice for each amino acid).

**Washing** - DMF (x7) after each deprotection and (x3) after each double coupling.

A final wash with ethanol was performed following deprotection of the last amino acid.

##### 2.1.2 Peptide Cleavage:

Global deprotection and cleavage of completed linear peptides was performed manually utilising the cleavage cocktail TFA:H_2_O:TIS:DODT:phenol (90:2.5:2.5:2.5:2.5, v/v/v/v/w, approx. 3 mL per 0.1 mmol synthesis). The reaction vessel was agitated for 2 hours, the cleavage solution collected and the peptides precipitated out by the addition of cold ether (approx. 15 mL). The peptide was pelleted by centrifugation at 4700 rpm for 10 mins at 4 °C and washed once more with cold ether (approx. 15 mL) before leaving overnight to dry. Peptides were then purified as outlined in **2.1.3** or, when sufficiently pure, lyophilised and used crude for cyclisation.

##### 2.1.3 Product Purification:

Flash chromatography was performed using a CombiFlash NextGen 300 (Teledyne ISCO), CombiFlash Rf 200 (Teledyne ISCO) or ECS28P01 Compact Preparative System (ECOM). Normal-phase purifications were carried out using RediSep Silver Silica columns (Teledyne) and reverse-phase purifications were carried out using RediSep Gold C18 columns (Teledyne). For closely eluting peptides, semi-preparatory and preparatory scale purifications were carried out on either an ACCQ Prep 150 (Teledyne ISCO) or an ECS28P01 Compact Preparative System (ECOM), utilising either a ZORBAX Eclipse XDB-C18 column (Agilent, semi-preparatory) or RediSep Prep C18 column (Teledyne ISCO, preparatory). For all purifications, column size was selected dependent on scale of the purification. Flow rates were scaled appropriately based on column dimensions and following the manufacturer’s instructions. Fraction collection was guided by UV absorbance.

For all reverse-phase purifications, the mobile phase was Milli-Q H_2_O and HPLC-grade MeCN, both supplemented with 0.1% formic acid (v/v). A 5-95% MeCN gradient was used as standard, with the length and inclusion of isocratic holds dependent on the scale and difficulty of purification.

##### 2.1.4 Analytical LC-MS:

Low-resolution LC-MS analysis for reaction monitoring and cyclisation tracking was conducted on an Agilent 1260 Infinity II HPLC coupled to an Agilent InfinityLab LC/MSD iQ (ESI, Single Quadrupole). Separations were carried out using either a Poroshell 2.7 µm 120 Å ES-C18 column (50 x 2.1 mm) or a Jupiter 4 μm Proteo 90 Å, LC C18 column (150 x 4.6 mm). The mobile phase consisted of HPLC-grade H_2_O and HPLC-grade MeCN, both supplemented with 0.1% (v/v) formic acid. The flow rate was 1 mL/min and gradients extended from either 2-95% or 5-95% MeCN, with the length of gradient tailored to the specific reaction or product.

##### 2.1.5 High-Resolution Mass Spectrometry:

High-resolution mass spectrometry was conducted using an Agilent 1260 Infinity II HPLC coupled to an Agilent 6545Q-TOF mass spectrometer (ESI, Triple Quadrupole/Time-Of-Flight). Separations were carried out a Zorbax Eclipse Plus C18 Rapid Resolution HD (1.8µm, 2.1 x 50mm) for small molecules or a bioZenTM 2.6um peptide XB-C18 (50 x 2.1 mm) for peptides.

#### 2.2 6-(bromomethyl)picolinaldehyde Linker Synthesis

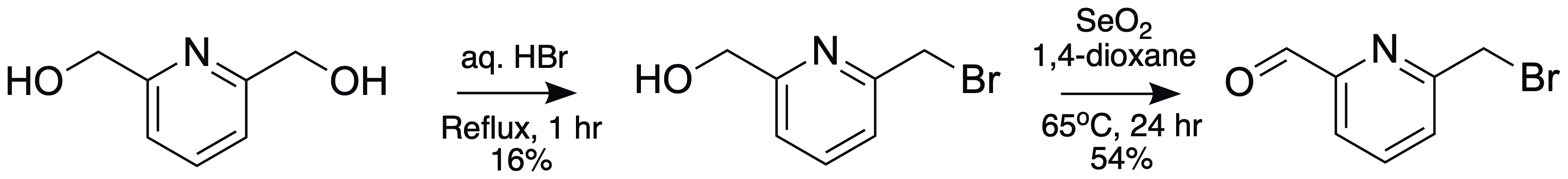

***Scheme 1: Synthetic strategy for BMP linker (6-(bromomethyl)picolinaldehyde).***

**(6-(bromomethyl)pyridin-2-yl)methanol (1):**

2,6-pyridinedimethanol (1.00 g, 1 equiv., 7.19 mmol) was added to 48% aq. HBr (10 mL, 12 equiv., 87.4 mmol) and the solution refluxed for 1 h. The reaction mixture was cooled to -15^o^C and neutralised by dropwise addition of aqueous NaOH, ensuring the temperature remained below -15 °C throughout. The solution was diluted with 10 mL H_2_O and the product extracted with 5x10 mL DCM. The organic fraction was dried with anhydrous MgSO_4_, concentrated under reduced pressure and the crude residue purified by flash chromatography using a 0-10% methanol/DCM gradient to yield **1** as a white solid (227 mg, 1.12 mmol, 16%). Synthesis guided by the literature.^1^

**LRMS** (ESI, positive mode) found 201.9, [C7H8BrNO+H]+ requires 201.9868.

**^1^H NMR** (400 MHz, CDCl_3_) δ 7.69 (t, J = 7.7 Hz, 1H), 7.35 (d, J = 7.7 Hz, 1H), 7.17 (d, J = 7.8 Hz, 1H), 4.76 (s, 2H), 4.55 (s, 2H).

**6-(bromomethyl)Picolinaldehyde (2):**

Selenium dioxide (62.1 mg, 0.5 equiv., 559 μmol) was added to a solution of compound **1** (226 mg, 1 equiv., 1.12 mmol) in 1,4-dioxane (4.5 mL). The reaction mixture was sonicated for 2 mins before stirring for 24 h at 65 °C. The reaction mixture was then cooled to RT, diluted with DCM (10 mL) and filtered through Celite. The filtrate was concentrated under reduced pressure and the crude residue purified by flash silica chromatography using a 0-10% methanol/DCM gradient to yield **2** as a light brown solid (224 mg, 607 µmol, 54%). Synthesis guided by the literature.^2^

**HRMS** (ESI, positive mode) found 199.9695, [C7H6BrNO+H]+ requires 199.9711.

**^1^H NMR** (400 MHz, CDCl_3_) δ 10.06 (s, 1H), 7.94 – 7.85 (m, 2H), 7.69 (dd, J = 5.4, 3.5 Hz, 1H), 4.63 (d, J = 0.7 Hz, 2H).

**^13^C NMR** (101 MHz, CDCl_3_) δ 193.14, 157.85, 152.64, 138.35, 127.91, 121.01, 32.91.

#### 2.3 6-(bromomethyl)picolinaldehyde Cyclisation Reactions

##### 2.3.1 Cyclisation Tracking on Synthetic Peptides

Lyophilised linear peptide (1 mM final concentration in a 1.2 mL reaction) was dissolved in the minimal volume of PBS (50 mM, pH 7.8)/MeCN required to achieve complete solubilisation. To reduce any intermolecular disulfide bonds prior to cyclisation, TCEP immobilised on agarose resin (0.1 equiv.) was then added and the solution incubated for 30 min at RT with continuous rotation. The resin was removed by centrifugation at 18,000 rpm and the supernatant collected. To the solution was then added 6-(bromomethyl)picolinaldehyde (1.2 mM final concentration) and the reaction volume adjusted to 1.2 mL with PBS (50 mM, pH 7.8) containing 20% (v/v) MeCN. The reaction mixture was incubated at 37 °C with continuous rotation and aliquots (50 µL) were taken after 30 min,1 h, 2 h, 3 h and 16 h. Samples were analysed by low-resolution LC-MS as outlined in **2.1.4**. The extent of cyclisation was determined by integrating the area under the curve of the corresponding peaks in the UV signal at 280 nm. Percentage of cyclised peptide was calculated relative to the total peptide signal (cyclised and linear alkylated peptide species combined).

##### 2.3.2 General Cyclisation Protocol for Scaled-Up Synthesis

Lyophilised linear peptide (1 equiv.) and 6-(bromomethyl)picolinaldehyde (1.2 equiv.) were dissolved at a peptide concentration of 10 mg/mL in PBS (50 mM, pH 7.8) containing 20% (v/v) MeCN. The reaction mixture was incubated for 2 h at 37 °C with continuous rotation, after which the cyclised peptide was purified by reverse-phase phase chromatography as described in section **2.1.3**.

##### 2.3.3 Investigating Imidazolidinone Reversion to Aldehyde

Peptides (AYYWQHTC) and (ALYWQHTC) were cyclised and purified as described in section **2.3.2.** Purified peptides were dissolved in PBS (10 mM, pH 7.4) at a concentration of 0.8 mg/mL and divided into three samples. One sample of each peptide was incubated at RT, the second at 37 °C and the third sample was pH adjusted to 1.5 (addition of 1% formic acid) before incubation at RT. Aliquots of each peptide under the three conditions were taken at 0 h, 1 h, 2 h, 3 h and 24 h and analysed by LC-MS as described in section **2.1.4**. Stability was assessed by monitoring the emergence of linear alkylated peptide overtime.

##### 2.3.4 Investigating Imidazolidinone Reversion to Imine

Peptide ALYWQHTC was cyclised and purified as outlined in section **2.3.2** and dissolved in DMSO (10 mM). The DMSO stock was diluted to a concentration of 1.1 mM with either PBS (10 mM, pH 7.5)^2^ or NaOAc (100 mM, pH 4.6)^3^. A solution of NaCNBH_3_ in H_2_O (30 µL, 250 mM, 25 equiv.) was added to each peptide solution (270 µL, final conc. 1 mM) and the reactions incubated at RT. Samples were taken at 30 min, 2 h and 24 h for both reaction conditions and run directly on the LC-MS with the extended gradient given in **Supplementary Table 1** to ensure separation of the cyclised peptide and reduced imine intermediate.

To confirm reduction conditions utilised were suitable for capturing imine formed, a crude cyclisation reaction (1 mM peptide ALYWQHTC, 1.2 mM BMP-linker, 80 % 10 mM PBS, pH 7.5, 20% MeCN) was incubated for 5 min at 30 °C after which LC-MS analysis confirmed alkylation of the peptide but minimal cyclisation. A solution of NaCNBH_3_ in H_2_O (30 µL, 250 mM, 28 equiv.) was then added to the reaction mixture (270 µL, final conc. 0.9 mM) which was incubated for a further 24 h at RT. Samples were taken at 30 min, 2 h and 24 h and run directly on the LC-MS with the extended gradient given in **Supplementary Table 1.**

| **Time (min)** | **Solvent A (%)**  **(H_2_O + 0.1% FA)** | **Solvent B (%)**  **(MeCN + 0.1% FA)** |
| --- | --- | --- |
| 0.00 | 95 | 5 |
| 10.00 | 95 | 5 |
| 35.00 | 70 | 30 |
| 40.00 | 5 | 95 |
| 45.00 | 5 | 95 |
| 50.00 | 95 | 5 |
| 60.00 | 95 | 5 |

***Supplementary Table 1: Timetable for analytical LC-MS of investigating imidazolidinone reversion to imine intermediate.***

#### 2.4 Peptide NMR Sample Preparation

Peptides ALYWQHTC and peptide **2a** (AFKPSRVNC) were cyclised and purified as outlined in section **2.3.2**. Purified samples were dissolved in d_6_-DMSO (10 mM) and spectra acquired at 298K on a Bruker Avance III HD spectrometer operating at 700 MHz equipped with a 1.7 mm TCI microcryocoil probe. Spectra were processed using MestReNova 14.2.0. ^1^H, HSQC, HSQC-TOCSY, HMBC, TOCSY, NOESY, ROESY, COSY and ^15^N HSQC spectra were acquired and utilised for ^1^H characterisation of both samples.

#### 2.5 Peptide 2a Isomer Characterisation by NMR

A conformer search was performed for both R and S diastereomers of the BMP-cyclised peptide **2** (AFKPSRVNC) using Rowan Scientific.^4^ Searching was performed using the ‘openconf – torsional Monte Carlo’ conformer generator^5^, conformer optimisation was performed using GFN2-xTB/ALPB(DMSO)^6, 7^ and final single point energies for each conformer calculated using GFN2-xTB/CPCM-X(DMSO)^6, 8^. Any conformers within < 3 kcal/mol of the lowest energy conformer were examined (3 conformers for R isomer, 2 conformers for S isomer). Interproton distances were calculated from the NOESY spectra of peptide **2a** and are more consistent with the R conformers. Specifically, the calculated interproton distances for the imidazolidinone proton (supplementary fig 31, no. 80) and the β-protons of the N-terminal alanine (**Fig. S26**, no. 78) aligned more closely to the R conformers over the S and no NOE was observed for the imidazolidinone proton and the α-protons of the N-terminal alanine, as would be expected for the S conformers. The R conformers also differ more significantly from each other than the S conformers, which is consistent with the NMR data within which some NH resonances are broadened or missing. Low energy conformers of the R and S isomers of cyclised peptide **2** are available from the corresponding author upon request.

#### 2.6 Peptide 2 Isomer Characterisation with NaCNBH_3_ Reduction

Peptides **2a** and **2b** (AFKPSRVNC) were cyclised and purified as outlined in section **2.3.2** and dissolved in DMSO (10 mM). The DMSO stock was diluted to a concentration of 1.1 mM with PBS (10 mM, pH 7.5). A solution of NaCNBH_3_ in H_2_O (30 µL, 500 mM, 25 equiv.) was added to each peptide solution (270 µL, final conc. 1 mM) and the reactions incubated at RT for 30 min before analysing by high-resolution mass spectrometry.

#### 2.7 Synthesis of Covalent Macrocyclic Peptides

##### 2.7.1 Sulfonyl Fluoride Peptides (14-17)

**Installation of 5-(Fluorosulfonyl)-2-methoxybenzoic acid covalent warhead**

Linear peptides **14-17** were synthesised by Fmoc SPPS as per section **2.1.1**, with Fmoc-Dab(alloc)-OH substituted into each position for electrophile incorporation and the N-terminal amino acid substituted for Boc-propargyl-OH. To selectively deprotect the Dab alloc protecting group, triphenylphosphine (64.8 μL, 3 equiv., 293 µmol) was dissolved in THF (3 mL). The solution was placed on ice and formic acid (37.3 μL, 10 equiv., 975 μmol) and diethylamine (101 μL, 10 equiv., 975 μmol) added. Still on ice, the solution was added to tetrakis(triphenylphosphine)palladium(o) (113 mg, 1 equiv., 97.5 μmol) and mixed before adding to the resin which was then agitated in the dark overnight at RT. The resin was washed with THF (2 x 3 mL), DMF (2 x 3 mL), DCM (2 x 3 mL) and 0.5% DIPEA in DMF (v/v, 1 x 3 mL) then 0.5% DIPEA in DMF (3 mL) was added and left for 10 min. Following this, a solution of 0.5% diethyldithiocabamate in DMF (w/v, 3 mL) was added to the resin which was then agitated for 30 min at RT before washing with DMF (3 x 3mL) and DCM (3 x 3 mL). The resin was then treated with a solution of 5-(fluorosulfonyl)-2-methoxybenzoic acid (34.3 mg, 1.5 equiv., 146 μmol), DIPEA (51.0 μL, 3 equiv., 293 μmol) and PyBOP (152 mg, 3 equiv., 293 μmol) in DMF (3 mL) and agitated for 1 h at RT. The resin was then washed with DMF (3 x 3 mL), DCM (3 x 3 mL) and dried under vacuum. The linear peptide was cleaved and cyclised as outlined in sections **2.1.2** and **2.3.2**.

***Scheme 2: Synthetic strategy for sulfonyl fluoride containing peptides (14-17).*** *a) Pd(PPh_3_)_4_ (1 equiv.), PPh_3_ (3 equiv.), NHEt_2_ (10 equiv.), HCOOH (10 Equiv.), THF (3 mL), 16 h, RT. b) 5-(fluorosulfonyl)-2-methoxybenzoic acid (1.5 equiv.), DIPEA (3 equiv.), PyBOP (3 equiv.), DMF, 1 h, RT. c) TFA (90%), TIS (2.5%), H_2_O (2.5%), DODT (2.5%), Phenol (2.5%), 2 h, RT. d) 6-(bromomethyl)picolinaldehyde (1.2 equiv.), PBS (50 mM, pH 7.8, 80%), MeCN (20%), 2 h, 37 ^o^C.*

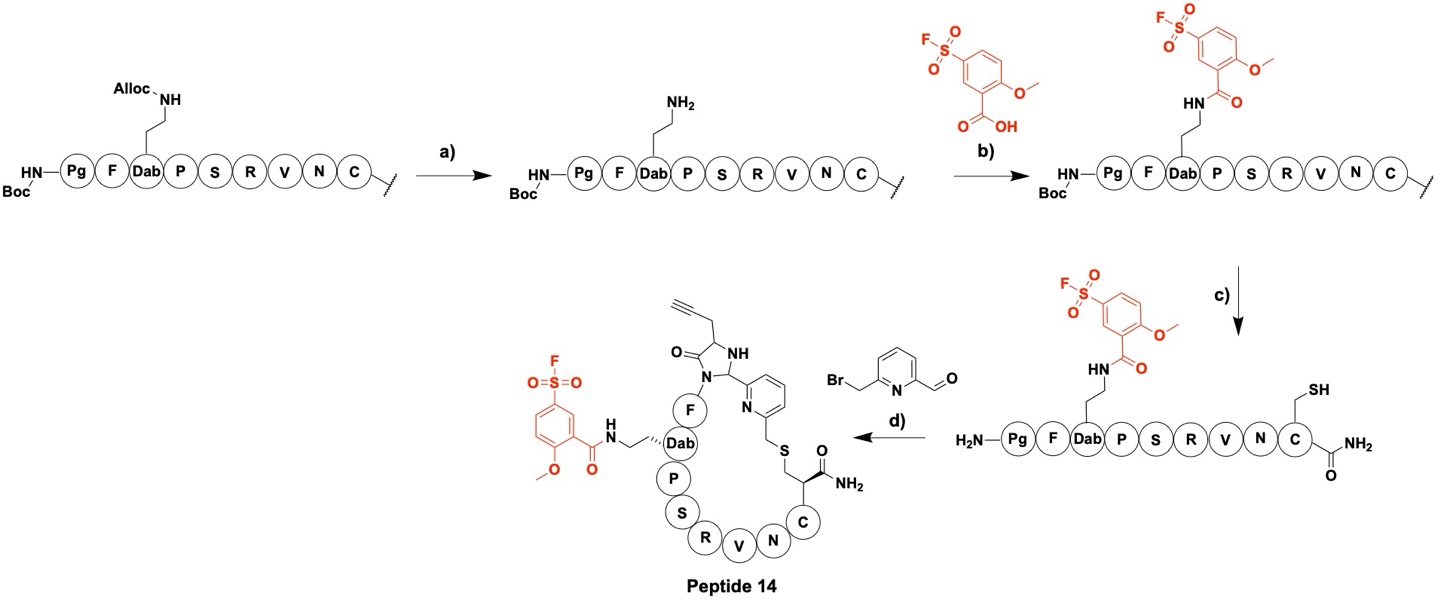

##### 2.7.2 Fluorosulfate Peptide (18)

Linear peptide **18** was synthesised by Fmoc SPPS as per section **2.1.1**, with Fmoc-Tyr(o-allyl)-OH substituted into position 3 and the N-terminal amino acid substituted for Boc-propargyl-OH. To selectively deprotect the o-allyl group of the tyrosine, triphenylphosphine (64.8 μL, 3 equiv., 293 µmol) was dissolved in THF (3 mL). The solution was placed on ice and formic acid (37.3 μL, 10 equiv., 975 μmol) and diethylamine (101 μL, 10 equiv., 975 μmol) added. Still on ice, the solution was added to tetrakis(triphenylphosphine)palladium(o) (113 mg, 1 equiv., 97.5 μmol) and mixed before adding to the resin which was then agitated in the dark overnight at RT. The resin was washed with THF (2 x 3 mL), DMF (2 x 3 mL), DCM (2 x 3 mL) and 0.5% DIPEA in DMF (v/v, 1 x 3 mL) then 0.5% DIPEA in DMF (3 mL) added and left for 10 min. A solution of 0.5% diethyldithiocabamate in DMF (w/v, 3 mL) was added to the resin which was then agitated for 30 min at RT before washing with DMF (3 x 3mL) and DCM (3 x 3 mL). A solution of 4-(Acetylamino)phenyl]imidodisulfuryl difluoride (AISF) (123 mg, 4 equiv, 390 µM) in THF (4 mL) was added to the resin followed by dropwise addition of 1,8-diazabicyclo[5.4.0]undec-7-ene (DBU) (117 µL, 8 equiv., 780 µM). The resin was agitated for 30 min at RT, washed with DCM and the AISF reaction repeated. The linear peptide was cleaved and cyclised as outlined in sections **2.1.2** and **2.3.2**.

***Scheme 3: Synthetic strategy for fluorosulfate containing peptide 18.*** *a) Pd(PPh_3_)_4_ (1 equiv.), PPh_3_ (3 equiv.), NHEt_2_ (10 equiv.), HCOOH (10 Equiv.), THF (3 mL), 16 h, RT. b) 4-(Acetylamino)phenyl]imidodisulfuryl difluoride (4 equiv.), 1,8-Diazabicyclo[5.4.0]undec-7-ene (8 equiv.), THF 30 min, RT. c) TFA (90%), TIS (2.5%), H2O (2.5%), DODT (2.5%), Phenol (2.5%), 2 h, RT. d) 6-(bromomethyl)picolinaldehyde (1.2 equiv.), PBS (50 mM, pH 7.8, 80%), MeCN (20%), 2 h, 37^o^C.*

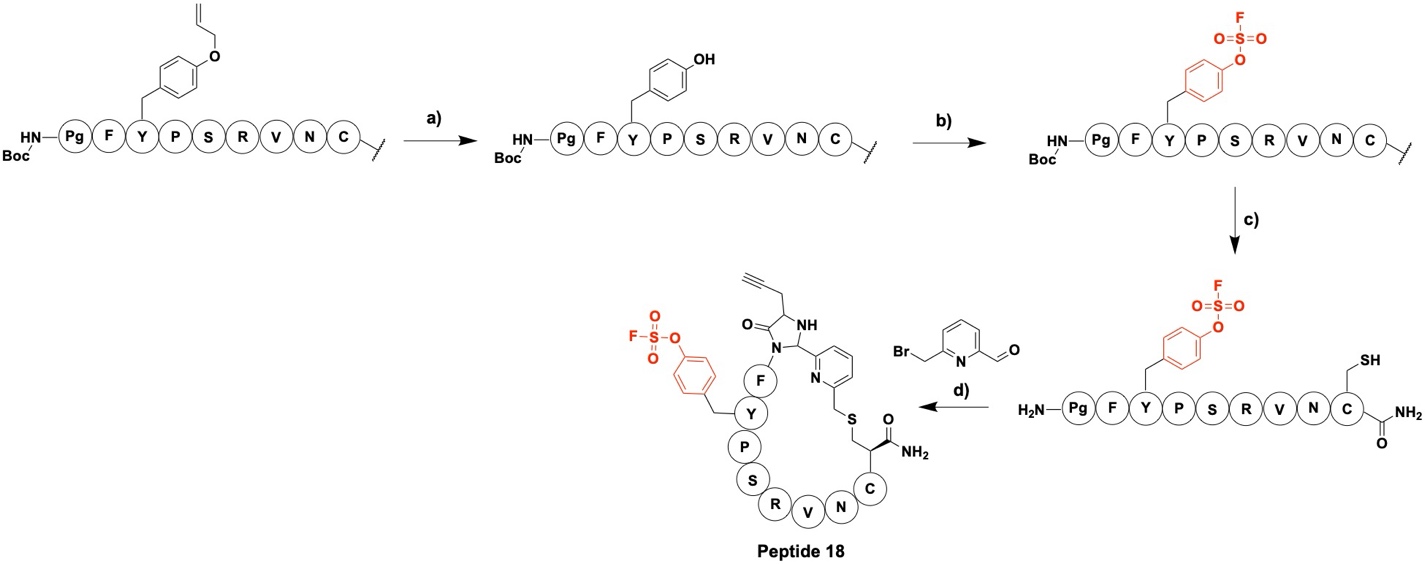

### 3 Biological and Biochemical Methods

#### 3.1 Peptide-pIII Fusion Protein Expression and Purification

##### 3.1.1 Expression Construct

A pET-28b(+) expression vector encoding the peptide-pIII protein was purchased from GenScript (CloneEZ method). The following sequence was incorporated to encode for the N1 and N2 domains of the disulfide-free phage pIII protein fused to the model peptide sequence AAYWQHTC, plus a C-terminal 6x His-tag for purification:

ATGGCGGCGTATTGGCAGCATACCTGTGGTGGCGGTAGCGAGGGTGGCTCTGGTGCTGAAACTGTTGAAAGTAGTTTAGCAAAATCCCATATAGAAGGTTCATTTACTAACGTCTGGAAAGACGACAAAACTTTAGATTGGTACGCTAACTATGAGGGCATCCTGTGGAAGGCTACAGGCGTTGTAGTTATAACTGGTGACGAAACTCAGGTATACGCTACATGGGTTCCTATTGGGCTTGCTATCCCTGAAAATGAGGGTGGTGGCTCTGAGGGTGGCGGTTCTGAGGGTGGCGGTTCTGAGGGTGGCGGTACTAAACCTCCTGAGTACGGTGATACACCTATTCCGGGCTATACCTATATCAACCCTCTCGACGGCACTTATCCGCCTGGTACTGAGCAAAACCCCGCTAATCCTAATCCTTCTCTTGAGGAGTCTCACCCTCTTAATACTTTCATGTTTCAGAATAATAGGTTCCGAAATAGGCAGGGGGCATTAACTGTTTATACGGGCACTGTTACTCAAGGCACTGACCCCGTTAAAACTTATTACCAGTACACTCCTGTATCATCAAAAGCCATGTATGACGCTTACTGGAACGGTAAATTCAGAGACGTGGCTTTCCATTCTGGCTTTAATGAGGATTTACTGGTTGCGGAATATCAAGGCCAATCGTCTTACCTGCCTCAACCTCCTGTCAATGCGGCCGCGGCCTCGGGGGCCATGGCTTCTGGTACCCCGGTTAACGCTCACCACCACCACCACCACTGA

##### 3.1.2 Expression and Purification

The expression construct (350 ng, 3.5 µL) was transformed into 100 µL electrocompetent BL21(DE3) *E. coli* cells (2.5 eV). The cells were recovered in 1 mL SOC outgrowth media (New England Biolabs) for 1 h at 37 °C with shaking (180 rpm) before plating on 2xYT agar plates supplemented with kanamycin (50 µg/mL). A single colony was then picked and grown for ~ 7 h at 37 °C and 180 rpm. The culture was centrifuged at 4000 rpm for 5 min at RT and the pellet resuspended in 50/50 (v/v) 2xYT/50% glycerol (400 µL), flash frozen and stored at -70 °C as a cell stock.

2xYT media (10 mL) supplemented with 50 µg/mL kanamycin was inoculated with the cell stock and grown at overnight at 37 °C with shaking (180 rpm). 5 mL of the overnight culture was then diluted (1:100) with 2xYT media supplemented with 50 µg/mL kanamycin and grown at 37 °C with shaking (180 rpm) until an OD_600_ of 0.6 was reached. To induce protein expression, IPTG (1 mM) was added and the culture grown overnight at 25 °C (180 rpm). The cells were collected by centrifugation at 8000 x g for 20 min at 4 °C and lysed by sonication (6 x 20 sec on/20 s off, 50% power) in 18 mL lysis buffer (30 mM PBS, pH7.4, 1 mM TCEP, 0.1 v/v Triton X-100, protease inhibitor cocktail (Merck)). The lysed cells were centrifuged at 8000 x g for 25 min at 4 °C and the supernatant collected. Imidazole (6.8 mg, 10 mM) was added to 10 mL of the cell lysate which was then purified on a 5 mL HisTrap HP column (Cytiva) at a flow rate of 5 mL/min. After protein loading, the column was first washed with 10 CV of buffer (30 mM PBS, pH 7.4, 10 mM imidazole, 1 mM TCEP) before eluting the protein by applying a linear gradient from 100-500 mM imidazole over 10 CV. Protein fractions were analysed by SDS-PAGE and pure fractions combined. The eluted protein was then further purified on a HiPrep 16/60 Sephacryl S-100 HP (Cytiva) using a flow rate of 0.5 mL and eluting with buffer (20 mM NH_4_HCO_3_, pH 8.0, 5 mM EDTA, 1 µM TCEP). Protein fractions were analysed by SDS-PAGE, pure fractions combined before concentrating using a protein concentrator (ThermoFisher Pierce) and storing at -70 °C.

#### 3.2 Cyclisation of Peptide on pIII Fusion Protein

The peptide-pIII construct was buffer exchanged into PBS (10 mM, pH 8) using a Zeba desalting column (ThermoFisher) and the concentration determined using the Bradford assay. The BMP-linker (1 mM, 50 equiv.) was added to the protein (20 µM) along with 20% MeCN (v/v) and incubated at 30 °C for 1 h. The protein was then buffer exchanged into fresh PBS (10 mM, pH 8), to remove excess linker and the MeCN, and incubated at 37 °C for a further 2 h. Samples were taken after the 1 h incubation at 30 °C and after an additional 1 h and 2 h incubation at 37 °C and analysed by high-resolution LC-MS on an Agilent 1260 Infinity II HPLC equipped with an PLRP-S (polymeric Reversed phase) 300 Å, 3.0 µm (2.1 x 50mm) column and coupled to an Agilent 6545Q-TOF mass spectrometer (ESI, Triple Quadrupole/Time-Of-Flight). Protein masses were assessed by biomolecular deconvolution using Agilent MassHunter BioConfirm Version 10.0.

#### 3.3 Phage Library Generation

The fd phage vector fdg3p0ss, obtained from a library of phage displaying an XCX_7_C peptide, was used as the template for a whole-plasmid PCR.^9-11^ The vector encodes for an fd phage with the pIII protein mutated to remove its three disulfide bonds. The DNA encoding the displayed peptide is incorporated between the pelB leader sequence and the N1 domain of the pIII protein, separated by a glycine-rich linker. The vector contains a chloramphenicol resistance and a single NgoMIV cleavage site. Incorporation of a second NgoMIV cleavage site during whole-plasmid PCR amplification allows for digestion and self-ligation of the library plasmid.

##### 3.3.1 Whole-plasmid PCR

Forward primers were designed to incorporate degenerate bases encoding the AX_7_C display peptide sequences plus the second NgoMIV cleavage site. A single reverse primer was designed to be used for both library generations. Primers were ordered from Merck and are given in **Supplementary Table 2**. Reagents and thermocycler conditions utilised are given in **Supplementary Table 3** and **4.** The PCR reaction was purified with a clean-up kit prior to digestion (GeneJET PCR Purification Kit, ThermoScientific).

| Primer | Fwd/Rv | Sequence (5’ 🡪 3’) | Encoded Peptide |
| --- | --- | --- | --- |
| 1 |  | AGCGCCGGCCATGGCAGCACGTNNKNNKNNK  NNKNNKNNKTGTGGTGGCGGTAGCGAGGGTG | ARX_6_C |
| 2 |  | AGCGCCGGCCATGGCAGCANNKCGTNNKNNK  NNKNNKNNKTGTGGTGGCGGTAGCGAGGGTG | AXRX_5_C |
| 3 |  | AGCGCCGGCCATGGCAGCANNKNNKCGTNNK  NNKNNKNNKTGTGGTGGCGGTAGCGAGGGTG | AX_2_RX_4_C |
| 4 |  | AGCGCCGGCCATGGCAGCANNKNNKNNKCGTN  NKNNKNNKTGTGGTGGCGGTAGCGAGGGTG | AX_3_RX_3_C |
| 5 |  | AGCGCCGGCCATGGCAGCANNKNNKNNKNNK  CGTNNKNNKTGTGGTGGCGGTAGCGAGGGTG | AX_4_RX_2_C |
| 6 |  | AGCGCCGGCCATGGCAGCANNKNNKNNKNNK  NNKCGTNNKTGTGGTGGCGGTAGCGAGGGTG | AX_5_RXC |
| 7 |  | AGCGCCGGCCATGGCAGCANNKNNKNNKNN  KNNKNNKCGTTGTGGTGGCGGTAGCGAGGGTG | AX_6_RC |
| 8 | Rv | CCAGAGCCACCCTCGCTACCG |  |

***Supplementary Table 2: Primers used for the whole-plasmid PCR generation of the AX_7_C library vector.***

| **Reagent** | **Quantity** |
| --- | --- |
| Fwd primer(s) | 200 nM |
| Rv primer | 200 nM |
| dNTP mix (New England Biolabs) | 200 µM |
| Template DNA | 20 ng |
| 5X Phusion HF buffer (ThermoScientific) | 10 µL |
| HF Phusion DNA Polymerase (ThermoScientific) | 2 µL (4U) |
| Nuclease free water | Made up to 50 µL |

***Supplementary Table 3: Reagents used for the whole-plasmid PCR generation of the AX_7_C library vector.***

| **Temperature (°C)** | **Time** | **Cycles** |
| --- | --- | --- |
| 95 | 5 min | 1 |
| 95 | 30 sec | 25 |
| 71 | 25 sec |  |
| 72 | 3 min 30 s |  |
| 72 | 7 min | 1 |
| 4 | $\infty$ | - |

***Supplementary Table 4: Thermocycler conditions used for the whole-plasmid PCR generation of the AX_7_C library vector.***

##### 3.3.2 Digestion

The amplified DNA was digested in 50 µL reaction volumes and incubated for 2 h at 37 °C. Reagents were as follows: Amplified DNA (1 µg), NgoMIV (1 µL,1U) (New England Biolabs),10X rCutSmart buffer, made up to 50 µL with nuclease free water. The digested DNA was then purified prior to ligation (GeneJET PCR Purification Kit, ThermoScientific).

##### 3.3.3 Ligation

The digested DNA was ligated in 50 µL reaction volumes and incubated overnight at 16 °C. Reagents were as follows: Digested DNA (2.5 µg), T4 DNA ligase (5 µL, 25 U) (ThermoScientific), 10X T4 DNA ligase buffer, made up to 50 µL with nuclease free water. The ligated DNA was then purified (Monarch Spin PCR & DNA Cleanup Kit, New England Biolabs).

##### 3.3.4 Transformation

Ligated DNA (1 µg) was transformed into freshly made electrocompetent TG1 *E. coli* cells (100 µL cells) at 2.5 eV. The cells were recovered in 1 mL pre-warmed SOC outgrowth media (New England Biolabs) for 1 h at 37 °C with shaking (180 rpm). 10 µL of cells were then serially diluted with 2xYT media and 10 µL of each dilution spotted in triplicate on small (9 cm) 2xYT agar plates supplemented with 25 µg/mL chloramphenicol. The remaining cells were spread over 5 large (14 cm) 2xYT agar plates supplemented with 25 µg/mL chloramphenicol. Plates were grown overnight at 37 °C and the cells from the large plates collected with 50/50 (v/v) 2xYT and 50% glycerol before storing at -70 °C.

The visible colonies on the small agar plate were counted and the following formula used to calculate the number of cells transformed with a library plasmid (i.e. the theoretical diversity generated):

$$\frac{Phage infected cells}{mL}= \frac{x}{10} \times{10}^{y}\times{10}^{3}$$

***Supplementary Formula 1: Formula used to calculate the number of cells infected with a phage plasmid per mL.***

Where $x$ = the average number of colonies across the three replicates for each dilution and $y$ = the dilution factor yielding the countable colonies.

Transformations were repeated until sufficient theoretical library diversity was obtained. All cell stocks were combined and re-aliquoted into 1 mL aliquots once the transformation process was complete.

#### 3.4 Phage Production

An aliquot (1 mL) of the desired phage library in TG1 *E.coli* was thawed on ice and used to inoculate 500 mL of 2xYT media supplemented with 25 µg/mL chloramphenicol. The culture was grown for 16 h at 30 °C with shaking (180 rpm) before centrifugation at 8500 rpm for 30 min at 4 °C. The supernatant was collected and the cell pellet discarded.

To precipitate out the phage, 25% (v/v) precipitation buffer (20% w/v PEG 8000, 2.5 M NaCl) was added to the culture supernatant. The mixture was incubated for 60 min at 4 °C with gentle swirling every 15 min to maximise phage recovery. Precipitated phage were then pelleted by centrifugation at 9000 rpm for 60 min at 4 °C. The supernatant was discarded and the inner walls of the tube carefully dried to maximise PEG removal without disturbing the phage pellet. The supernatant was re-suspended in sterile PBS (10 mL, 10 mM) and transferred to a fresh tube containing 25% (v/v) of the PEG precipitation solution. This was incubated for 60 min at 4 °C before again pelleting the phage by centrifugation at 9000 rpm for 60 min at 4 °C. The supernatant was discarded, and the final pellet was resuspended in 1mL PBS with 25% glycerol (v/v). A final purification step was performed by centrifugation at 18000 rpm for 20 min at 4 °C to remove cell debris. The purified phage stock was stored at -20 °C until further use.

To monitor phage production, serial 10-fold dilutions of the phage stock were prepared in sterile PBS. A culture of TG1 *E.coli* cells, grown overnight at 37 °C from a single colony, was diluted 1:100 with fresh 2xYT media and grown at 37 °C until exponential growth phase (OD_600_ = 0.4) was reached. Separate180 μL aliquots of the exponential cells were infected with 20 µL of suitable phage dilutions (e.g., 10^6^, 10^7^, 10^8^, 10^9^). The infection mixtures were incubated for 90 min at 37 °C (without shaking) before spotting in triplicate 10 µL of each dilution on a small 2xYT agar plate supplemented with 25 µg/mL chloramphenicol. The plate was incubated overnight at 37 °C and the next day visible colonies counted. The number of phage particles per mL (assumed to be equal to the number of cells infected per mL) was determined using the **Supplementary Formula 1**.

#### 3.5 On-Phage Peptide Cyclisation

Any intermolecular disulfide bonds between displayed peptides were first reduced by incubating the collected phage (1 mL, in 25% v/v glycerol in PBS) with TCEP (2 mM) for 30 min at RT. The reduced phage were then precipitated out with the PEG/NaCl buffer, as described above, but with 30 min incubation and centrifugation times due to the smaller working volume. The pelleted phage were resuspended in an alkylation buffer (800 µL, 50 mM NH_4_HCO_3_, 5 mM EDTA, pH 8.5) and the BMP-linker added (40 µM) in MeCN (200 µL) to give a final reaction volume of 1 mL with 20% MeCN (v/v). The reaction mixture was incubated with rotation for 60 min at 30 °C. To remove excess linker and MeCN, the phage were precipitated as described above and resuspended in sterile PBS (500 µL,10 mM, pH 8) before incubating with rotation for a further 60 min at 37 °C. A sterile 50% glycerol solution (500 µL) was then added to give a final volume of 1 mL (25% v/v glycerol total) and the stock stored in at -20 °C until further use.

#### 3.6 Toxicity Titres

Samples of phage were set aside after reduction, alkylation for 60 min at 30 °C and cyclisation for a further 60 min at 37 °C. Serial dilutions were performed in sterile PBS and phage numbers monitored as described in section **3.4**.

#### 3.7 Phage Panning

Three rounds of panning were performed with the AX_7_C library against biotinylated plasma kallikrein (PK) (Innovative Research) immobilised onto either streptavidin- or neutravidin-coated magnetic beads (Cytiva). To prevent the enrichment of matrix-specific background binders, the capture matrix was alternated between rounds (i.e., streptavidin-coated beads were used for the first and third rounds and neutravidin-coated beads for the second round). Each panning round included a parallel negative control; wherein half of the input phage was panned against target-free magnetic beads to monitor for non-specific binding.

The day prior to biopanning, a 5 mL 2xYT culture was inoculated with a single colony of TG1 *E. coli*. On the day of panning, 2 mL of the overnight culture was diluted 1:100 with fresh 2xYT media and grown at 37 °C until exponential growth phase (OD_600_ = 0.4) was reached. These cells were used for phage titres and infecting with the output phage.

The cyclised AX_7_C phage library was precipitated using the PEG/NaCl protocol described above and resuspended in PK activity buffer (10 mM Tris–base, pH 7.4, 150 mM NaCl, 10 mM MgCl_2_, 1 mM CaCl_2_) supplemented with 0.1% (v/v) Tween-20 and 1% (w/v) BSA. 10 µL of the resuspended phage solution was set aside for titres to monitor the input number of phage. The phage were allowed to block by rotating for 30 min at RT.

Prior to target incubation, the cyclised phage library was subjected to a pre-clearing step to deplete non-specific bead binders. A 50 μL aliquot of a 1:1 mixture of streptavidin and neutravidin magnetic beads was washed three times with 200 μL PK activity buffer. The beads were then blocked by incubating in 450 µL PK activity buffer supplemented with 0.1% (v/v) Tween-20 and 1% (w/v) BSA for 30 mins at RT, with continuous rotation. The blocking solution was then removed and the beads re-suspended with the blocked phage library. This mixture was incubated for 60 min at RT, with continuous rotation.

Whilst the phage were pre-clearing, biotinylated PK was immobilised onto the streptavidin/neutravidin beads (dependent on the panning round). To do so, 100 µL of beads were washed three times with 200 µL PK activity buffer. They were then resuspended in 100 µL PK activity buffer and divided equally between two eppendorfs. Plasma kallikrein (5 µg) was added to one and equal volume of PK activity buffer to the other (to act as a negative control). Both bead solutions were rotated for 10 min at RT. The beads were then captured, washed three times with 1 mL PK activity buffer and blocked by incubating for 30 min at RT in 450 µL PK activity buffer supplemented with 0.1% (v/v) Tween-20 and 1% (w/v) BSA.

Following pre-clearing of the phage, the beads were captured and the supernatant containing unbound phage collected. The phage supernatant was then split equally: half was added to the beads with PK immobilised, and the other half was added to the beads without PK (the negative control). Both samples were then incubated at 37 °C for 30 min with continuous slow rotation.

After target incubation, the beads were captured and washed first with PK activity buffer supplemented with 0.1% (v/v) Tween-20 then with just PK activity buffer. To increase the selection stringency, the duration and number of washes was increased across successive panning rounds (see **Supplementary Table 5**). To minimise the retention of plastic-binding phage, the samples were transferred to clean eppendorfs three times across the course of the washes.

| **Panning Round** | **Washes** |
| --- | --- |
| 1 | 8 x 2 mins PK activity buffer + 0.1% (v/v) Tween-20  2 x 2 min PK activity buffer |
| 2 | 4 x 2 mins PK activity buffer + 0.1% (v/v) Tween-20  2 x 10 mins PK activity buffer + 0.1% (v/v) Tween-20  4 x 2 mins PK activity buffer + 0.1% (v/v) Tween-20  2 x 2 min PK activity buffer |
| 3 | 4 x 2 mins PK activity buffer + 0.1% (v/v) Tween-20  3 x 20 mins PK activity buffer + 0.1% (v/v) Tween-20  4 x 2 mins PK activity buffer + 0.1% (v/v) Tween-20  2 x 2 min PK activity buffer |

***Supplementary Table 5: Washing steps for successive phage panning rounds.***

Following removal of the final wash, the beads were resuspended in 100 µL elution buffer (50 mM glycine, pH 2.2) and incubated for 10 min at RT with rotation. The beads were then captured and the supernatant containing the eluted phage collected and transferred to eppendorfs containing 50 µL neutralisation buffer (1 M Tris–base, pH 8.0). 10 µL of both the PK and negative control samples was set aside and titres performed as outlined in **3.4** to monitor the phage count. The remaining phage were infected into separate 50 mL aliquots of the exponential TG1 *E. coli*. The cultures were incubated for 90 min at 37 °C without shaking before pelleting the cells by centrifugation at 4,000 rpm for 10 min at 4 °C. The cell pellets were resuspended in 1 mL of 2xYT media and plated onto two large (14 cm) 2xYT agar plates supplemented with 25 µg/mL chloramphenicol and incubated overnight at 37 °C. The following day, the amplified cells were harvested by scraping the plates with 2xYT media (2 mL per plate). To the harvested cells was added an equal volume of a 50% glycerol solution to give a final glycerol concentration of 25% (v/v). 1 mL aliquots were stored at -70 °C. To initiate the next round of panning, cyclised input phage were produced from these glycerol stocks following the procedures described in Sections **3.4** and **3.5**.

The enrichment factor for each panning round was determined by calculating the ratio of phage/mL recovered from the PK output against the phage/mL from the negative control output.

#### 3.8 Sample Preparation for Next-Generation Sequencing

Following the third round of selection, DNA from the phage enriched against PK and the negative control were prepared for Next-Generation Sequencing (NGS) analysis. All samples were prepared in triplicate. A two-step PCR amplification strategy was employed. The primary PCR utilised a conserved set of five forward and five reverse primers to amplify the peptide-encoding region and incorporate an annealing region for barcoding primers, plus an ‘N’ spacer to improve sequencing quality. The secondary PCR step utilised unique combinations of forward and reverse primers to append sample-specific indices (barcodes) for multiplexed sequencing analysis. All NGS primers were ordered from Merck.

To isolate template DNA for the primary PCR, plasmid DNA was extracted from three 200 µL aliquots of the glycerol stocks of cells collected after the third round of selection against both PK and the negative control. This was done using the Monarch Plasmid Miniprep Kit (New England Biolabs) according to the manufacturer’s instructions. The primary PCR amplification was performed using the primers (Merck), reagents and thermocycler conditions given in **Supplementary Tables 6, 7 and 8**.

| **Round** | **Fwd/Rv** | **Primer** | **Sequence** |
| --- | --- | --- | --- |
| 1 | Fwd | 1a | GTCTCGTGGGCTCGGAGATGTGTATAAGAGACAGG**CTATGCGGCCCAGCCGGCC** |
|  |  | 1b | GTCTCGTGGGCTCGGAGATGTGTATAAGAGACAGGN**CTATGCGGCCCAGCCGGCC** |
|  |  | 1c | GTCTCGTGGGCTCGGAGATGTGTATAAGAGACAGGNN**CTATGCGGCCCAGCCGGCC** |
|  |  | 1d | GTCTCGTGGGCTCGGAGATGTGTATAAGAGACAGGNNN**CTATGCGGCCCAGCCGGCC** |
|  |  | 1e | GTCTCGTGGGCTCGGAGATGTGTATAAGAGACAGGNNNN**CTATGCGGCCCAGCCGGCC** |
|  | Rv | 2a | TCGTCGGCAGCGTCAGATGTGTATAAGAGACAG**CCAGAGCCACCCTCGCTACCG** |
|  |  | 2b | TCGTCGGCAGCGTCAGATGTGTATAAGAGACAGN**CCAGAGCCACCCTCGCTACCG** |
|  |  | 2c | TCGTCGGCAGCGTCAGATGTGTATAAGAGACAGNN**CCAGAGCCACCCTCGCTACCG** |
|  |  | 2d | TCGTCGGCAGCGTCAGATGTGTATAAGAGACAGNNN**CCAGAGCCACCCTCGCTACCG** |
|  |  | 2e | TCGTCGGCAGCGTCAGATGTGTATAAGAGACAGNNNN**CCAGAGCCACCCTCGCTACCG** |
| 2 | Fwd | S505 | AATGATACGGCGACCACCGAGATCTACACGTAAGGAG**TCGTCGGCAGCGTC** |
|  |  | S506 | AATGATACGGCGACCACCGAGATCTACACACTGCATA**TCGTCGGCAGCGTC** |
|  |  | S510 | AATGATACGGCGACCACCGAGATCTACACCGTCTAAT**TCGTCGGCAGCGTC** |
|  |  | S520 | AATGATACGGCGACCACCGAGATCTACACAAGGCTAT**TCGTCGGCAGCGTC** |
|  |  | S521 | AATGATACGGCGACCACCGAGATCTACACGAGCCTTA**TCGTCGGCAGCGTC** |
|  |  | S522 | AATGATACGGCGACCACCGAGATCTACACTTATGCGA**TCGTCGGCAGCGTC** |
|  | Rv | N715 | CAAGCAGAAGACGGCATACGAGATCCTGAGAT**GTCTCGTGGGCTCGG** |
|  |  | N716 | CAAGCAGAAGACGGCATACGAGATTAGCGAGT**GTCTCGTGGGCTCGG** |

***Supplementary Table 6: Primers used for two-step PCR amplification in Next-Generation Sequencing sample preparation.*** *Annealing sections are shown in bold and barcodes are shown in red.*

| **Reagent** | | **Quantity** |
| --- | --- | --- |
| Fwd primers: | 1a | 200 nM |
|  | 1b | 200 nM |
|  | 1c | 200 nM |
|  | 1d | 200 nM |
|  | 1e | 200 nM |
| Rv primers: | 2a | 200 nM |
|  | 2b | 200 nM |
|  | 2c | 200 nM |
|  | 2d | 200 nM |
|  | 2e | 200 nM |
| dNTP mix (New England Biolabs) | | 200 µM |
| Template DNA | | 200 ng |
| 5X Phusion HF buffer (ThermoScientific) | | 10 µL |
| HF Phusion DNA Polymerase (ThermoScientific) | | 2 µL (4U) |
| Nuclease free water | | Made up to 50 µL |

***Supplementary Table 7: Reagents used for primary PCR amplification in Next-Generation Sequencing sample preparation.***

| **Temperature (°C)** | **Time** | **Cycles** |
| --- | --- | --- |
| 95 | 5 min | 1 |
| 95 | 30 sec | 25 |
| 72 | 45 sec |  |
| 72 | 1 min |  |
| 72 | 1 min | 1 |
| 4 | $\infty$ | - |

***Supplementary Table 8: Thermocycler conditions used for primary PCR amplification in Next-Generation Sequencing sample preparation.***

The unpurified amplicon from the primary PCR was utilised directly as the template for the secondary PCR amplification. This was performed using the primers, reagents and thermocycler conditions given in **Supplementary Tables 6, 9 and 10**. Specific barcodes for each sample are given in **Supplementary Table 11.**

| **Reagent** | **Quantity** |
| --- | --- |
| Fwd primer | 200 nM |
| Rv primers | 200 nM |
| dNTP mix (New England Biolabs) | 200 µM |
| PCR 1 mixture | 2 µL |
| 5X Phusion HF buffer (ThermoScientific) | 10 µL |
| HF Phusion DNA Polymerase (ThermoScientific) | 2 µL (4U) |
| Nuclease free water | Made up to 50 µL |

***Supplementary Table 9: Reagents used for secondary PCR amplification in Next-Generation Sequencing sample preparation.***

| **Temperature (°C)** | **Time** | **Cycles** |
| --- | --- | --- |
| 95 | 5 min | 1 |
| 95 | 30 sec | 25 |
| 59 | 45 sec |  |
| 72 | 1 min |  |
| 72 | 7 min | 1 |
| 4 | $\infty$ | - |

***Supplementary Table 10: Thermocycler conditions used for secondary PCR amplification in Next-Generation Sequencing sample preparation.***

| **Sample** | **Fwd Primer** | **Rv Primer** |
| --- | --- | --- |
| PK 1 | S522 | N715 |
| PK 2 | S505 | N716 |
| PK 3 | S506 | N716 |
| Ctrl 1 | S510 | N715 |
| Ctrl 2 | S520 | N715 |
| Ctrl 3 | S510 | N715 |

***Supplementary Table 11: Barcodes used for each Next-Generation Sequencing sample.***

The secondary PCR products were resolved by electrophoresis on a 1% (w/v) agarose gel in Tris-Acetate-EDTA (TAE) buffer at 85 V for 65 min, utilising a 1 kb DNA Ladder (ThermoScientific). Amplicons corresponding to the expected size of 215-223 bp were excised from the gel and purified using the Monarch DNA Gel Extraction Kit (New England Biolabs) according to the manufacturer’s instructions. The intensity of each sample band was normalised against a standard to allow for equimolar amounts of the purified, barcoded amplicons to be pooled and sent together for NGS analysis (Source Bioscience).

#### 3.9 Next-Generation Sequencing Analysis

Forward and reverse reads from the raw sequencing data were merged using VSEARCH and reads failing marge criteria discarded to ensure high read quality. DNA sequences were then trimmed, translated and filtered to retain only peptide sequences of the library form (AX_7_C). No minimum count threshold was applied to the peptide sequences. Peptide counts from all samples were then arranged into a count matrix for downstream statistical analysis.

Statistical analysis was performed in R using the DESeq2 package. Peptide sequences were assessed for differential enrichment in the PK samples relative to the bead control samples using the default DESeq2 pipeline. Peptides were considered significantly enriched based on the following thresholds: $log2(Fold Change)$ > 1.5 and adjusted p-value $(p_{adj})$: < 0.05. Significantly enriched peptides were grouped into six clusters based on sequence similarity using the Gibbs clustering algorithm (GibbsCluster version 2.0). Clustered peptide motifs were visualised as sequence logos using WebLogo.

Custom scripts utilised for sequence filtering, quality control, and downstream data analysis are available from the corresponding author upon request.

#### 3.10 Enzyme Assays

The inhibitory activity of cyclic peptides was evaluated by incubating purified human plasma kallikrein (Innovative Research) (2.5 nM) with serially diluted peptide in a flat bottom 96 well plate (Corning) and quantifying residual enzyme activity using the fluorogenic substrate Z-Phe-Arg-7-amido-4-methylcoumarin hydrochloride (Merck). Peptide inhibitors were serially diluted in assay buffer (10 mM Tris–base, pH 7.4, 150 mM NaCl, 10 mM MgCl_2_, 1 mM CaCl_2_, 0.01% (w/v) BSA, 0.01% (v/v) Triton X-100, 5% (v/v) DMSO) and preincubated with plasma kallikrein at 37 °C for 15 min. The substrate was then added and enzyme activity determined by measuring fluorescence intensity at 1 min intervals over a 30 min period.

#### 3.11 Haemolysis Assay

Peptide **2a**, **12** and **13** were serially diluted in PBS in triplicate in a 96 well v-bottom plate (Corning) to a final volume of 50 µL. To each well was then added 50 µL of 5% (v/v) ice cold PBS (10 mM, pH 7.4) washed horse erythrocytes (ThermoFischer) before incubating the plate for 1 h at 37 °C. Following incubation, 100 µL of PBS (10 mM, pH 7.4) was added to each well following which the plate was centrifuged at 1000 x g for 15 min at RT. From each well, 100 µL of supernatant was then transferred to a flat bottom 96 well plate (Corning) and the amount of haemoglobin released measured by absorbance at 405 nm. Data was normalised to a positive control of 1% Triton X-100 only and a negative control of PBS only using the following formula:

$$\% Haemolysis=\left( \frac{absorbance of sample-absorbance of negative control}{absorbance of positive control-absorbance of negative control} \right)\times100$$

***Supplementary Formula 2: Formula used to calculate the percentage haemolysis for each peptide.***

Data was plotted and fitted in GraphPad Prism and is representative of 2 independent biological replicates.

#### 3.12 In-gel Fluorescence

The concentration of target protein and cyclic peptide probe was optimised for each in-gel fluorescence experiment. Control samples were treated with an equivalent volume of DMSO in place of the cyclic peptide probe. All primary incubations were performed in PBS (10 mM, pH 7.4) for 1 h at 37 °C and in a final reaction volume of 25 μL. Proteins were obtained from the following suppliers: Innovative Research (purified human plasma kallikrein), AcroBiosystems (urokinase-type plasminogen activator, recombinant human kallikrein 2, recombinant human kallikrein 14), R&D systems (human plasminogen, recombinant human coagulation factor II/thrombin), Promega (trypsin).

To perform the copper-catalysed azide-alkyne click (CuAAC) reaction, 2.5 µL of a freshly prepared ‘click’ chemistry cocktail was added to each target-probe reaction mixture and incubated for a further 60 min at RT. The ‘click’ chemistry cocktail consisted of 100 μM TAMRA-azide, 1 mM CuSO_4_, 1 mM TCEP and 100 μM TBTA (Tris((1-benzyl-4-triazolyl)methyl)amine). The CuAAC reactions were quenched with 4X SDS Laemmli Loading Buffer containg 5% β-mercaptoethanol (v/v) and denatured by heating at 95 °C for 10 min. Proteins were resolved via SDS-PAGE alongside a 10-250 kDa protein ladder (BioRad). In-gel TAMRA fluorescence was visualised using the Cy3 excitation/emission channels on a Gel Doc Imaging System (Bio-Rad).

#### 3.13 Intact Protein Mass Spectrometry

Intact mass spectrometry analysis of covalent labelling was performed using a non-glycosylated variant of the PK catalytic domain (I^371^-A^622^). Residues N^357^-R^371^ constitute part of the N-terminal heavy chain subunit of plasma kallikrein and are cleaved during expression to give active PK.^12^ This was purchased from GenScript, who expressed the protein in Sf9 cells followed by a HisTrap purification. To prevent heterogeneous glycosylation and eliminate unpaired thiols, the native sequence was engineered with the following substitutions: three N-linked glycosylation sites were mutated to glutamic acid (N377E, N434E, and N475E, highlighted in blue), and two specific cysteine residues were mutated to serine (C364S and C484S, highlighted in red).

N^357^TGDNSV**S**TTKTSTRI^372^VGGT**E**SSWGEWPWQVSLQVKLTAQRHLCGGSLIGHQWVLTAAHCFDGLPLQDVWRIYSGIL**E**LSDITKDTPFSQIKEIIIHQNYKVSEGNHDIALIKLQAPL**E**YTEFQKPISLP**S**KGDTSTIYTNCWVTGWGFSKEKGEIQNILQKVNIPLVTNEECQKRYQDYKITQRMVCAGYKEGGKDACKGDSGGPLVCKHNGMWRLVGITSWGEGCARREQPGVYTKVAEYMDWILEKTQSSDGKAQMQSPA^622^HHHHHH

To evaluate covalent-adduct formation, the engineered non-glycosylated PK (5 μM) was incubated with covalent probe (50 µM) in mass spectrometry assay buffer (10 mM Tris-HCl, 150 mM NaCl, pH 7.4). The reactions were incubated for 4 h at 37 °C with continuous rotation. The extent of covalent modification was analysed by high-resolution LC-MS on an Agilent 1260 Infinity II HPLC equipped with an PLRP-S (polymeric Reversed phase) 300 Å, 3.0 µm (2.1 x 50mm) column and coupled to an Agilent 6545Q-TOF mass spectrometer (ESI, Triple Quadrupole/Time-Of-Flight). Data processing was performed by biomolecular deconvolution using Agilent MassHunter BioConfirm Version 10.0.

#### 3.14 Expression of Plasma Kallikrein Mutants

Target nucleophilic residues for mutagenesis were selected based on spatial proximity to the active site of the PK catalytic domain, as determined from the published crystal structure (PDB: 2ANY). The corresponding mutant constructs were expressed and purified by GenScript using the TurboCHO platform and HisTrap columns, respectively. Mutant residues are highlighted in green. Mutant Y94F and H40N were activated by incubation with trypsin (1:100, w/w) for 1 h at 37 °C.

**H40N**

N^357^TGDNSVSTTKTSTRI^372^VGGTESSWGEWPWQVSLQVKLTAQR**N**LCGGSLIGHQWVLTAAHCFDGLPLQDVWRIYSGILELSDITKDTPFSQIKEIIIHQNYKVSEGNHDIALIKLQAPLEYTEFQKPISLPSKGDTSTIYTNCWVTGWGFSKEKGEIQNILQKVNIPLVTNEECQKRYQDYKITQRMVCAGYKEGGKDACKGDSGGPLVCKHNGMWRLVGITSWGEGCARREQPGVYTKVAEYMDWILEKTQSSDGKAQMQSPA^622^HHHHHH

**Y94F**

N^357^TGDNSVSTTKTSTRI^372^VGGTESSWGEWPWQVSLQVKLTAQRHLCGGSLIGHQWVLTAAHCFDGLPLQDVWRIYSGILELSDITKDTPFSQIKEIIIHQN**F**KVSEGNHDIALIKLQAPLEYTEFQKPISLPSKGDTSTIYTNCWVTGWGFSKEKGEIQNILQKVNIPLVTNEECQKRYQDYKITQRMVCAGYKEGGKDACKGDSGGPLVCKHNGMWRLVGITSWGEGCARREQPGVYTKVAEYMDWILEKTQSSDGKAQMQSPA^622^HHHHHH

**K147R**

N^357^TGDNSVSTTKTSTRI^372^VGGTESSWGEWPWQVSLQVKLTAQRHLCGGSLIGHQWVLTAAHCFDGLPLQDVWRIYSGILELSDITKDTPFSQIKEIIIHQEYKVSEGNHDIALIKLQAPLEYTEFQKPISLPSKGDTSTIYTNCWVTGWGFSKE**R**GEIQNILQKVNIPLVTNEECQKRYQDYKITQRMVCAGYKEGGKDACKGDSGGPLVCKHNGMWRLVGITSWGEGCARREQPGVYTKVAEYMDWILEKTQSSDGKAQMQSPA^622^HHHHHH

**Y172F**

N^357^TGDNSVSTTKTSTRI^372^VGGTESSWGEWPWQVSLQVKLTAQRHLCGGSLIGHQWVLTAAHCFDGLPLQDVWRIYSGILELSDITKDTPFSQIKEIIIHQEYKVSEGNHDIALIKLQAPLEYTEFQKPISLPSKGDTSTIYTNCWVTGWGFSKEKGEIQNILQKVNIPLVTNEECQKR**F**QDYKITQRMVCAGYKEGGKDACKGDSGGPLVCKHNGMWRLVGITSWGEGCARREQPGVYTKVAEYMDWILEKTQSSDGKAQMQSPA^622^HHHHHH

**Y175F**

N^357^TGDNSVSTTKTSTRI^372^VGGTESSWGEWPWQVSLQVKLTAQRHLCGGSLIGHQWVLTAAHCFDGLPLQDVWRIYSGILELSDITKDTPFSQIKEIIIHQEYKVSEGNHDIALIKLQAPLEYTEFQKPISLPSKGDTSTIYTNCWVTGWGFSKEKGEIQNILQKVNIPLVTNEECQKRYQD**F**KITQRMVCAGYKEGGKDACKGDSGGPLVCKHNGMWRLVGITSWGEGCARREQPGVYTKVAEYMDWILEKTQSSDGKAQMQSPA^622^HHHHHH

**K192R**

N^357^TGDNSVSTTKTSTRI^372^VGGTESSWGEWPWQVSLQVKLTAQRHLCGGSLIGHQWVLTAAHCFDGLPLQDVWRIYSGILELSDITKDTPFSQIKEIIIHQEYKVSEGNHDIALIKLQAPLEYTEFQKPISLPSKGDTSTIYTNCWVTGWGFSKEKGEIQNILQKVNIPLVTNEECQKRYQDYKITQRMVCAGYKEGGKDAC**R**GDSGGPLVCKHNGMWRLVGITSWGEGCARREQPGVYTKVAEYMDWILEKTQSSDGKAQMQSPA^622^HHHHHH

#### 3.15 Plasma Kallikrein Detection in Human Plasma

Human plasma was obtained from a single healthy donor using BD vacutainer sodium citrate tubes. Plasma was obtained following centrifugation of blood tubes at 1500 x g for 10 min at 4 °C. Plasma was then aliquoted and stored at -70 °C. The volunteer provided written informed consent, and all methods and experimental protocols were conducted in accordance with and approved by the NHS Research Ethics Committee (REC permit number 25/NI/0035). Prior to incubation with the covalent probes, the plasma was diluted 1:10 in PBS (10 mM, pH 7.4).

Reaction mixtures (25 μL) were prepared using the diluted plasma, which was spiked with 1 μM blood-purified human PK and 1 μM of peptide **14** or FP-TAMRA. The mixtures were incubated for 30 min at 37 °C. Following incubation, the samples were conjugated with TAMRA-azide using the CuAAC protocol described above, and protein labelling assessed by in-gel fluorescence.

#### 3.16 Peptide 14 Stability Tests

The hydrolytic stability of peptide **14** was evaluated in PBS (10 mM, pH 7.4) at RT. Peptide **14** was prepared at a final concentration of 500 μM. Fmoc-Lys(Boc)-OH (500 μM) was added as a control to serve as an internal standard for quantitative LC-MS analysis.

To capture the degradation kinetics, incubation times were sampled at 0 h, 4 h, 8 h, 12 h, 16 h, 24 h and 48 h. Samples were analysed by analytical LC-MS with an extended gradient (see **Supplementary Table 10**).

| **Time (min)** | **Solvent A (%)**  **(H_2_O + 0.1% FA)** | **Solvent B (%)**  **(MeCN + 0.1% FA)** |
| --- | --- | --- |
| 0.00 | 95 | 5 |
| 10.00 | 95 | 5 |
| 100.00 | 5 | 95 |
| 105.00 | 5 | 95 |
| 110.00 | 95 | 5 |
| 120.00 | 95 | 5 |

***Supplementary Table 12: Timetable of gradient used for analytical LC-MS of peptide 14 stability samples.***

The relative quantity of the remaining intact peptide at each timepoint was determined by integrating the area under the curve (AUC) of the UV signal at 254 nm and normalising it against the AUC of the Fmoc-Lys(Boc)-OH. Data processing was performed using Agilent OpenLab CDS Data Analysis Software Version 2.8. To calculate the hydrolytic half-life (*t_1/2_*), the normalised percentage of remaining intact peptide was plotted against time and fitted to a one-phase exponential decay model using GraphPad Prism.

### 4 Supplementary Figures

#### 4.1 Mechanism for Cyclisation

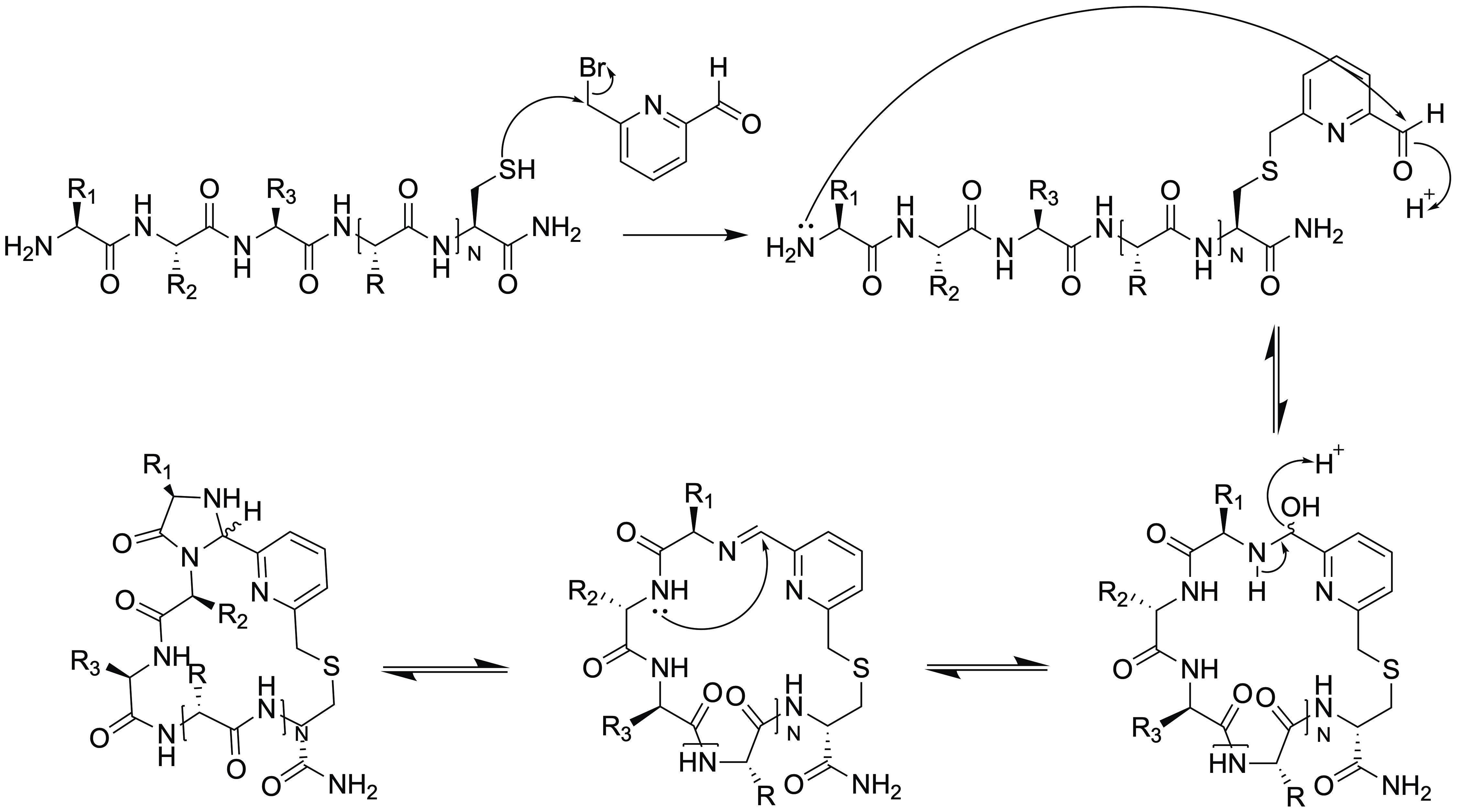

***Supplementary Figure 1: Mechanism for linear peptide cyclisation with the* 6-(bromomethyl)Picolinaldehyde (BMP) *linker.*** *The linker first alkylates at the cysteine side chain. The peptide N-terminus then attacks the linker aldehyde to form the imidazolidinone structure, proceeding via a planar imine intermediate.*

#### 4.2 NMR Characterisation of Test Peptide ALYWQHTC

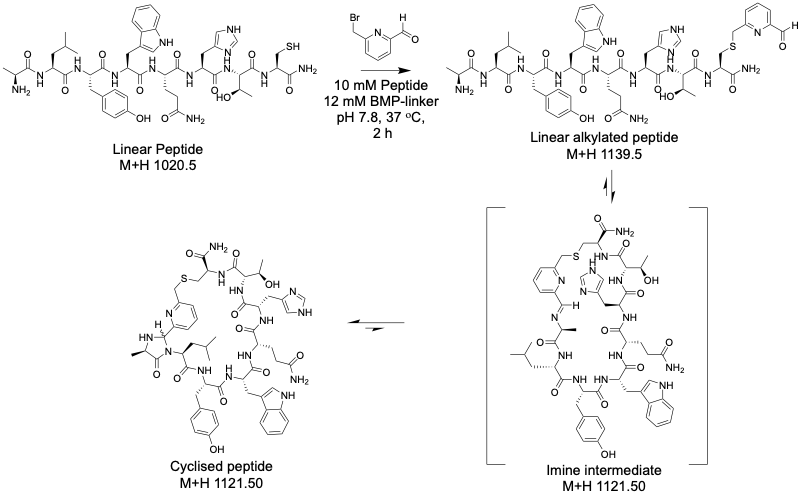

***Supplementary Figure 2: Sample preparation for NMR characterisation of test peptide ALYWQHTC. (A)*** *Reaction scheme for cyclisation with BMP linker. A sample of the linear peptide and a sample of the peptide purified after two hours cyclising were both submitted for 1 and 2 D NMR analysis.* ***(B)*** *Analytical low-resolution LC-MS of the linear and cyclised ALYWQHTC samples for NMR analysis.*

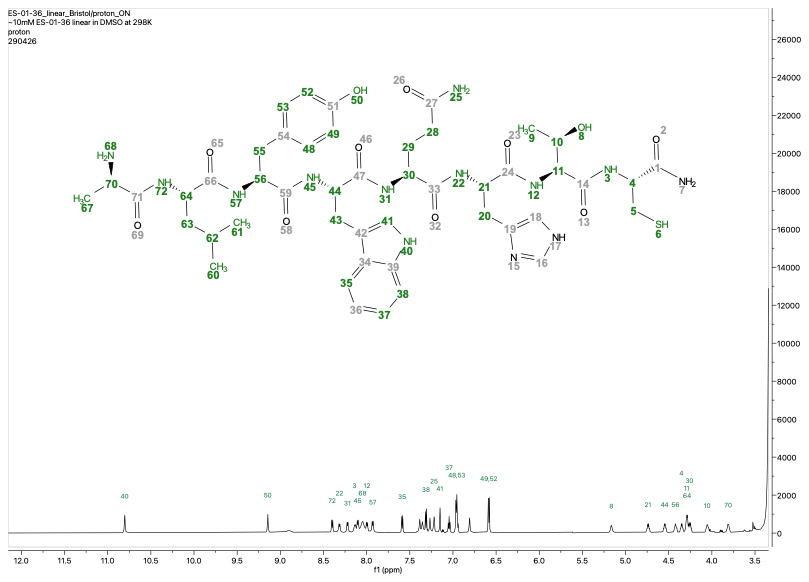

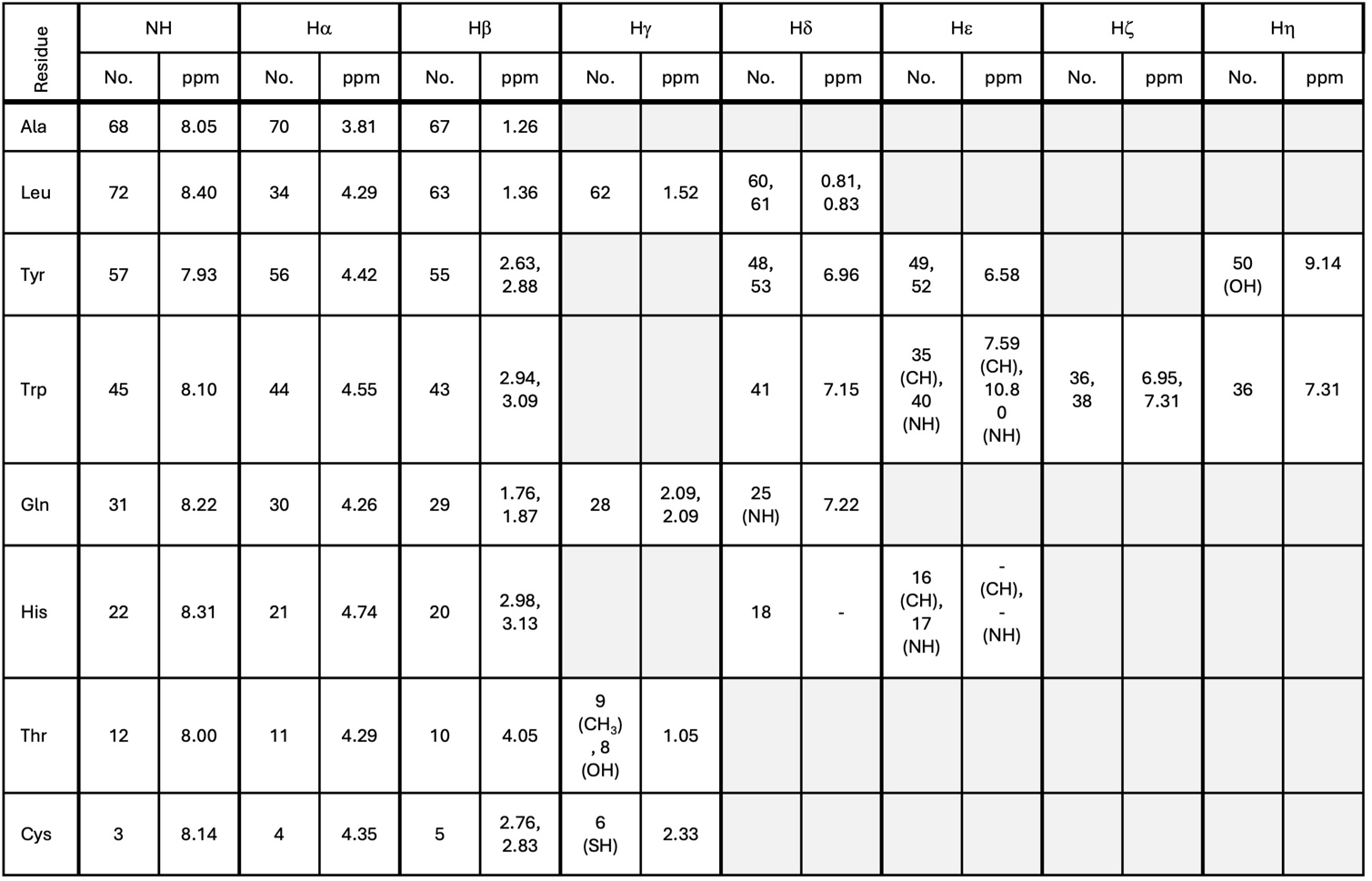
***Supplementary Figure 3: ^1^H NMR Chemical shift assignments of linear ALYWQHTC.***

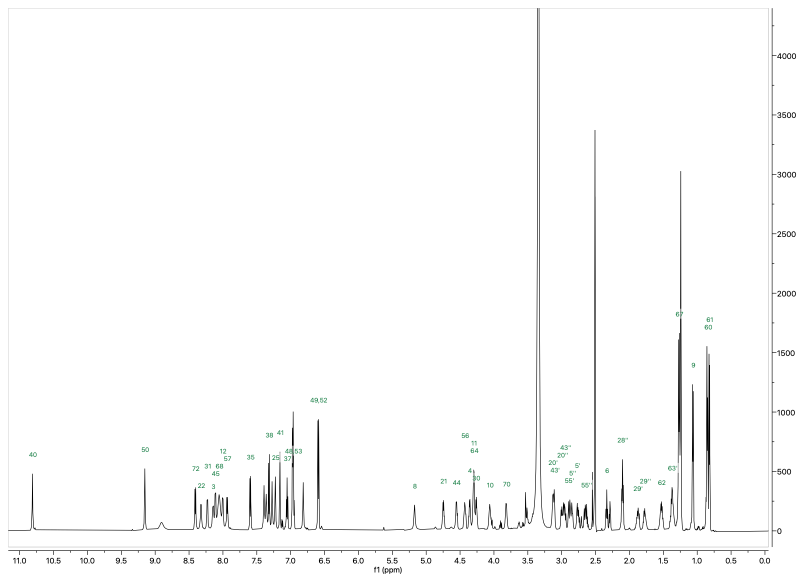
***Supplementary Figure 4: ^1^H spectrum of linear ALYWQHTC.*** *Peptide was dissolved in d_6_-DMSO (10 mM) and spectra acquired at 298K on a Bruker Avance III HD spectrometer operating at 700 MHz equipped with a 1.7 mm TCI microcryocoil probe.*

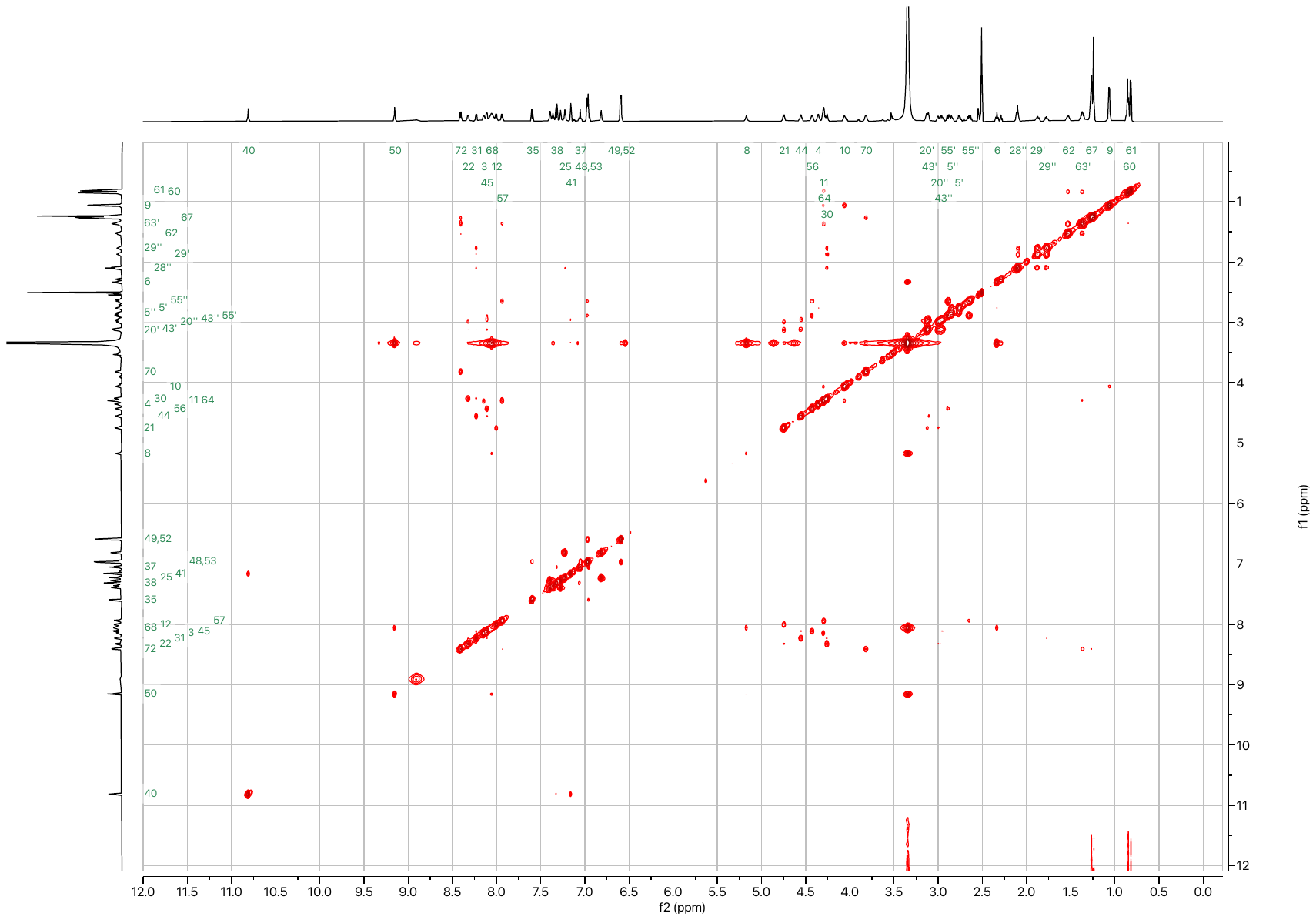

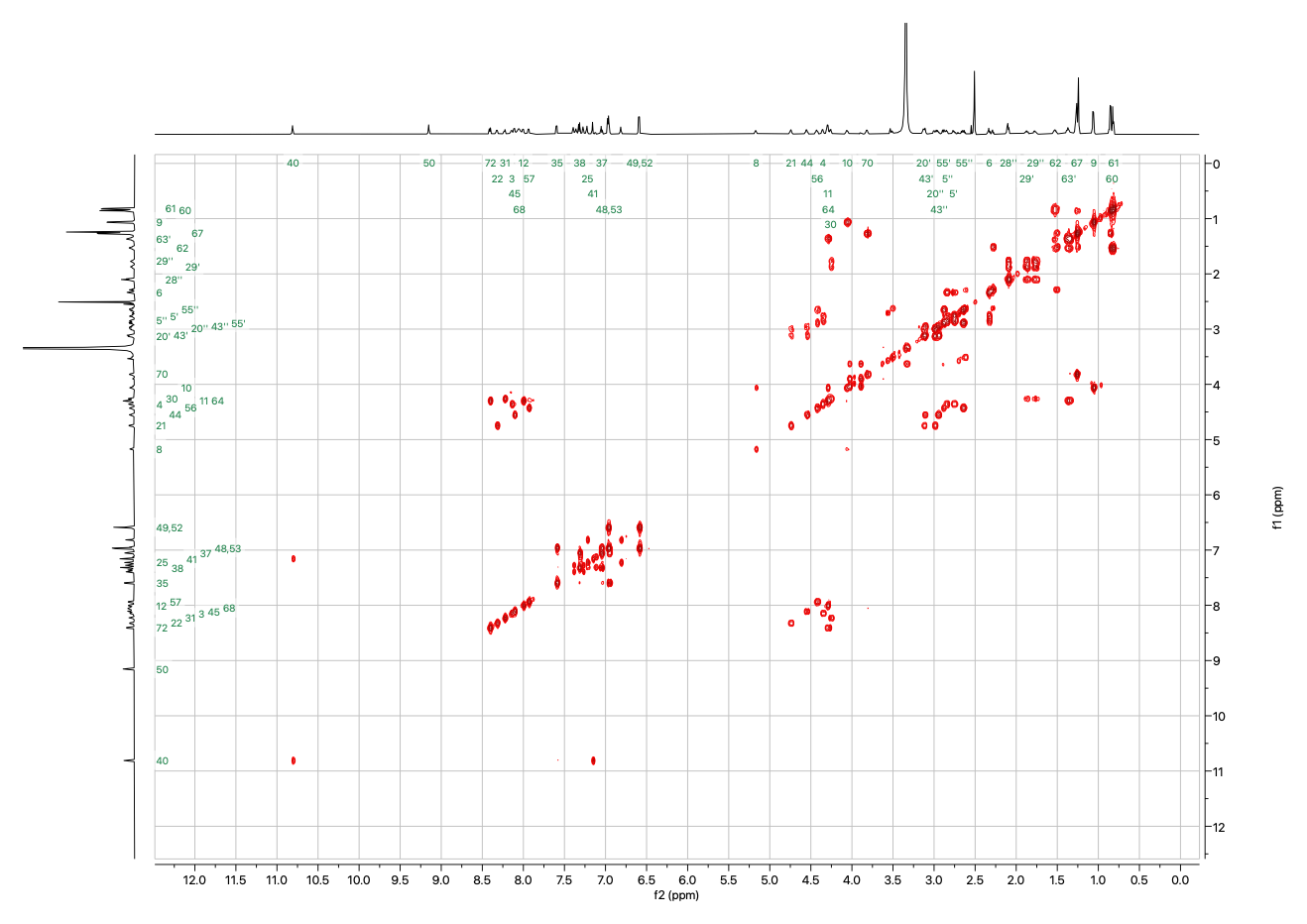
***Supplementary Figure 5: COSY spectrum of linear ALYWQHTC.***

***Supplementary Figure 6: NOESY spectrum of linear ALYWQHTC.***

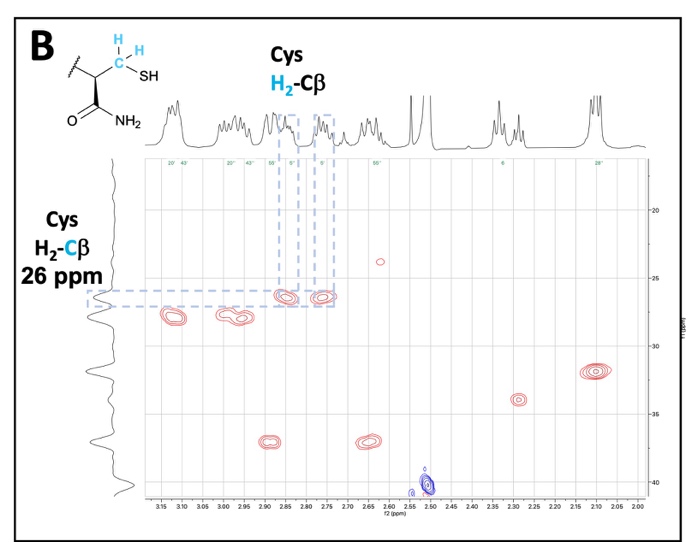

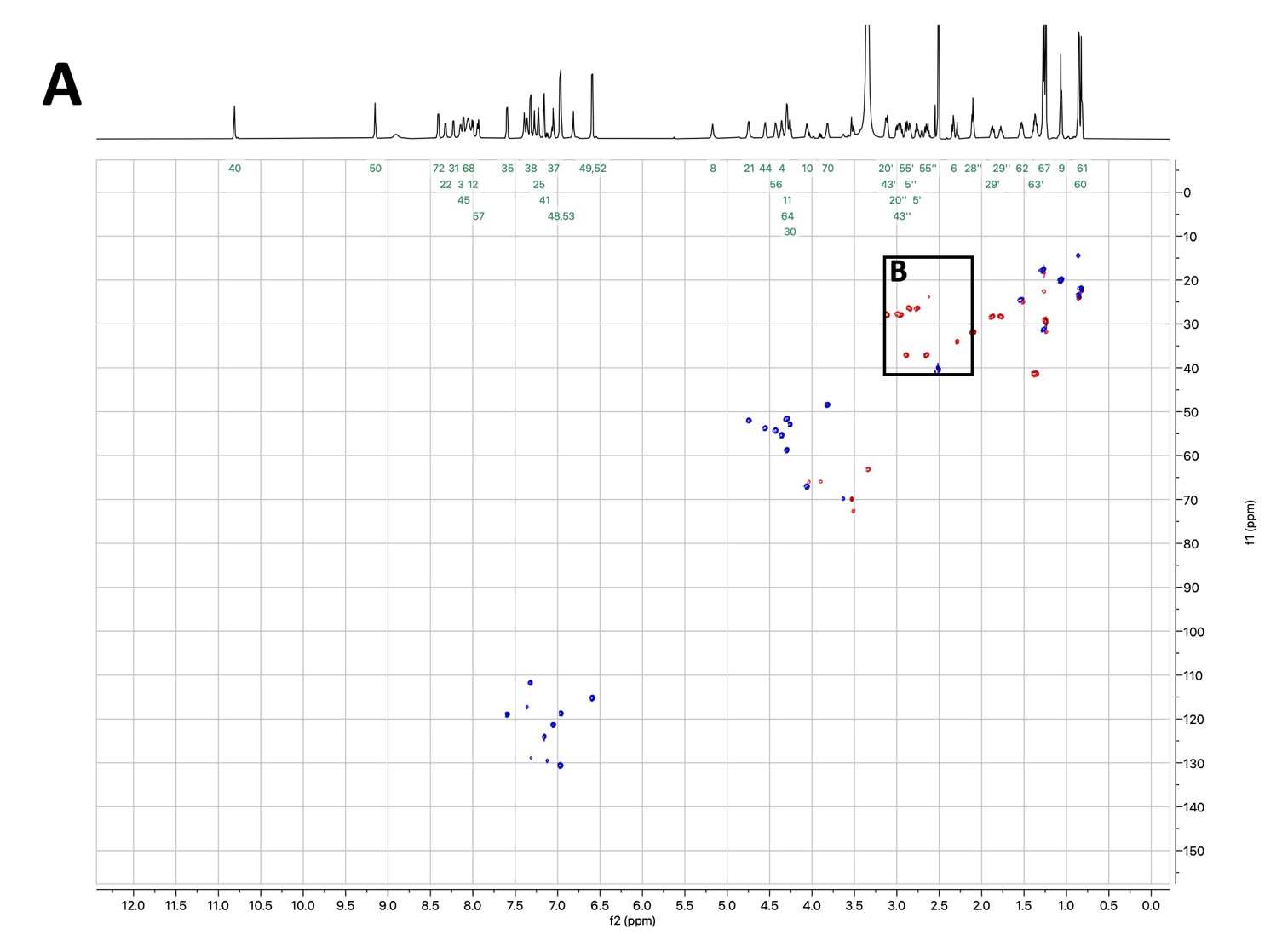

***Supplementary Figure 7: ^13^C-HSQC spectrum of linear ALYWQHTC. (A)*** *Full spectrum.* ***(B)*** *Excerpt showing 1-bond coupling between the cysteine β-protons and carbon. The cysteine β-carbon is upfield relative to the β-carbon of the cyclised peptide. This is indicative of cysteine alkylation during cyclisation.*

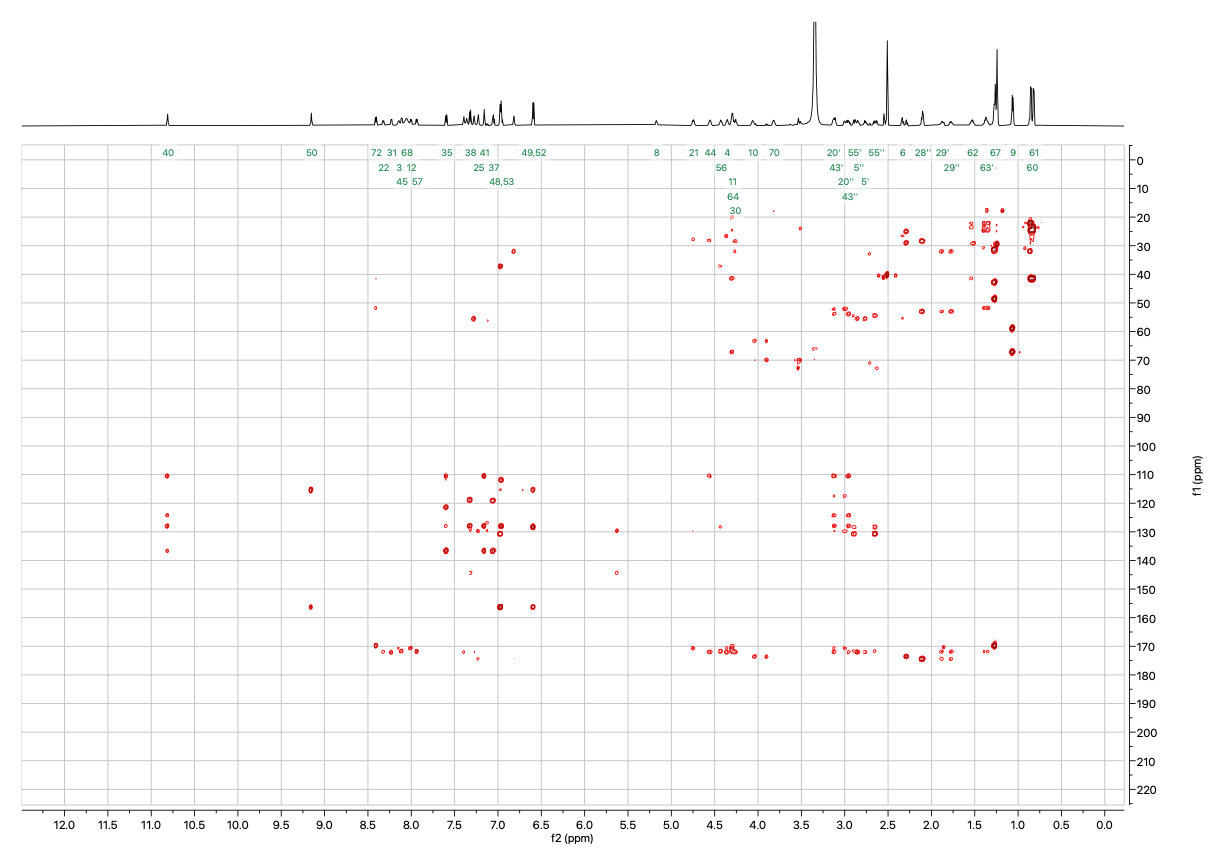

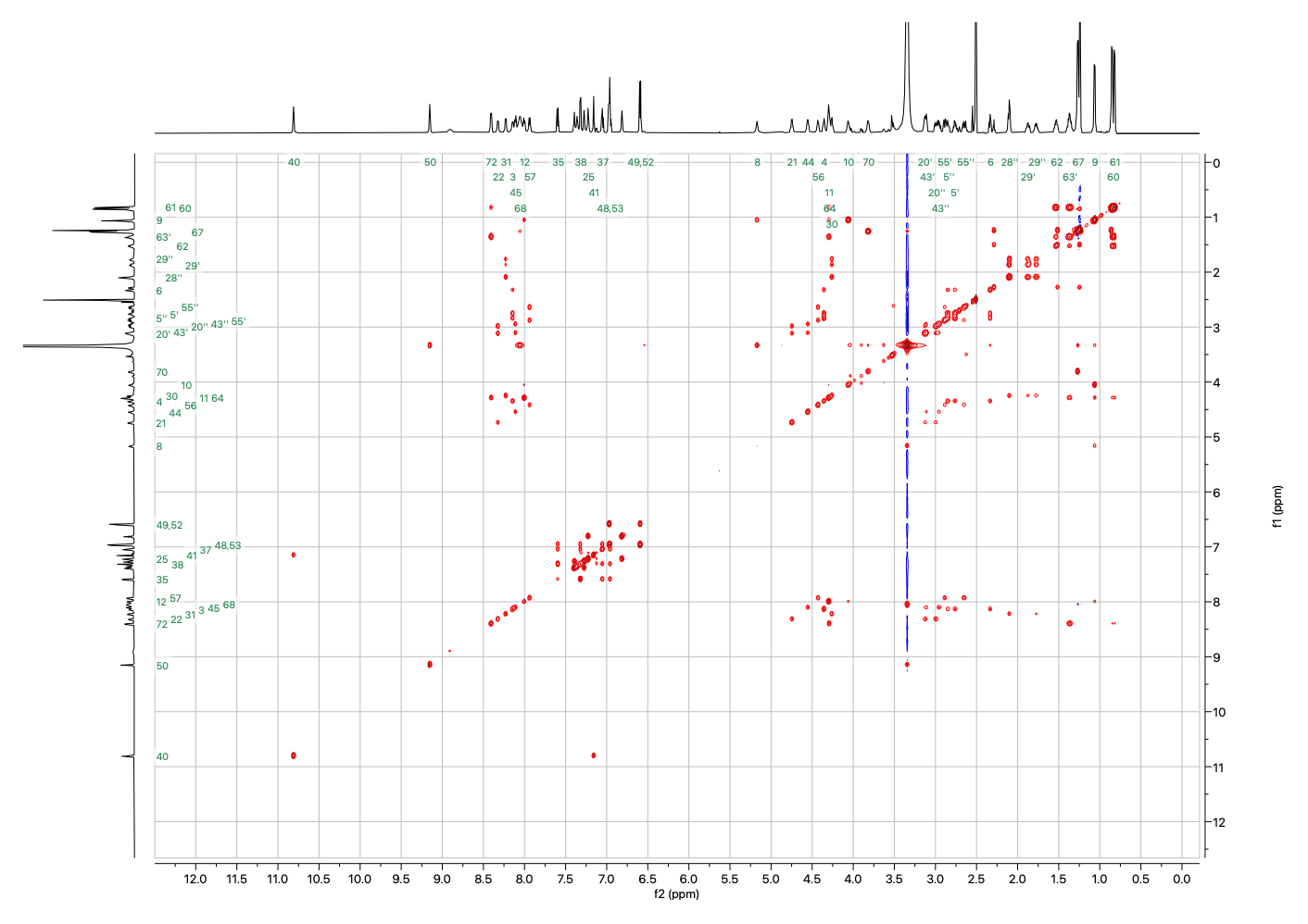
***Supplementary Figure 8: ^13^C-HMBC spectrum of linear ALYWQHTC.***

***Supplementary Figure 9: TOCSY spectrum of linear ALYWQHTC.***

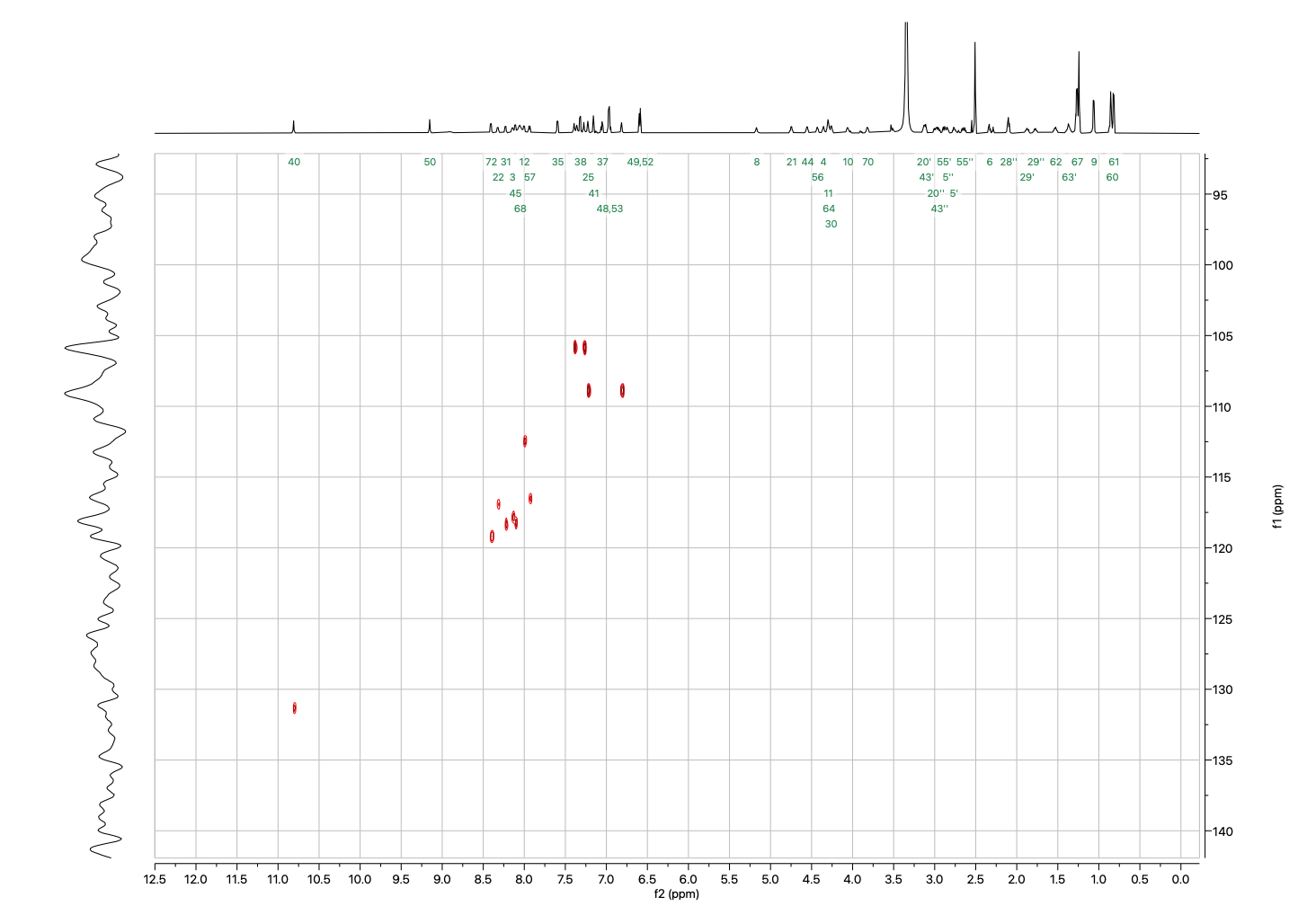

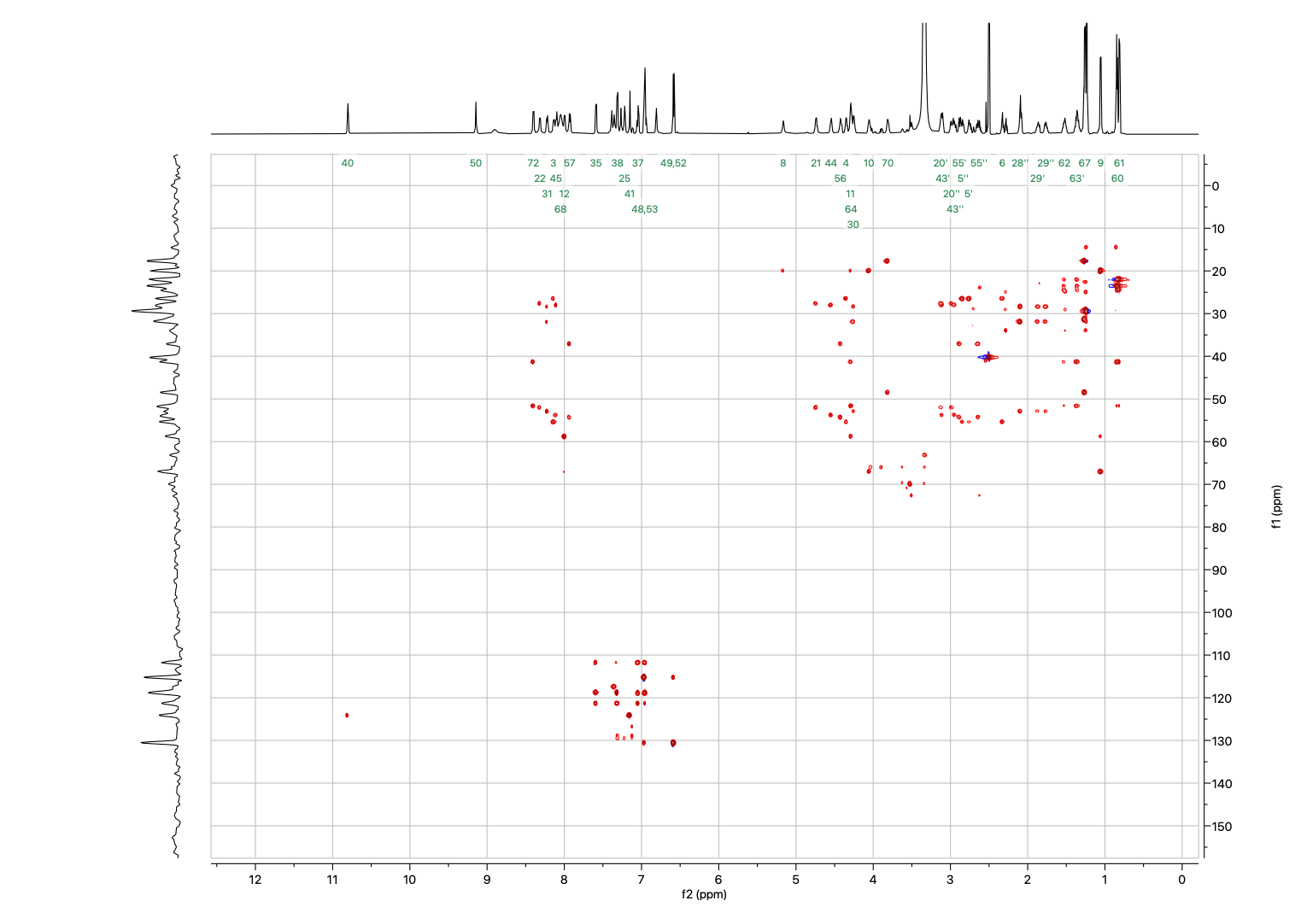
***Supplementary Figure 10: ^13^C-HSQC-TOCSY spectrum of linear ALYWQHTC.***

***Supplementary Figure 11: ^15^N-HSQC spectrum of linear ALYWQHTC.***

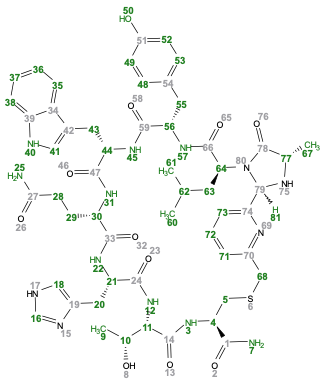

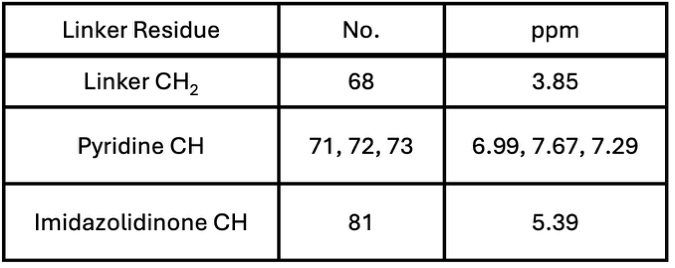

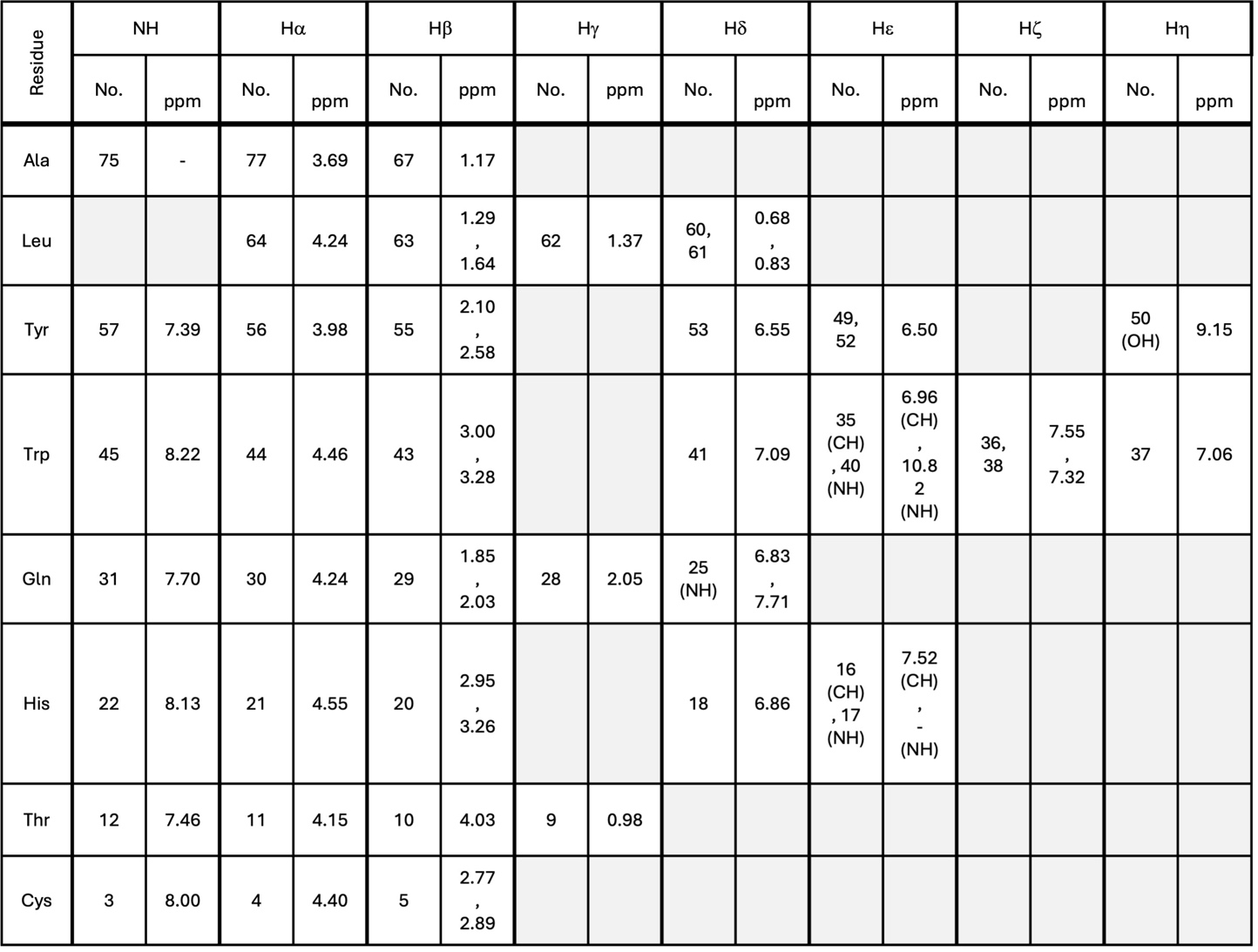

***Supplementary Figure 12: ^1^H NMR chemical shift assignments of cyclised ALYWQHTC.***

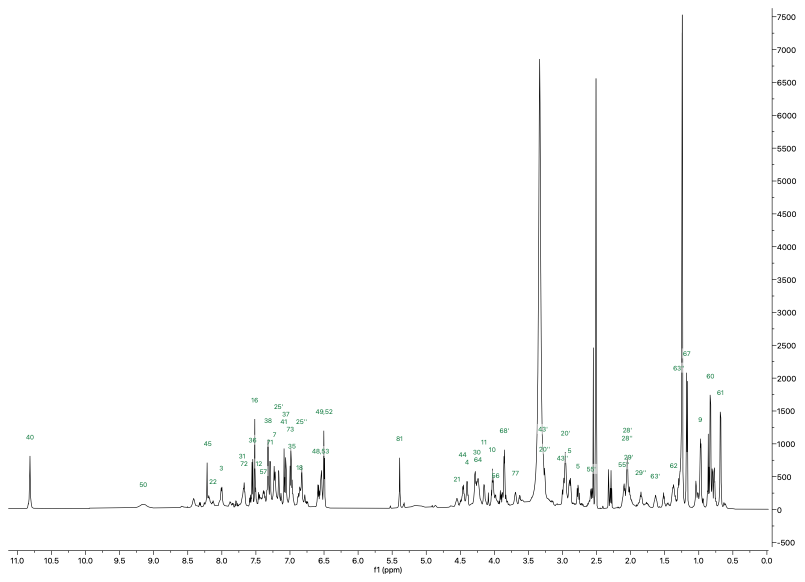
***Supplementary Figure 13: ^1^H spectrum of cyclised ALYWQHTC.*** *Peptide was dissolved in d_6_-DMSO (10 mM) and spectra acquired at 298K on a Bruker Avance III HD spectrometer operating at 700 MHz equipped with a 1.7 mm TCI microcryocoil probe.*

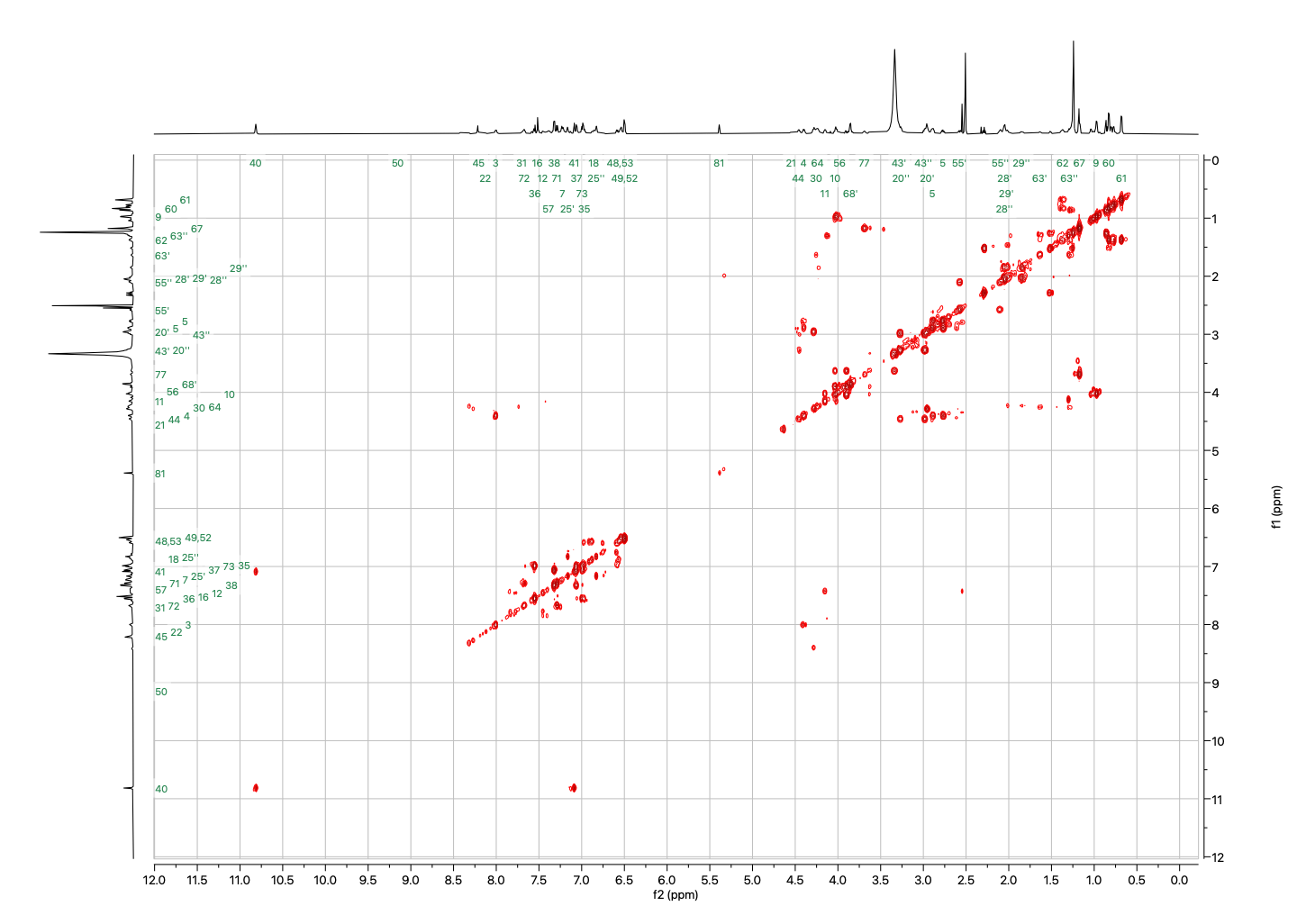

***Supplementary Figure 14: COSY spectrum of cyclised ALYWQHTC.***

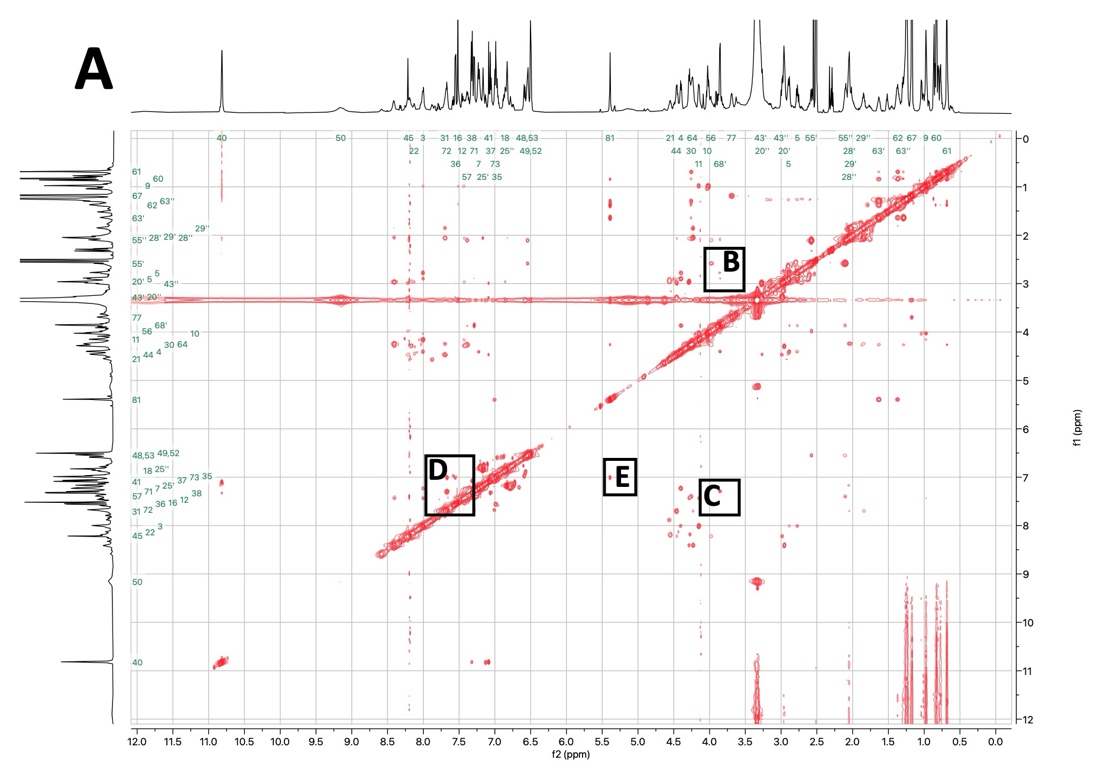

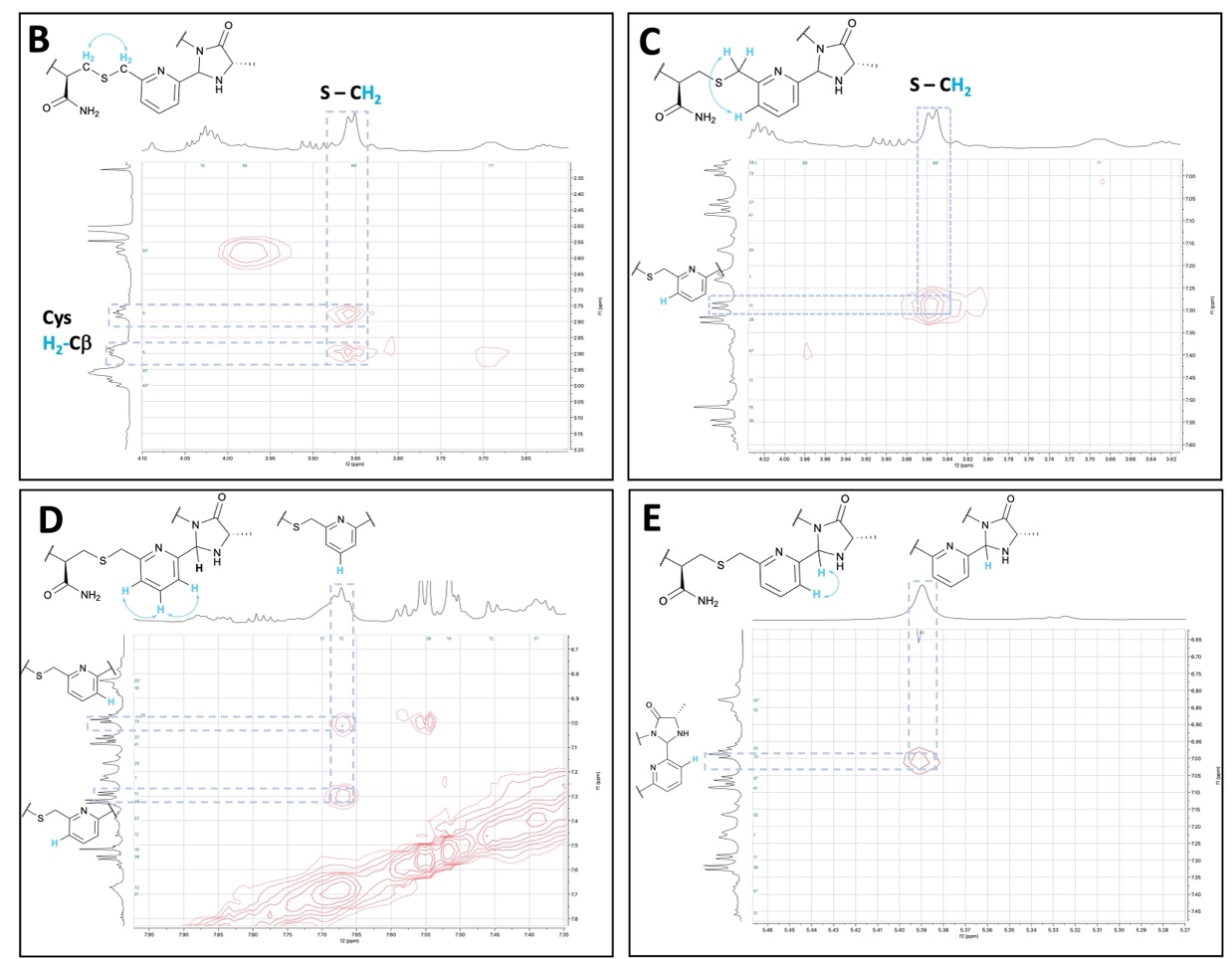

***Supplementary Figure 15: NOESY spectrum of cyclised ALYWQHTC. (A)*** *Full spectrum.* ***(B)-(E)*** *Excerpts showing interactions that run from the cysteine β-protons, around the pyridine of the linker and to the characteristic imidazolidinone proton formed.*

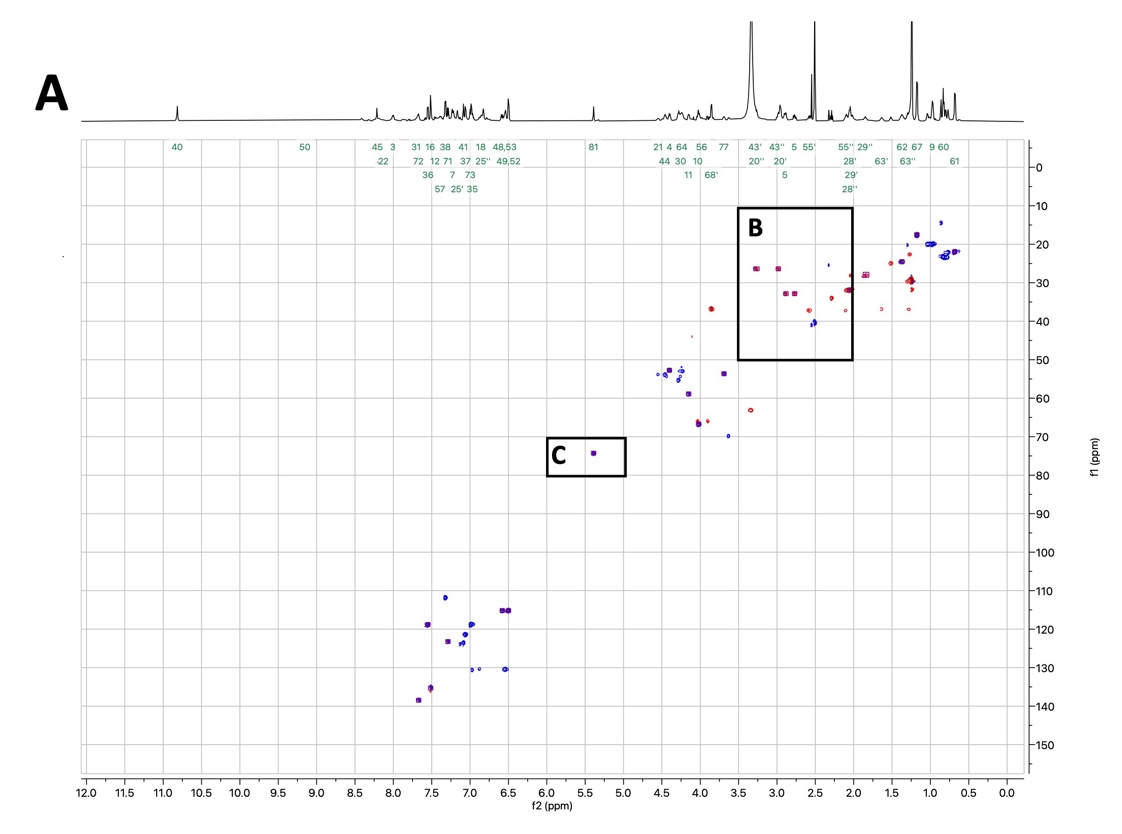

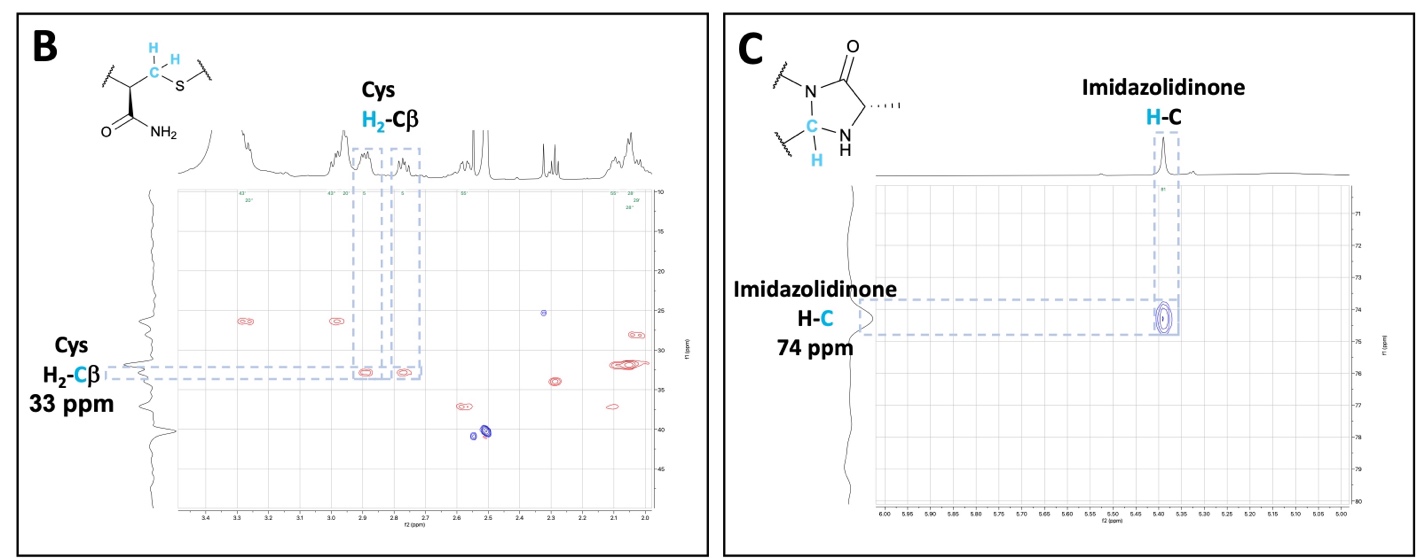

***Supplementary Figure 16: ^13^C-HSQC spectrum of cyclised ALYWQHTC. (A)*** *Full spectra.* ***(B)*** *Excerpt showing 1-bond coupling between the cysteine β-protons and carbon. This highlights the downfield shift of the β-carbon relative to the linear peptide, indicative of the cysteine alkylation and the presence of a second electron-donating CH_2_ group in proximity to the cysteine following cyclisation.* ***(C)*** *Excerpt showing 1-bond coupling between the imidazolidinone proton and carbon, both with chemical shifts characteristic of the imidazolidinone structure (5.39 ppm and 74 ppm, respectively).*

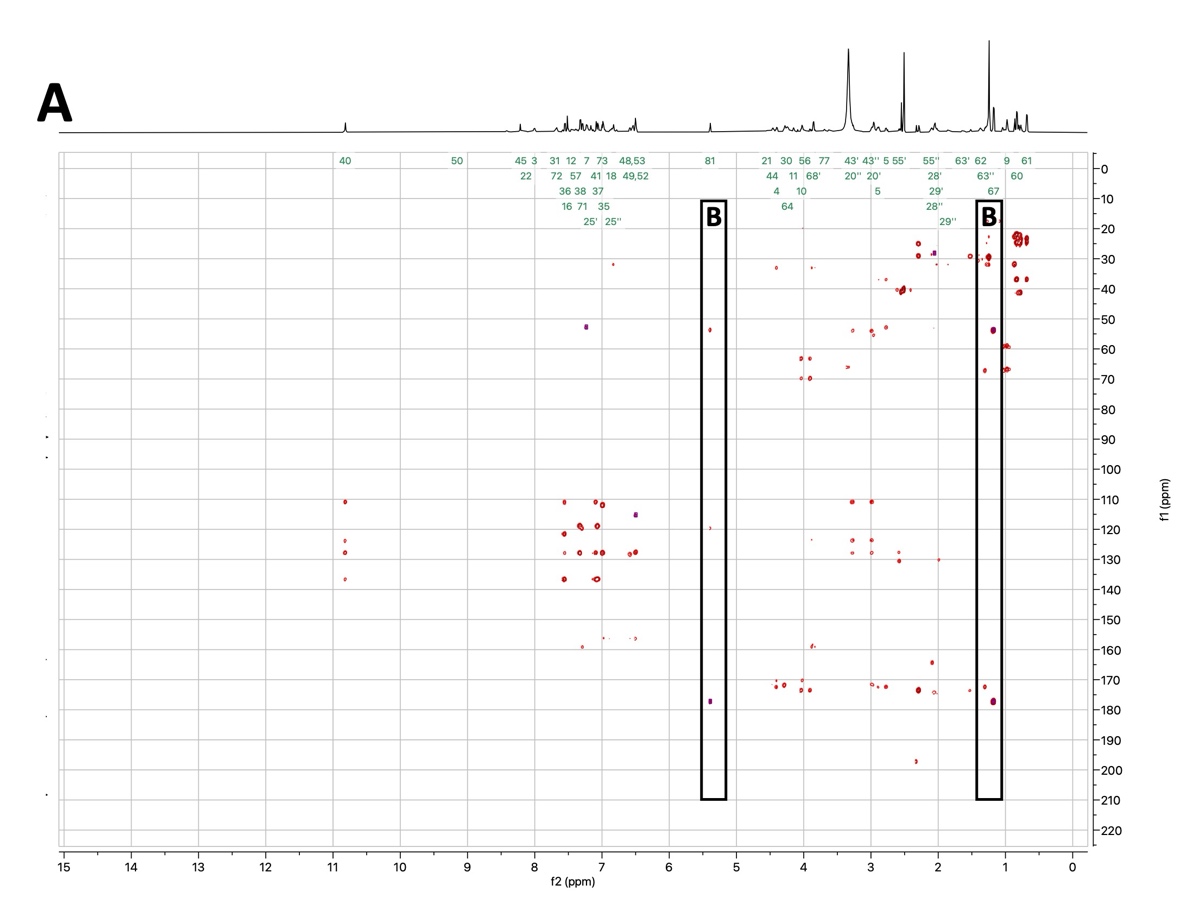

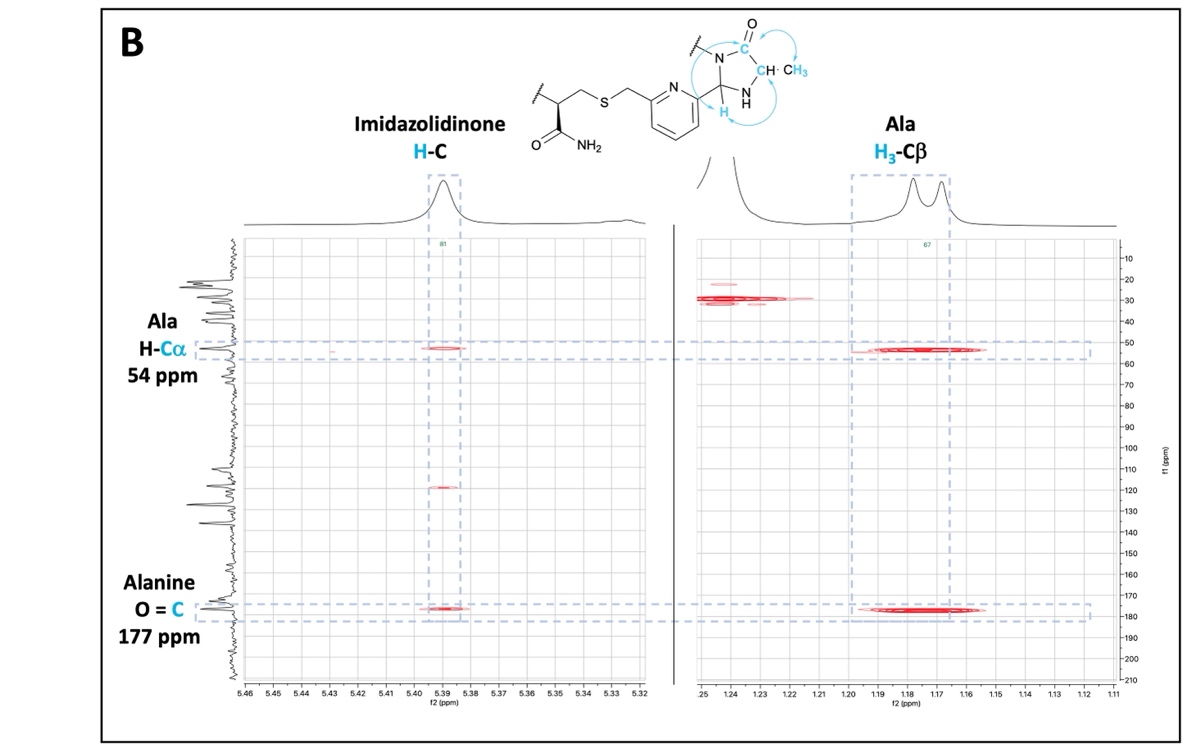

***Supplementary Figure 17: ^13^C-HMBC Spectrum of cyclised ALYWQHTC. (A)*** *Full spectra.* ***(B)*** *Excerpt showing 3-bond couplings between the characteristic imidazolidinone proton and the alanine Cα and C=O. A coupling can also be seen between the C=O and the characteristic alanine β-protons (CH_3_), confirming formation of the imidazolidinone with the alanine N-terminus.*

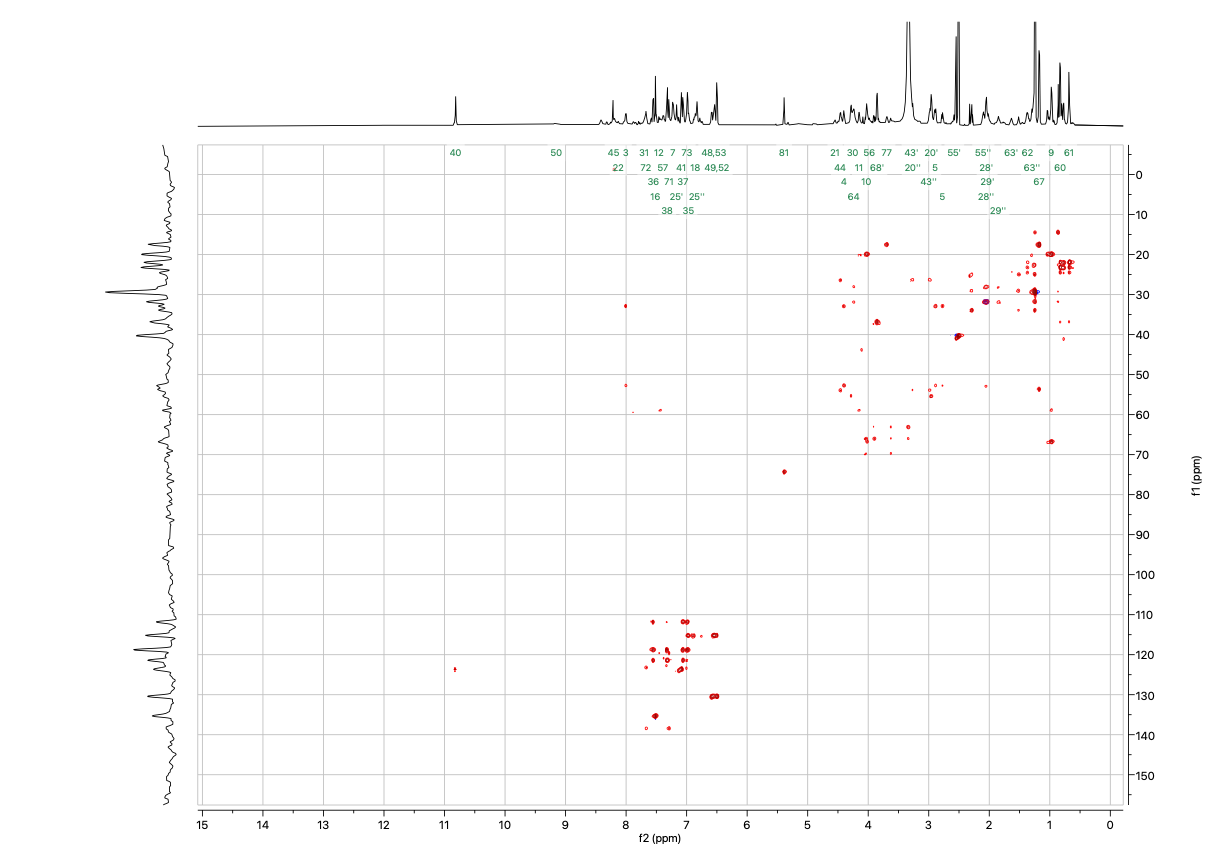

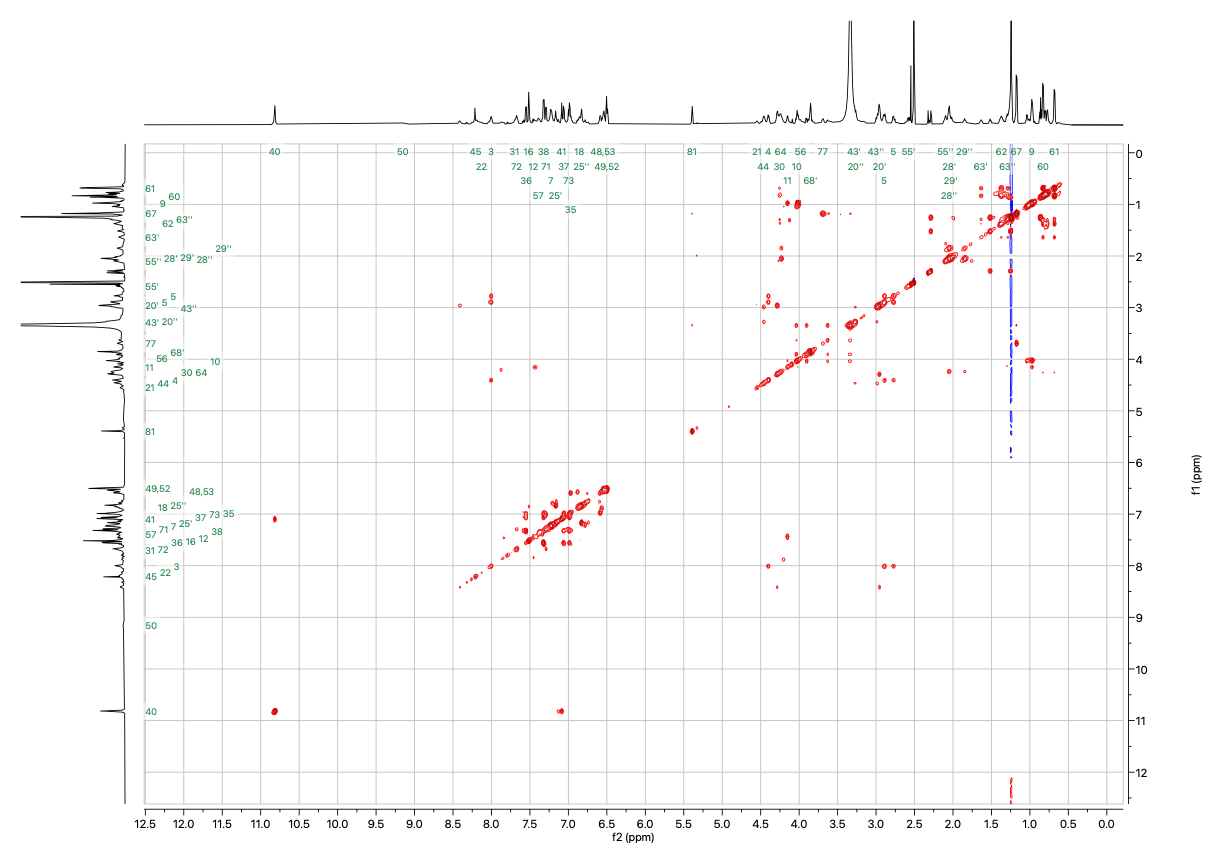
***Supplementary Figure 18: TOCSY spectrum of cyclised ALYWQHTC.***

***Supplementary Figure 19: ^13^C-HSQC-TOCSY spectrum of cyclised ALYWQHTC.***

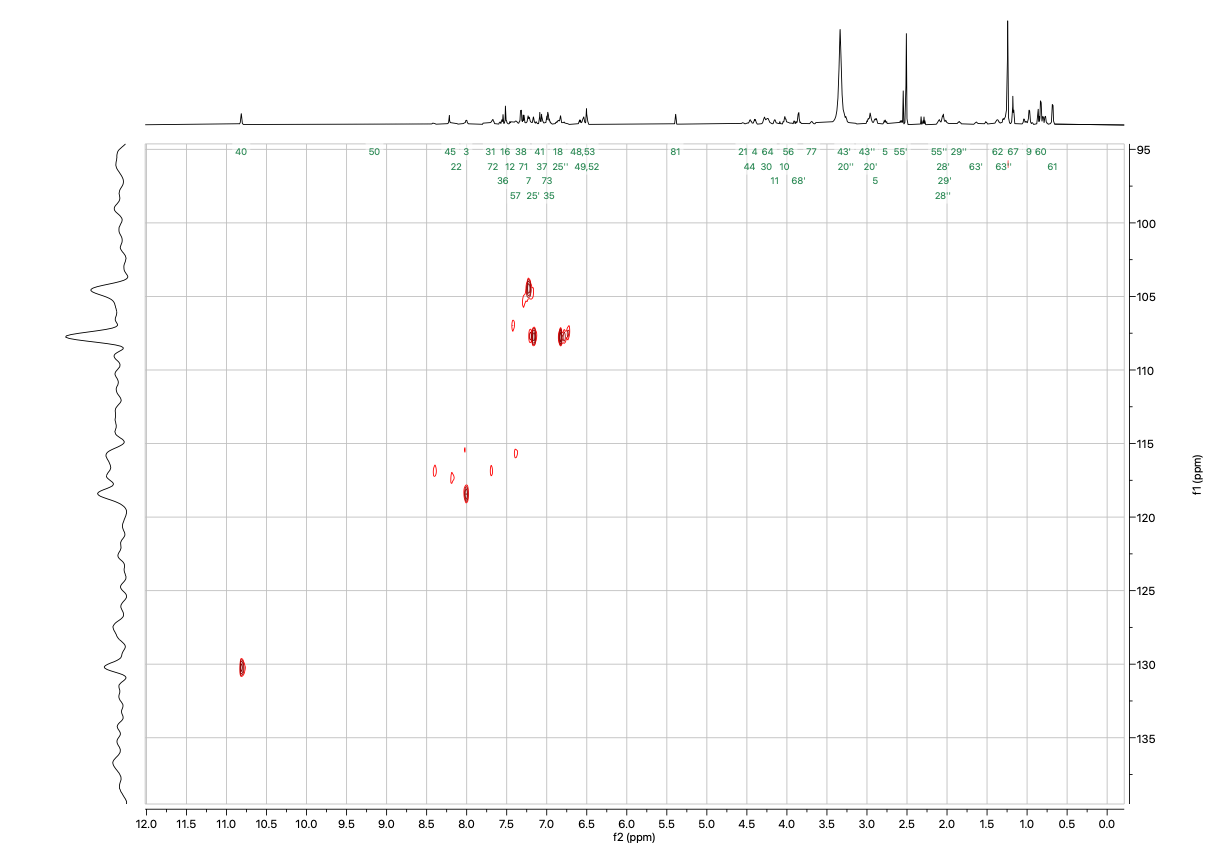
***Supplementary Figure 20: ^15^N-HSQC spectrum of cyclised ALYWQHTC.***

#### 4.3 Stability of Test Peptides under Aqueous Conditions

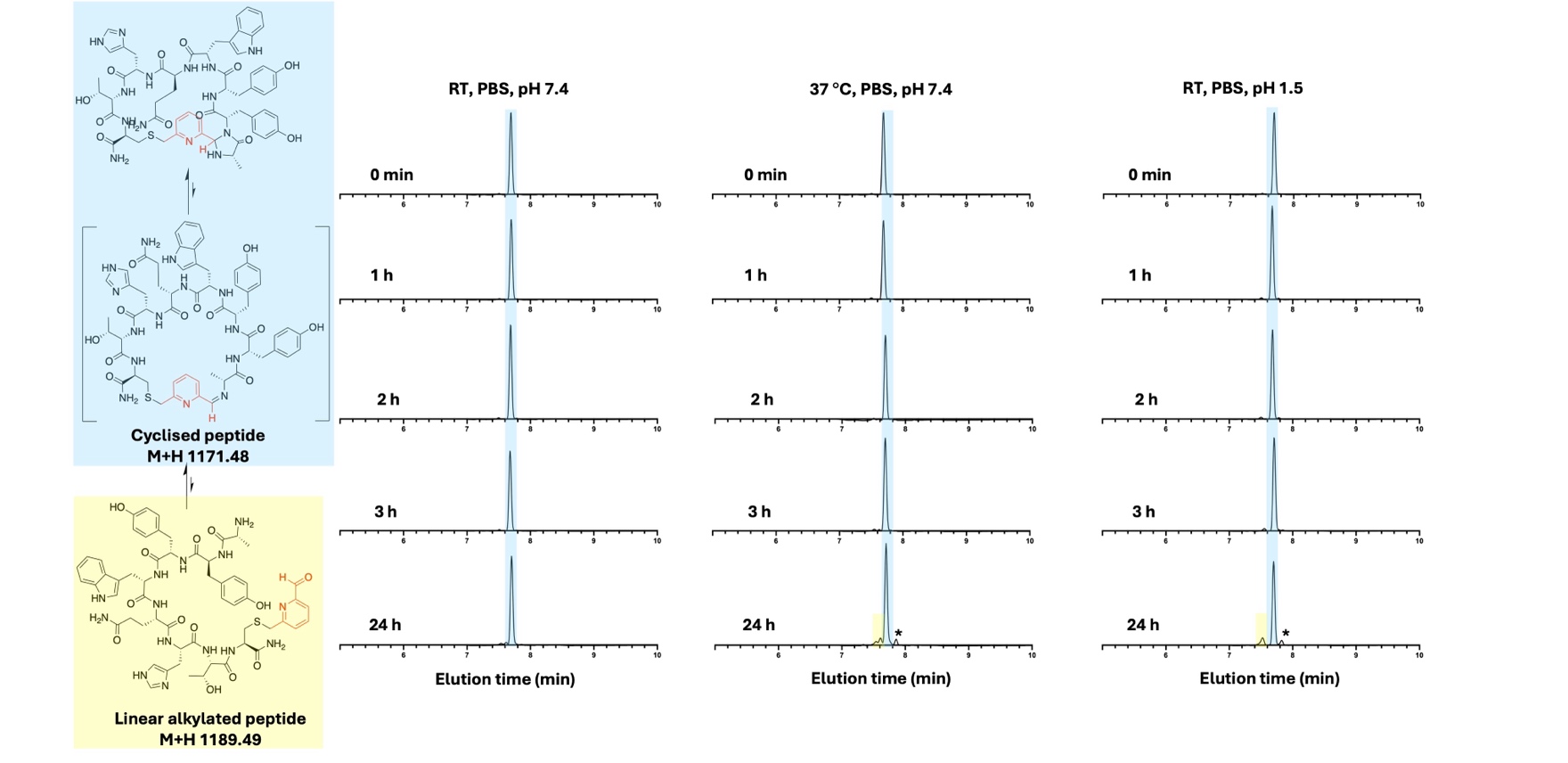

***Supplementary Figure 21: Stability of the peptide sequence AYYWQHTC cyclised with the BMP-linker.*** *UV traces (280 nm) are shown for monitoring reversion to alkylated peptide at near neutral pH at both RT* ***(A)*** *and 37* °C ***(B)*** *and under highly acidic conditions at RT* ***(C)****.* *Species with mass corresponding to cyclised peptide are highlighted in blue (M+H obs 1172.0), species with mass corresponding to linear alkylated peptide are highlighted in yellow (M+H obs 1189.8). * corresponds to a species with Mobs 601.1.*

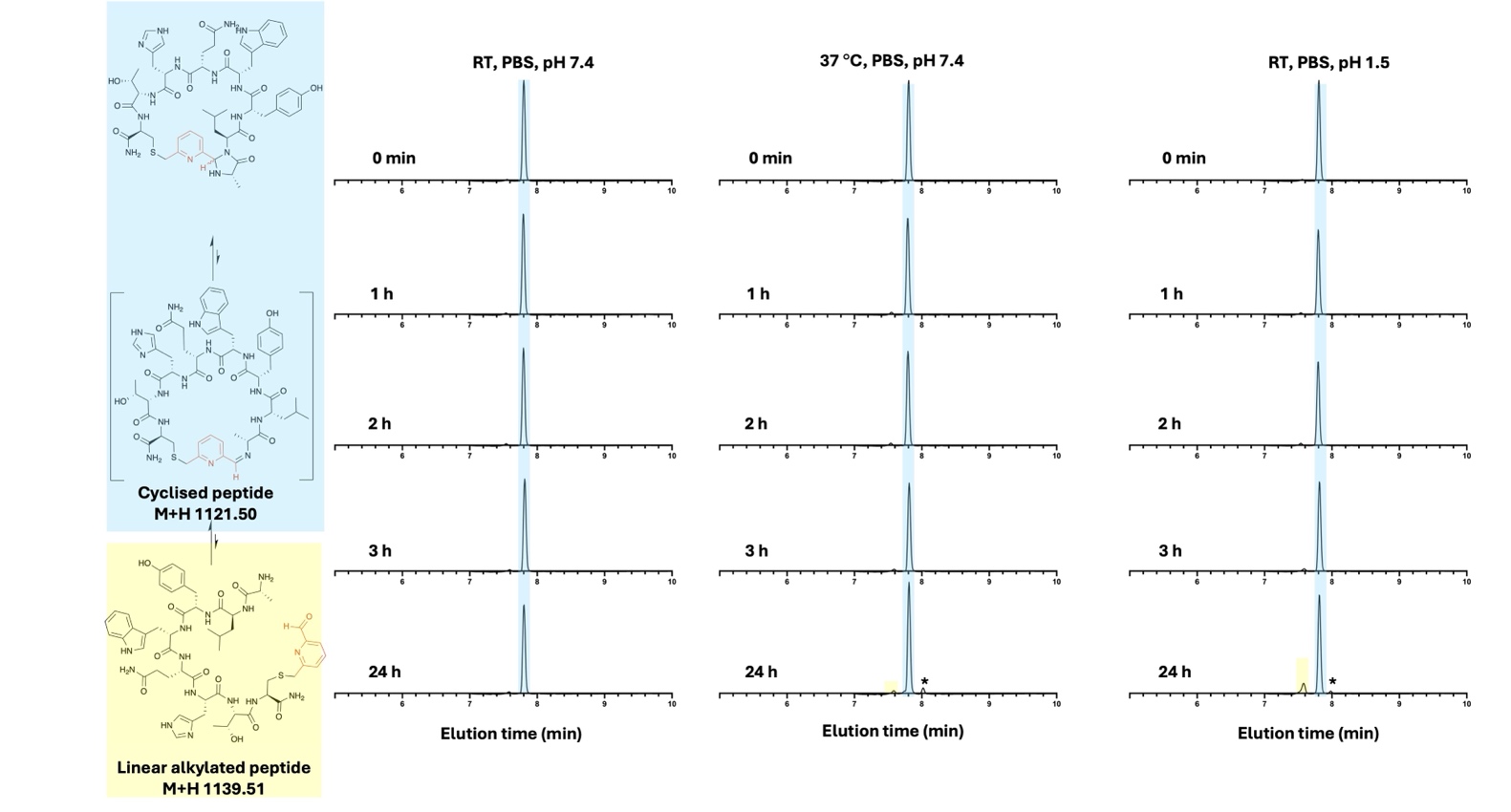

***Supplementary Figure 22: Stability of the peptide sequence ALYWQHTC cyclised with the BMP-linker.*** *UV traces (280 nm) are shown for monitoring reversion to alkylated peptide at near neutral pH at both RT* ***(A)*** *and 37* °C ***(B)*** *and under highly acidic conditions at RT* ***(C)****.* *Species with mass corresponding to cyclised peptide are highlighted in blue (M+H obs 1121.6), species with mass corresponding to linear alkylated peptide are highlighted in yellow (M+H obs 1140.0). * corresponds to a species with Mobs 576.1.*

#### 4.4 Reduction of Test Peptide ALYWQHTC with NaCNBH_3_

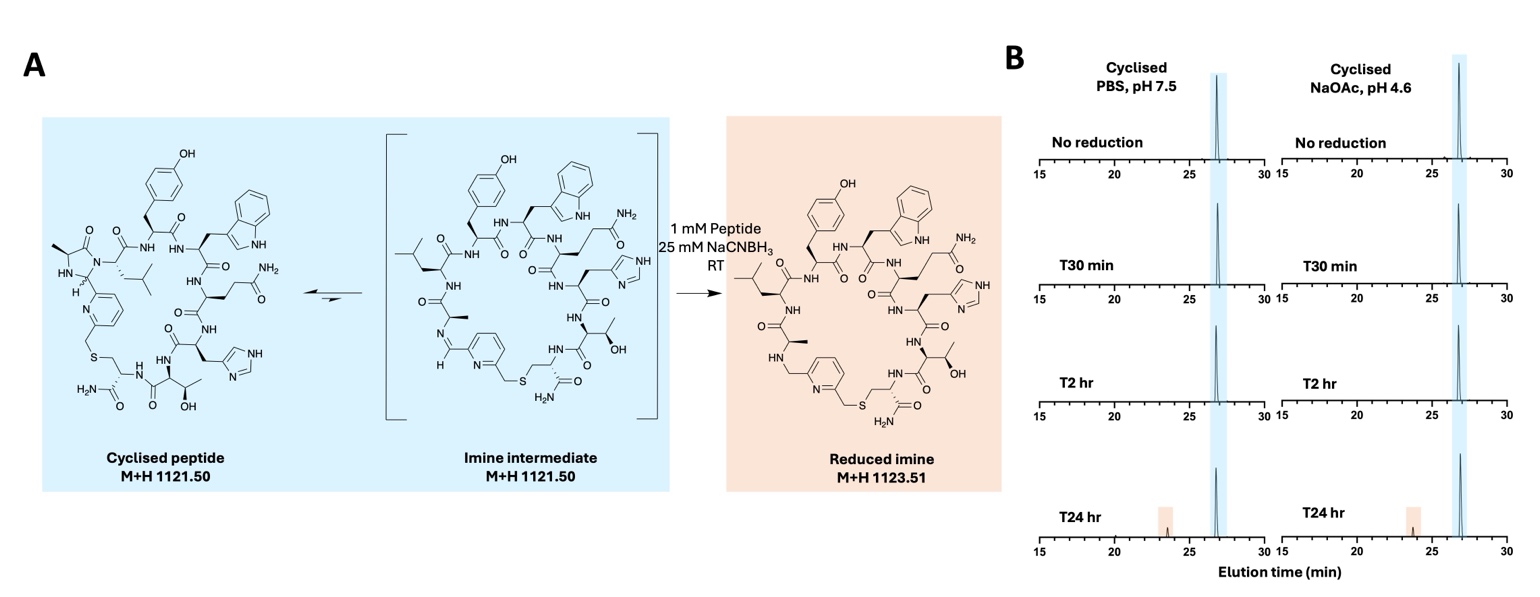

***Supplementary Figure 23: Reduction of test peptide ALYWQHTC. (A)*** *Reaction scheme for capturing imine formed from the purified cyclised peptide by irreversible reduction with NaCNBH_3_.* ***(B)*** *UV traces (280 nm) from monitoring the capture of reduced imine formed from purified cyclised peptide. Species with mass corresponding to cyclised peptide are highlighted in blue (M+H obs 1121.6), species with mass corresponding to reduced imine are highlighted in red (M+H obs 1123.8).*

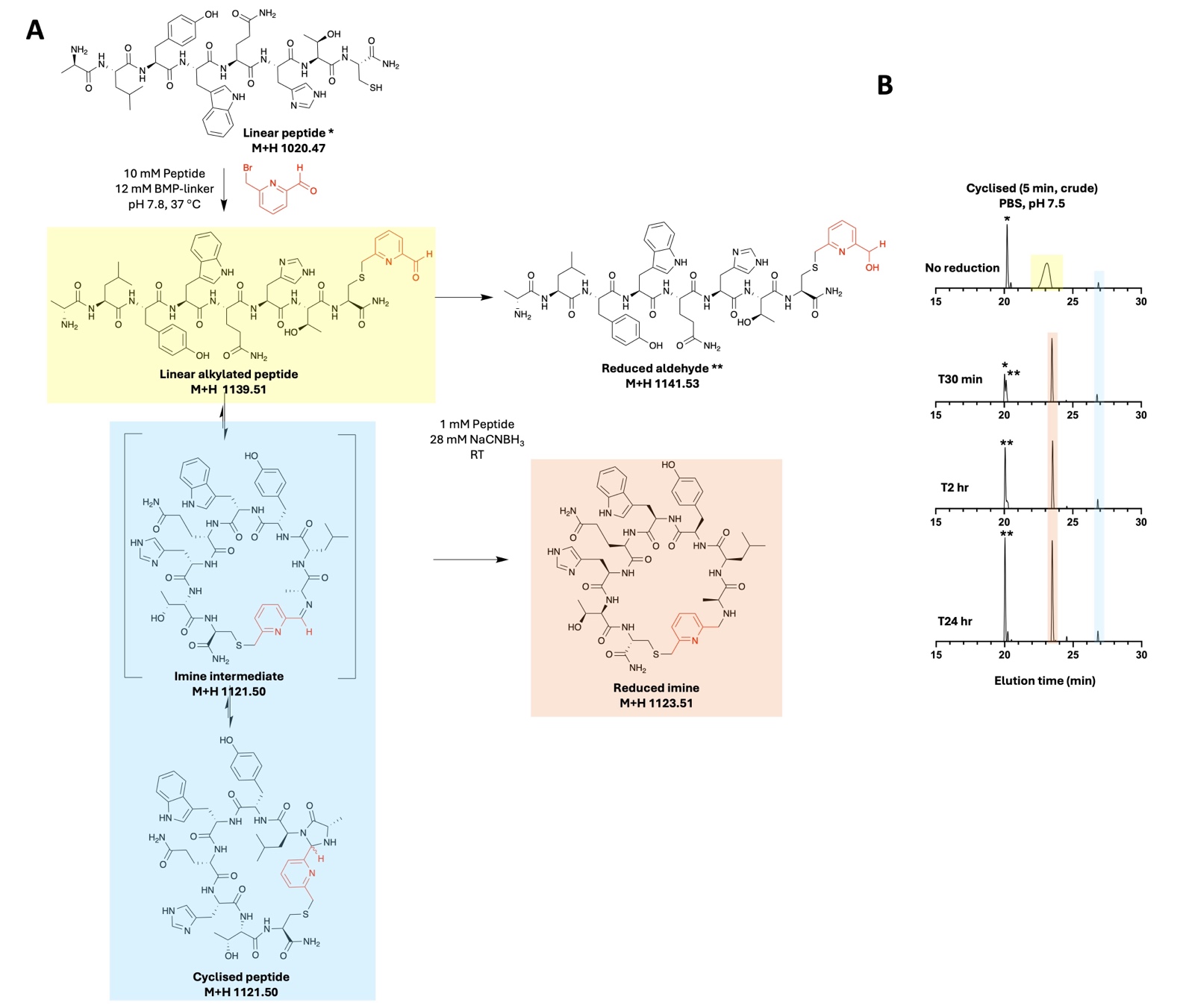

***Supplementary Figure 24: Confirming reduction conditions for capturing imine intermediate. (A)*** *Reaction scheme for capturing the imine intermediate formed from the crude cyclisation reaction after peptide alkylation, by irreversible reduction with NaCNBH_3_.* ***(B)*** *UV traces (280 nm) from monitoring the capture of reduced imine formed from crude cyclisation reaction. Species with mass corresponding to cyclised peptide are highlighted in blue (M+H obs 1121.6), species with mass corresponding to linear alkylated peptide are highlighted in yellow (M+H obs 1140.0), species with mass corresponding to reduced imine are highlighted in red (M+H obs 1124.0). * corresponds to linear peptide (Mobs+H obs 1020.8), ** corresponds to reduced alkylated peptide (Mobs+H obs 1142.0).*

#### 4.5 Expression and Purification of Peptide-pIII Fusion Protein

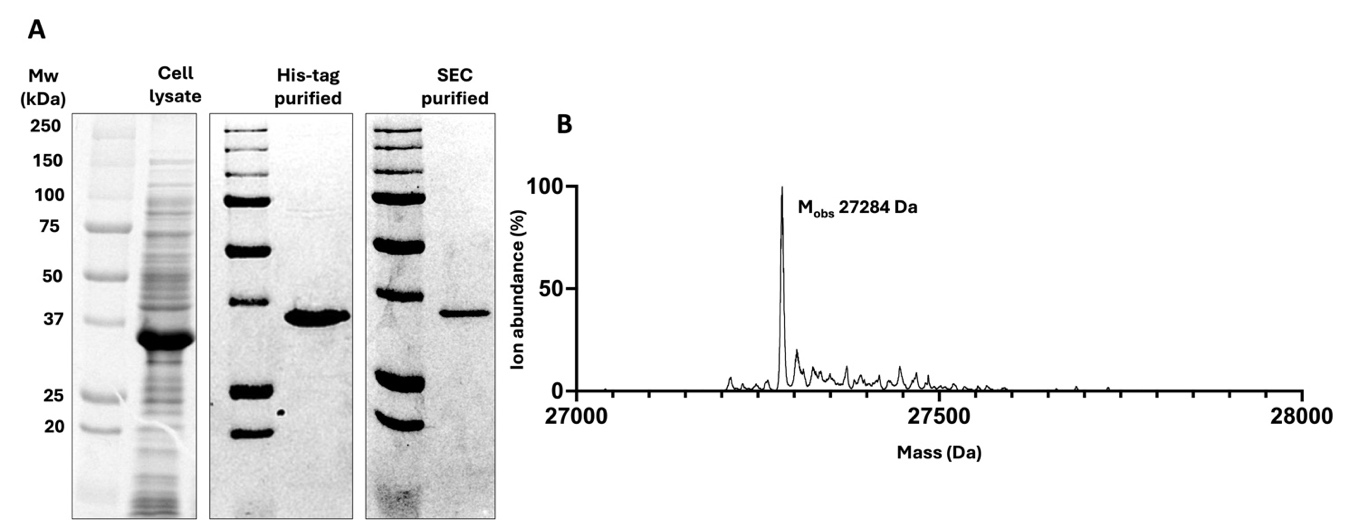

***Supplementary Figure 25: Expression and purification of AAYWQHTC-pIII fusion protein. (A)*** *SDS-PAGE gels after cell lysis, His-tag purification and size-exclusion chromatography purification.* ***(B)*** *Mass spectrometry analysis of the unmodified construct after purification.*

#### 4.6 Cyclisation Toxicity Titers

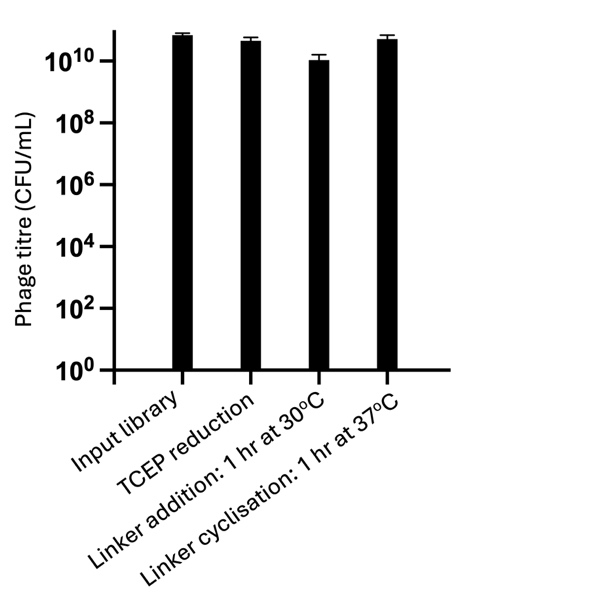

***Supplementary Figure 26: Phage titers for the BMP-cyclisation strategy.*** *Toxicity of the proposed cyclisation strategy was assessed by monitoring the number of phage after reduction with TCEP (2 mM, 30 min, RT), BMP linker addition (40 µM, 1 h, 30 °C, 20% MeCN) and further BMP cyclisation (1 h, 37 °C, no MeCN).*

#### 4.7 Volcano Plot of NGS Data

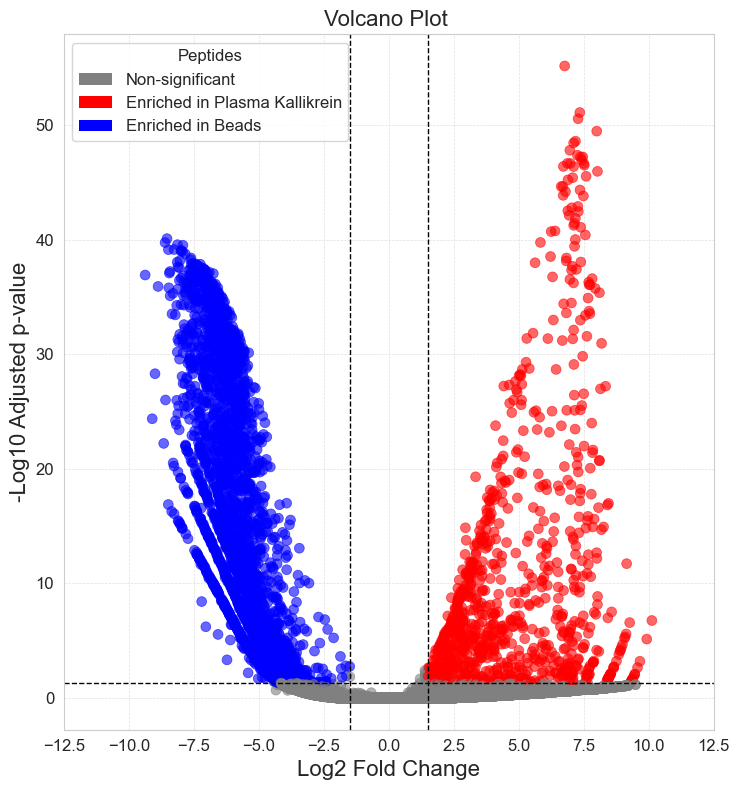

***Supplementary Figure 27: Volcano plot of peptide sequences from the third round of panning against plasma kallikrein and a streptavidin bead control.*** *Dashed lines indicate the significance thresholds (adj. p < 0.05, log_2_FC > 1.5). 895 sequences were statistically significantly enriched against plasma kallikrein versus the streptavidin bead control.*

#### 4.8 Inhibition Plots for Peptides 10 and 13

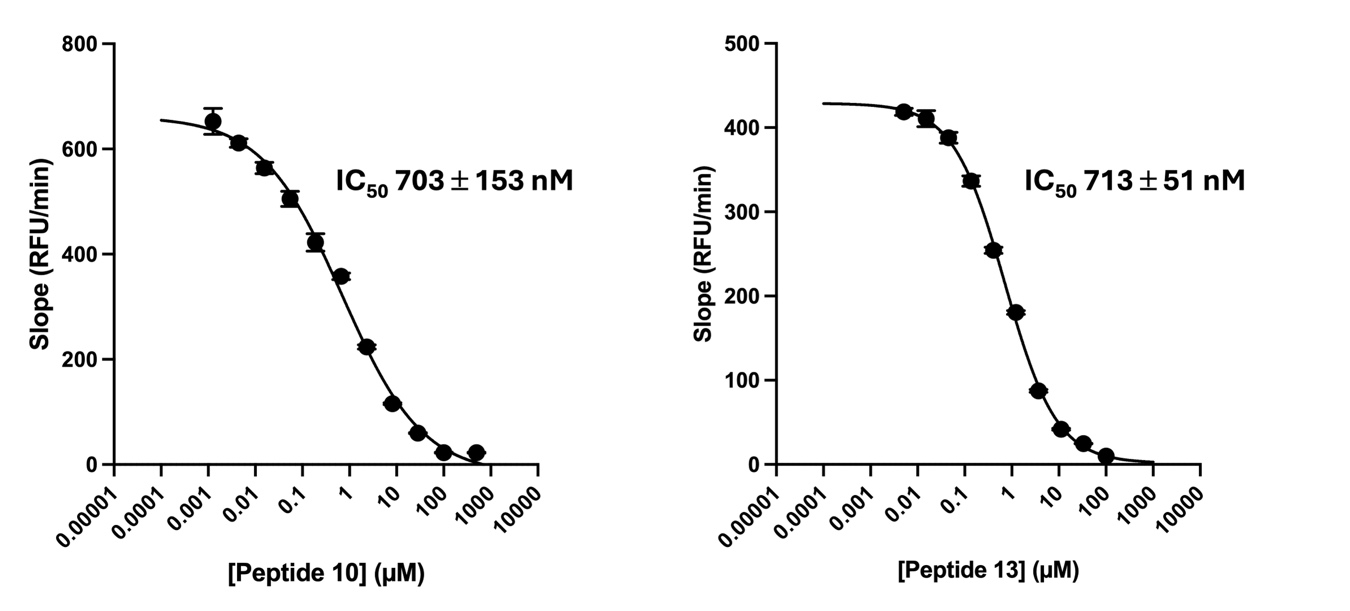

***Supplementary Figure 28: Determination of inhibition constants (IC_50_) for peptide 10 and 13.*** *Dose-response curves for human plasma kallikrein activity in the presence of increasing concentrations of peptides* ***10*** *and* ***13****. Enzyme activity is plotted as the slope of the kinetic progress curves. Data points indicate the mean ± SD of three independent replicates. Solid lines represent the non-linear regression fit to a standard dose-response model using GraphPad Prism.*

#### 4.9 Peptide 2 Isomer Characterisation with NaCNBH_3_ Reduction

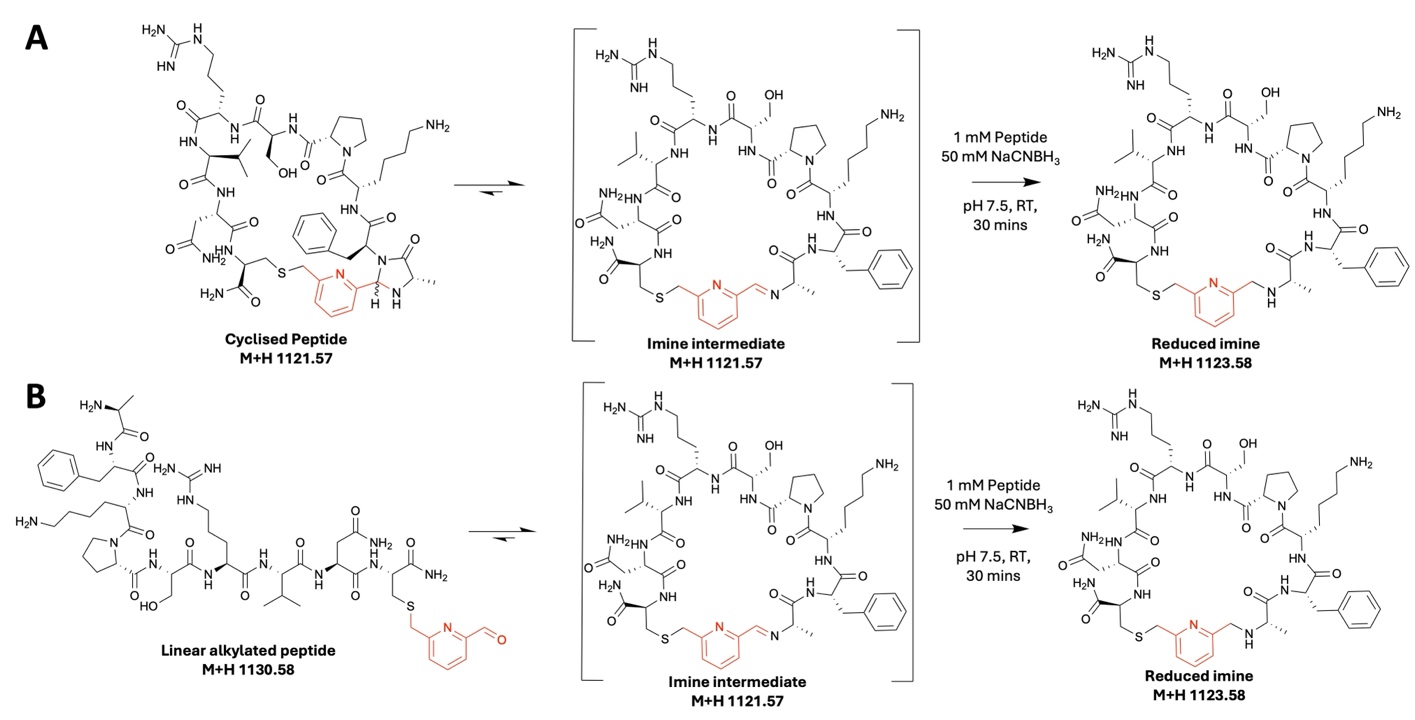

***Supplementary Figure 29: Scheme for characterisation of Peptide 2 cyclised products by NaCNBH_3_ reduction.*** *Two peaks with mass corresponding to the cyclised product were separated.* ***(A)*** *Reduction of the purified product peaks with NaCNBH_3_ would enable irreversible capture and hence identification of any imine intermediate present.* ***(B)*** *Reduction of the linear alkylated peptide was performed under the same conditions as the cyclised peptide to confirm the conditions were suitable for capturing imine intermediate present or formed during the period of the reduction.*

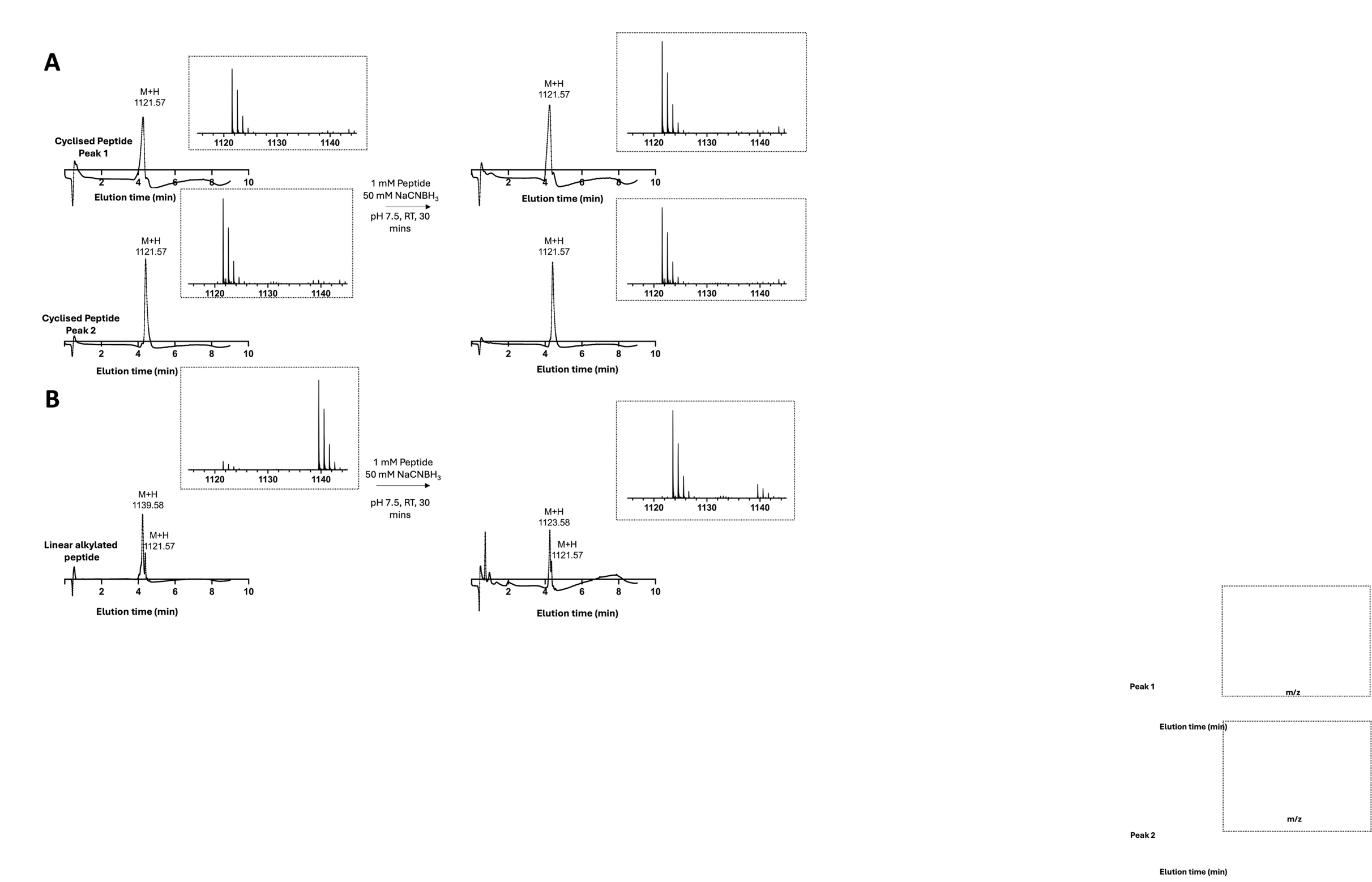
***Supplementary Figure 30: High-resolution LC-MS results for characterisation of peptide 2 by NaCNBH_3_ reduction. UV absorbance traces at 280 nm and the positive ionisation mass spectra for the primary UV peak are presented.*** ***(A)*** *Results for the purified cyclised product peaks. Neither peak shows a mass shift following NaCNBH_3_ reduction, suggesting each peak corresponds to an imidazolidinone diastereomer.* ***(B)*** *Results for the linear alkylated product. No linear alkylated product remains following reduction, only reduced imine and the small amount of cyclised peptide also present in the sample prior to reduction.*

#### 4.10 NMR Characterisation of Peptide 2a

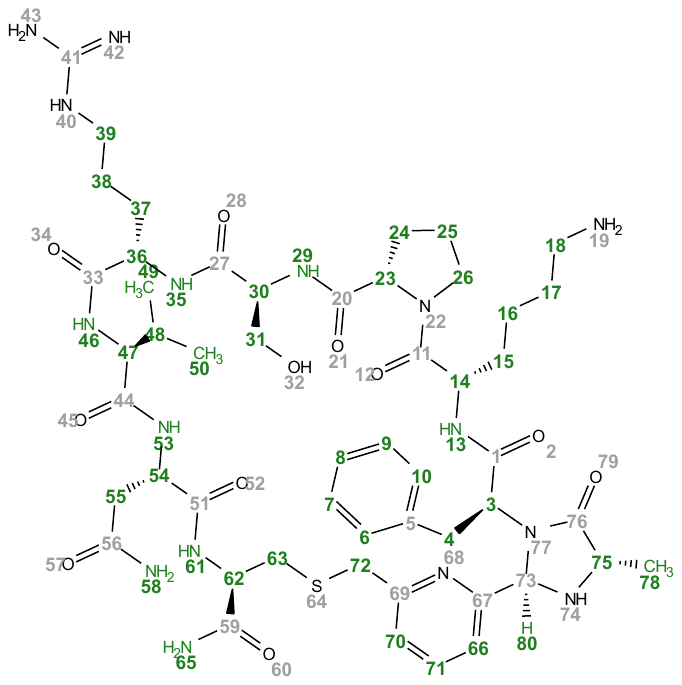

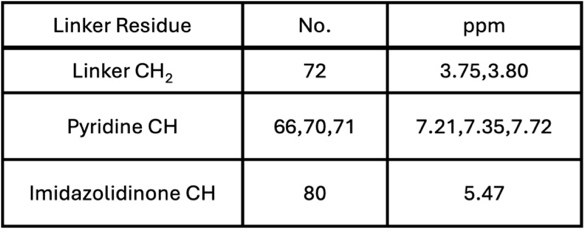

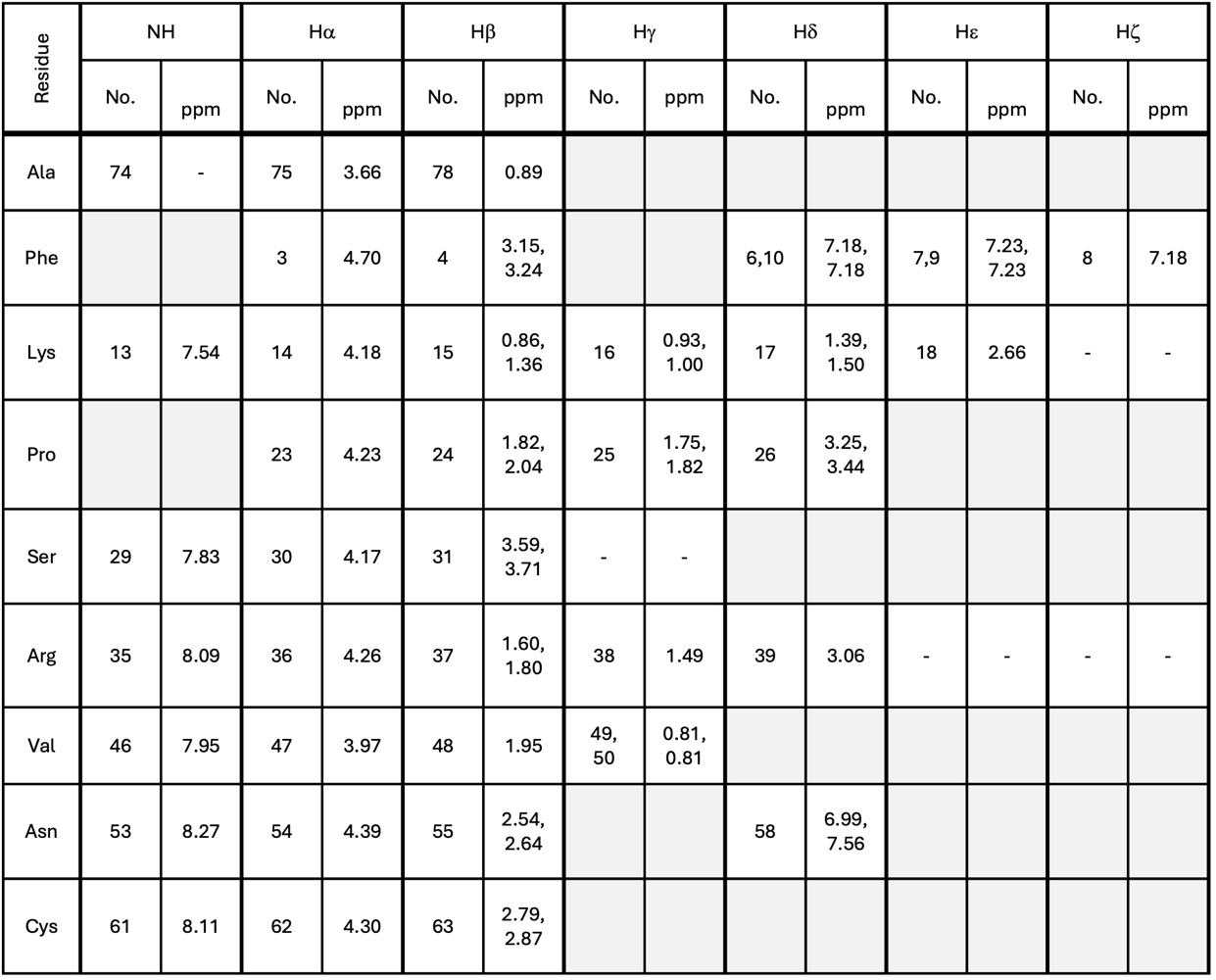

***Supplementary Figure 31: ^1^H NMR chemical shift assignments for ^1^H NMR of the R- isomer of cyclised Peptide 2, with IC_50_ 56 nM.***

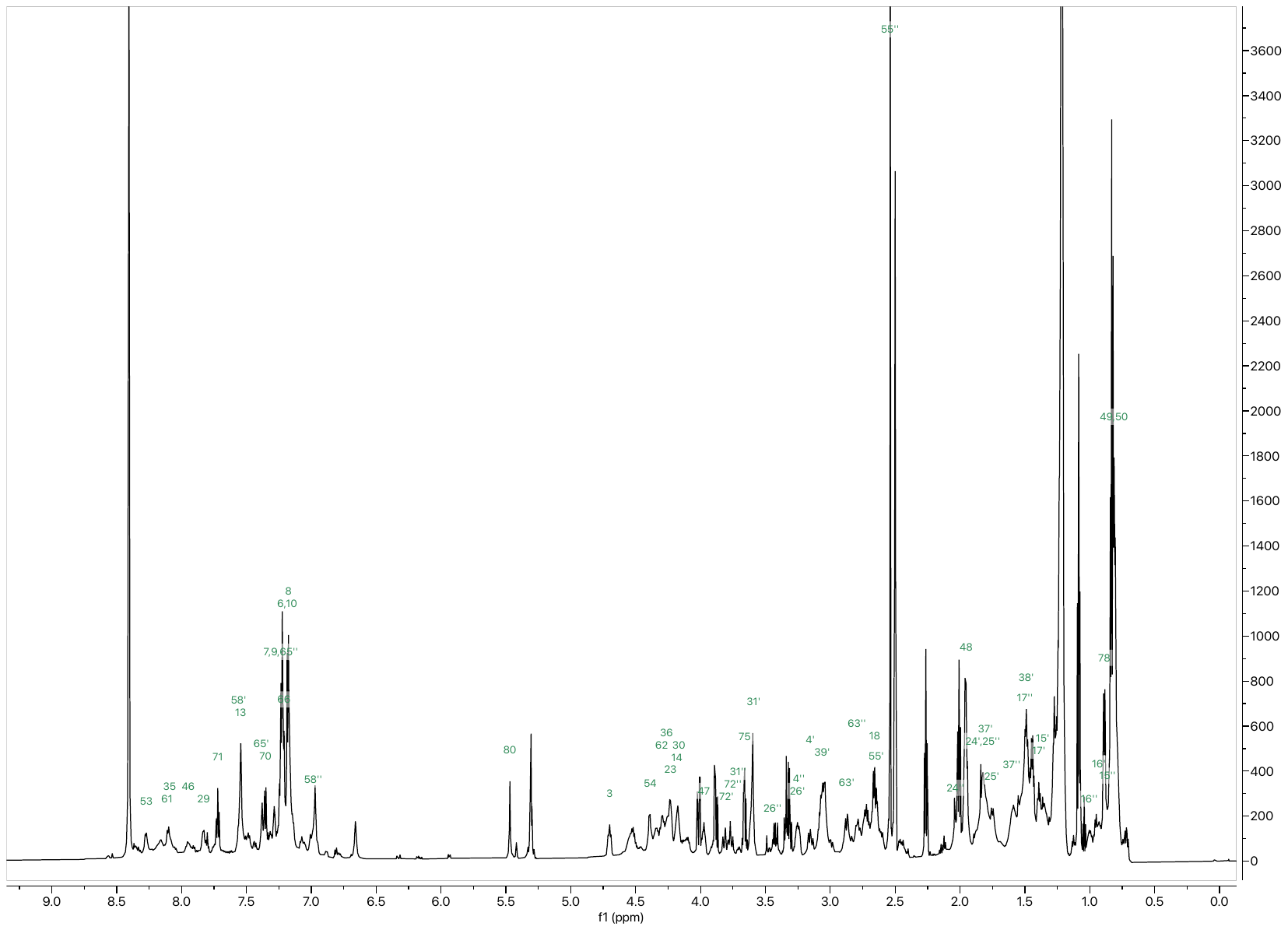
***Supplementary Figure 32: ^1^H spectrum of the R-isomer of cyclised Peptide 2, with IC50 56 nM.*** *Peptide was dissolved in d_6_-DMSO (10 mM) and spectra acquired at 298K on a Bruker Avance III HD spectrometer operating at 700 MHz equipped with a 1.7 mm TCI microcryocoil probe.*

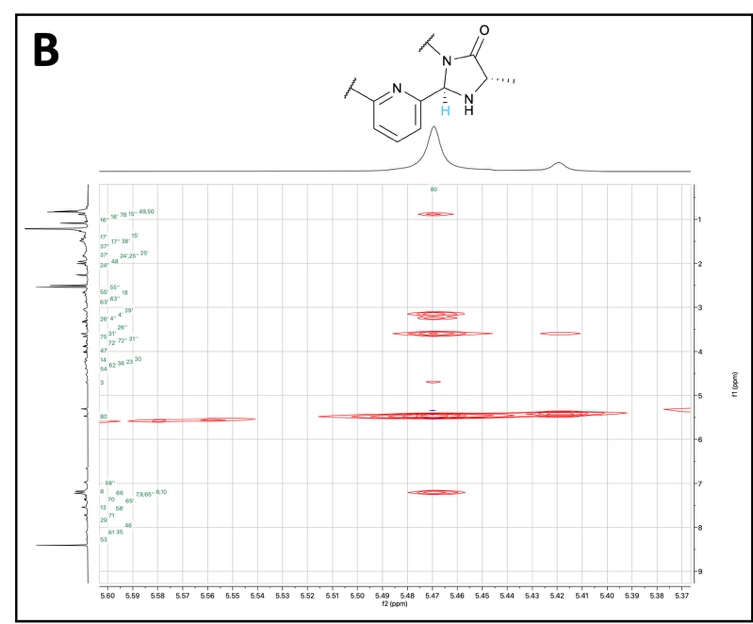

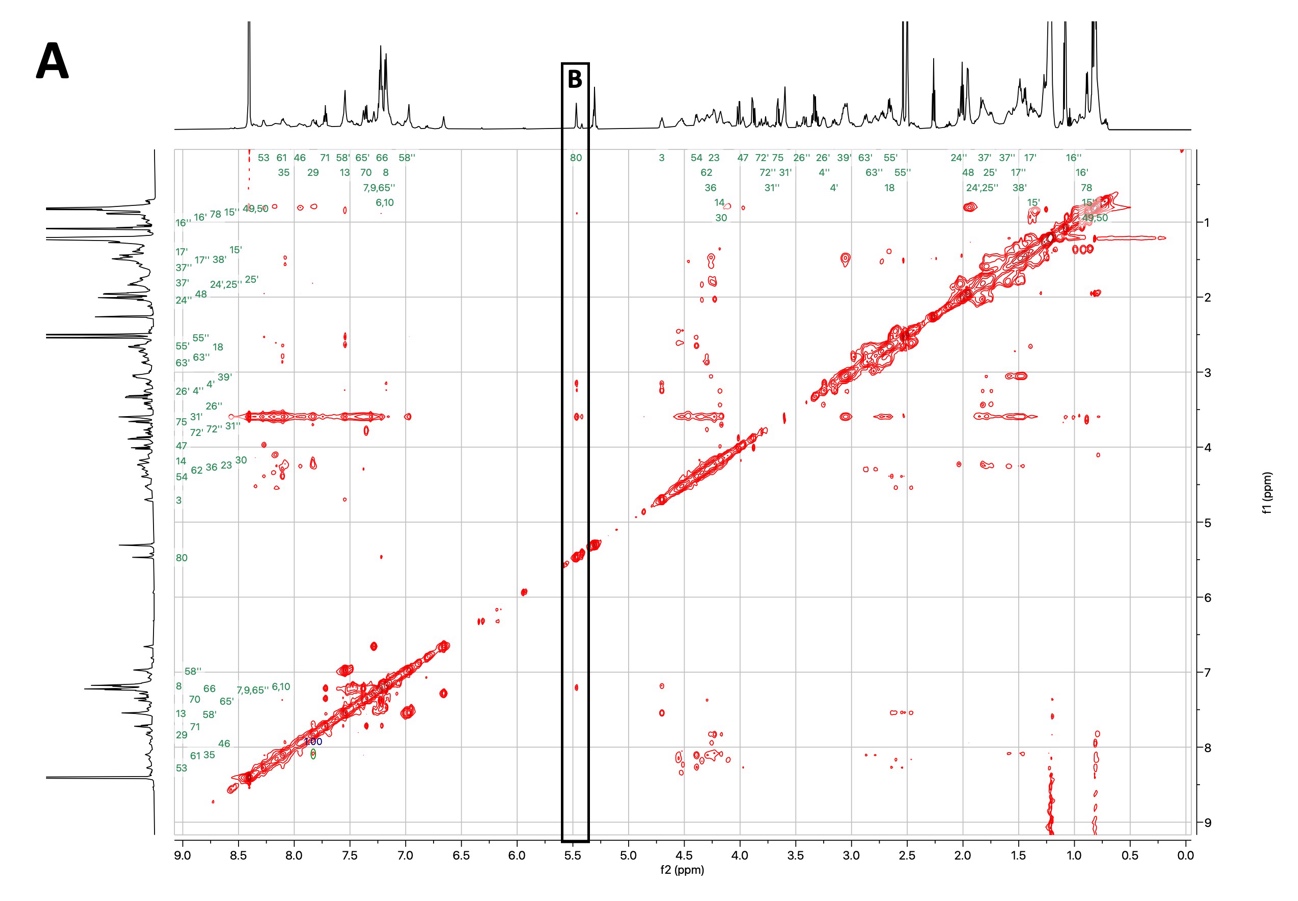

***Supplementary Figure 33: NOESY spectrum of the R-isomer of cyclised Peptide 2. (A)*** *Full spectra.* ***(B)*** *Excerpt showing characteristic through-space interactions with the imidazolidinone/diastereomeric proton.*

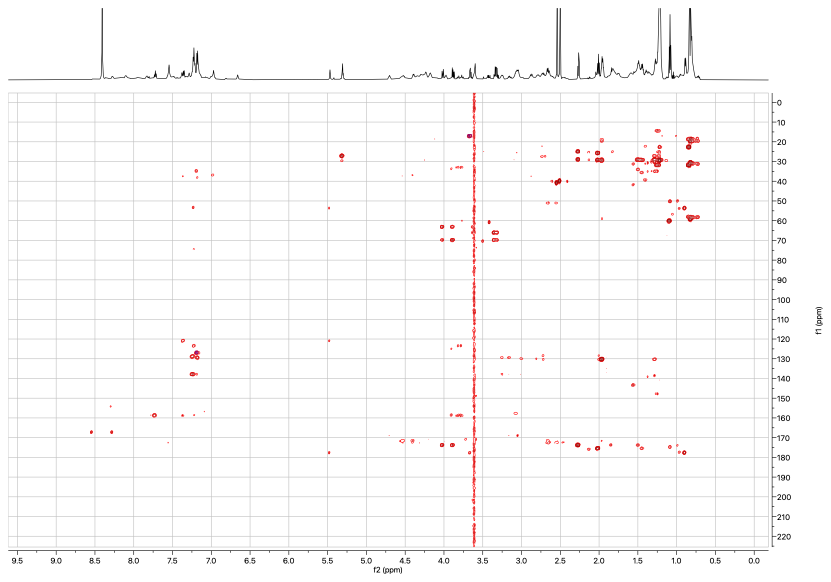

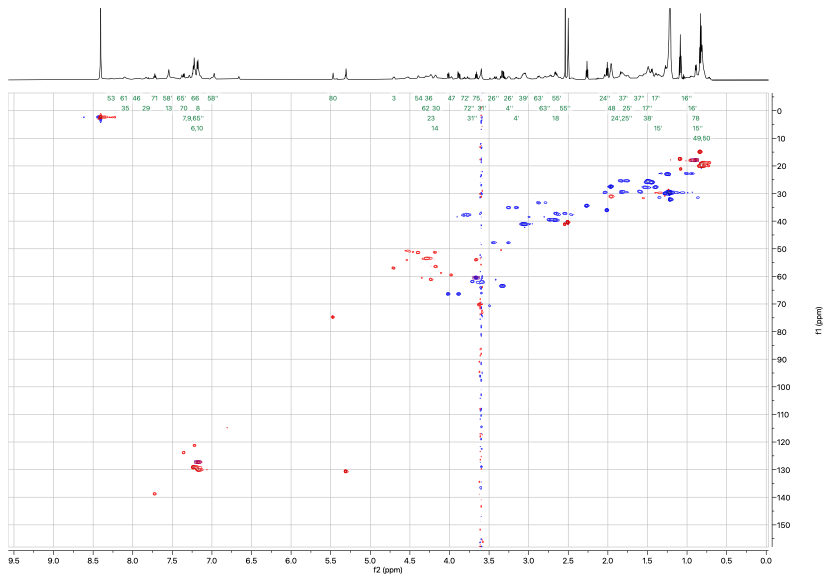
***Supplementary Figure 34: HSQC spectrum of the R-isomer of cyclised Peptide 2.***

***Supplementary Figure 35: HMBC spectrum of the R-isomer of cyclised Peptide 2.***

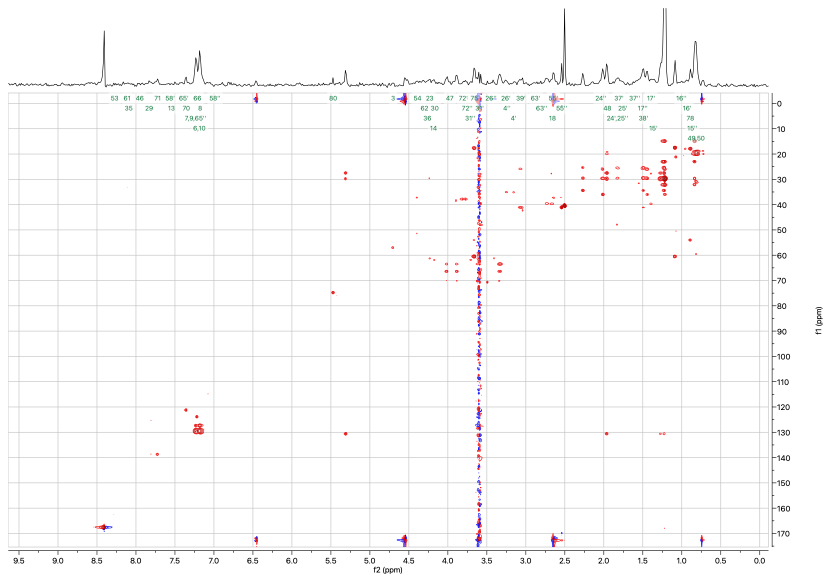

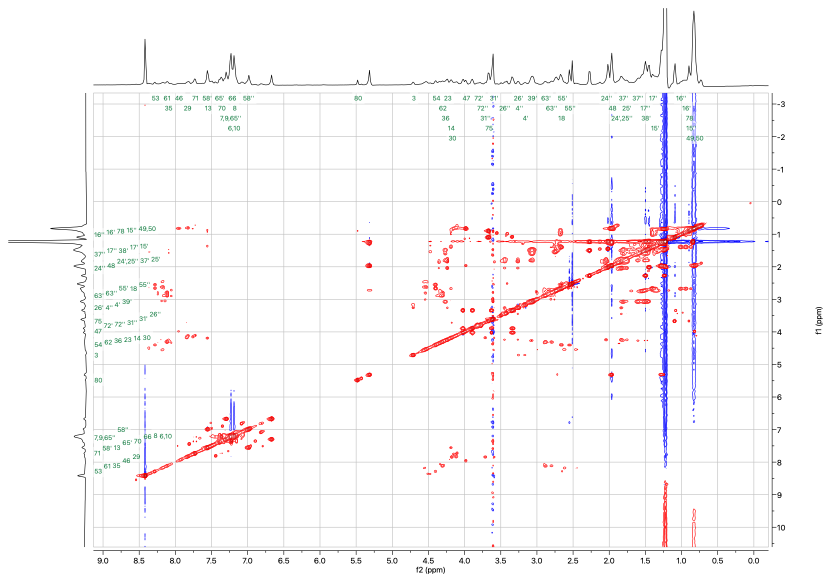
***Supplementary Figure 36: TOCSY spectrum of the R-isomer of cyclised Peptide 2.***

***Supplementary Figure 37: HSQC-TOCSY spectrum of the R-isomer of cyclised Peptide 2.***

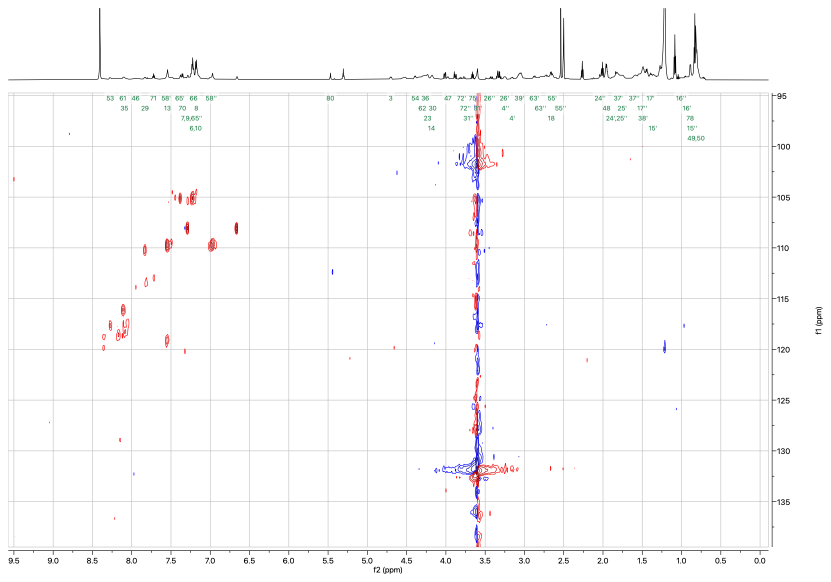
***Supplementary Figure 38: ^15^N-HSQC spectrum of the R-isomer of cyclised Peptide 2.***

#### 4.11 Inhibition Constant Plots for Peptide 2 Isomers

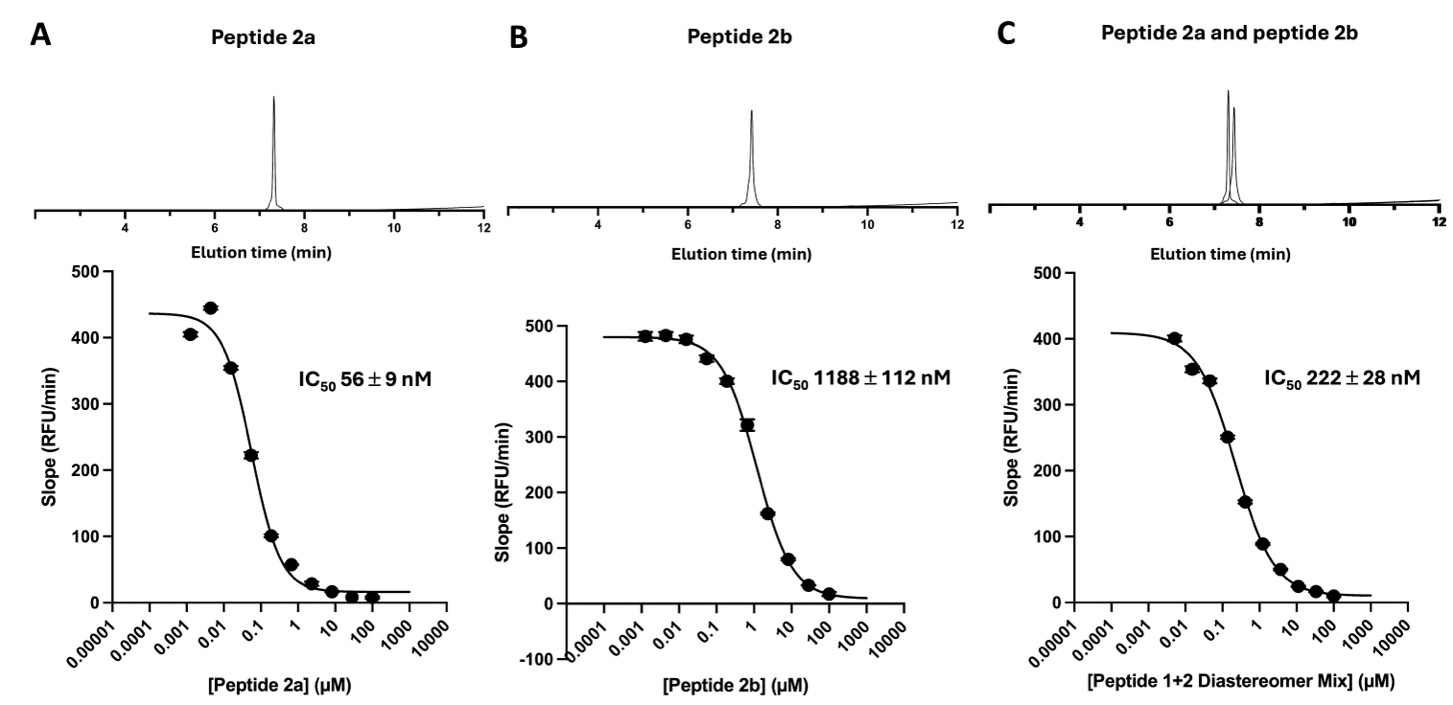

***Supplementary Figure 39: Determination of inhibition constants (IC_50_) for peptide 2 diastereomers.*** *Dose-response curves for human PK activity in the presence of increasing concentrations of peptide* ***2*** *diastereomers. Peptide* ***2a*** *(R isomer)* ***(A)****, peptide* ***2b*** *(S isomer)* ***(B)*** *and a 50:50 mix of the two* ***(C)*** *are shown alongside their UV traces (254 nm) from analytical low-resolution LCMS. Enzyme activity is plotted as the slope of the kinetic progress curves. Data points indicate the mean ± SD of three independent replicates. Solid lines represent the non-linear regression fit to a standard dose-response model using GraphPad Prism.*

#### 4.12 Haemolysis Assay

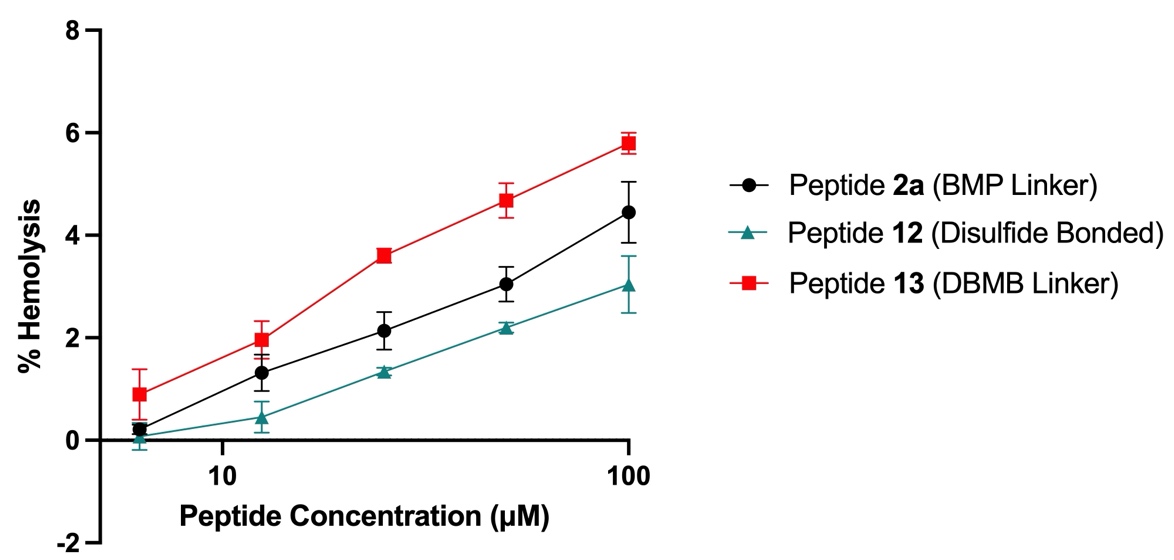

***Supplementary Figure 40. Haemolytic effect of different cyclisation strategies.*** *Percentage haemolysis induced after 1h incubation of a 5% (v/v) horse erythrocyte solution for the peptide sequence (A/C)FKPSRVNC cyclised with the BMP-linker (****2a****), a disulfide bond (****12****) or DBMB-Linker (****13****).*

#### 4.13 Intact Mass Spectrometry Analysis of Plasma Kallikrein Covalent Labelling by Peptide 14

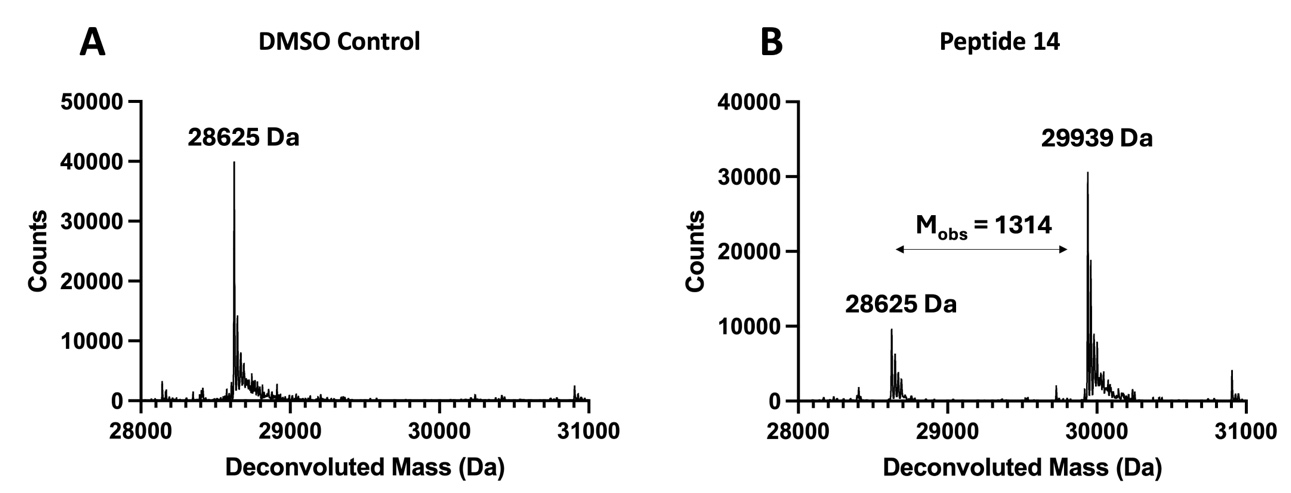

***Supplementary Figure 41. Intact mass spectrometry analysis of plasma kallikrein labelling by peptide 14****. Non-glycosylated PK (5 µM) was incubated with either a DMSO control* ***(A)*** *or peptide* ***14*** *(50 µM)* ***(B)*** *for 4 h at 37 °C before LC-MS analysis and biomolecular deconvolution. Incubation with peptide* ***14*** *resulted in a mass shift of 1314. M_exp_ for covalent modification by peptide* ***14*** *is 1313.5195.*

#### 4.14 In-gel Fluorescence of Serine Protease Panel Labelling

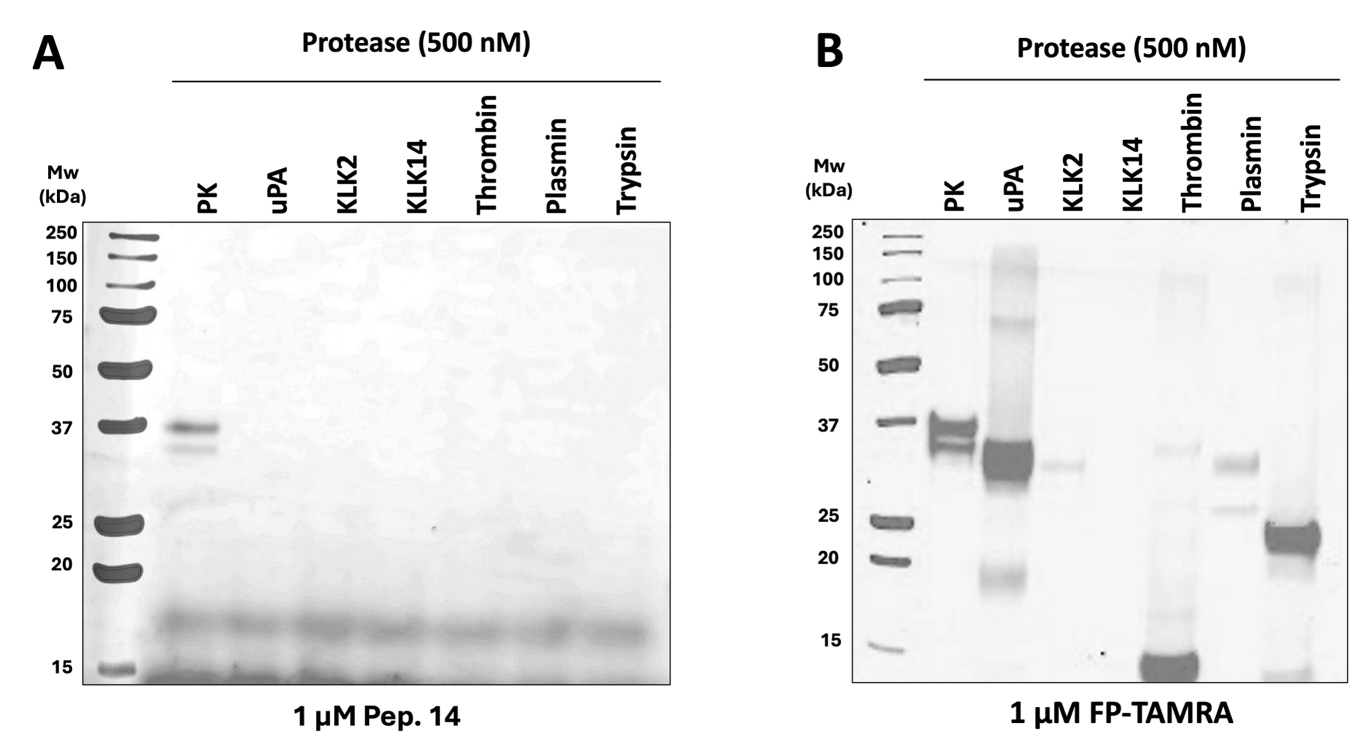

**Supplementary Figure 42. *In-gel fluorescence profiling of FP-TAMRA reactivity across several serine proteases.*** *Serine proteases (500 nM) were incubated with the broad-spectrum serine protease probe FP-TAMRA (1 μM) in PBS (pH 7.4) for 60 min at 37 °C. Following SDS-PAGE resolution, covalent labelling was visualised via in-gel TAMRA fluorescence. Protease abbreviations: PK, plasma kallikrein; uPA, urokinase-type plasminogen activator; KLK2, kallikrein-2; KLK14, kallikrein-14.*

#### 4.15 Peptide 14 Stability Data

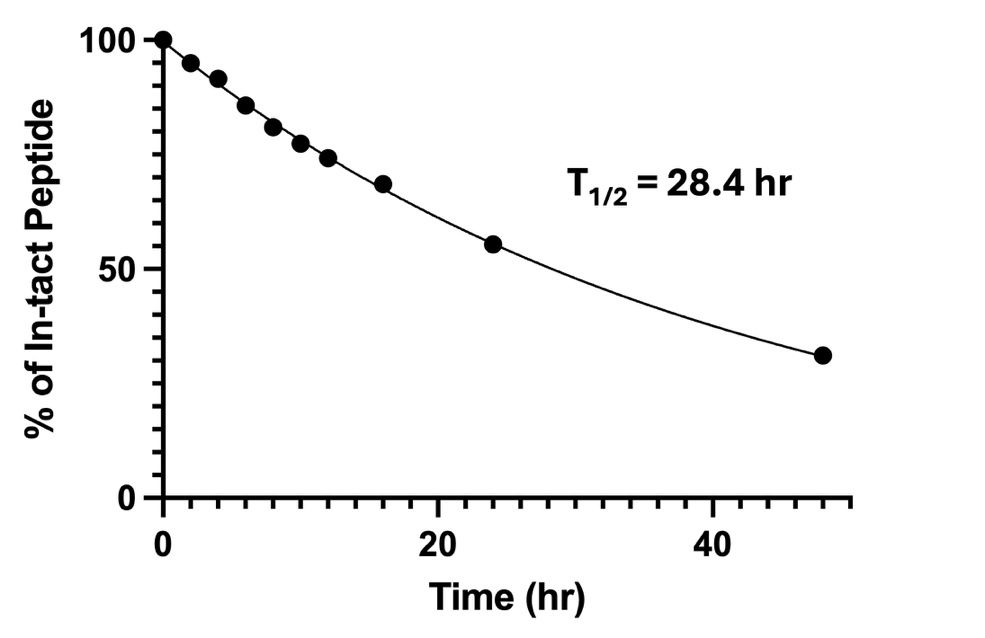

***Supplementary Figure 43. Stability of peptide 14 in PBS (pH 7.4).*** *Percentage intact peptide, normalised against an internal control, at timepoints over a 48 h period was determined by analytical LC-MS. A half-life of 28.4 h was determined by fitting the normalised percentage intact peptide to a one-phase exponential decay model.*

### 5 Macrocyclic Peptide Analytical Data

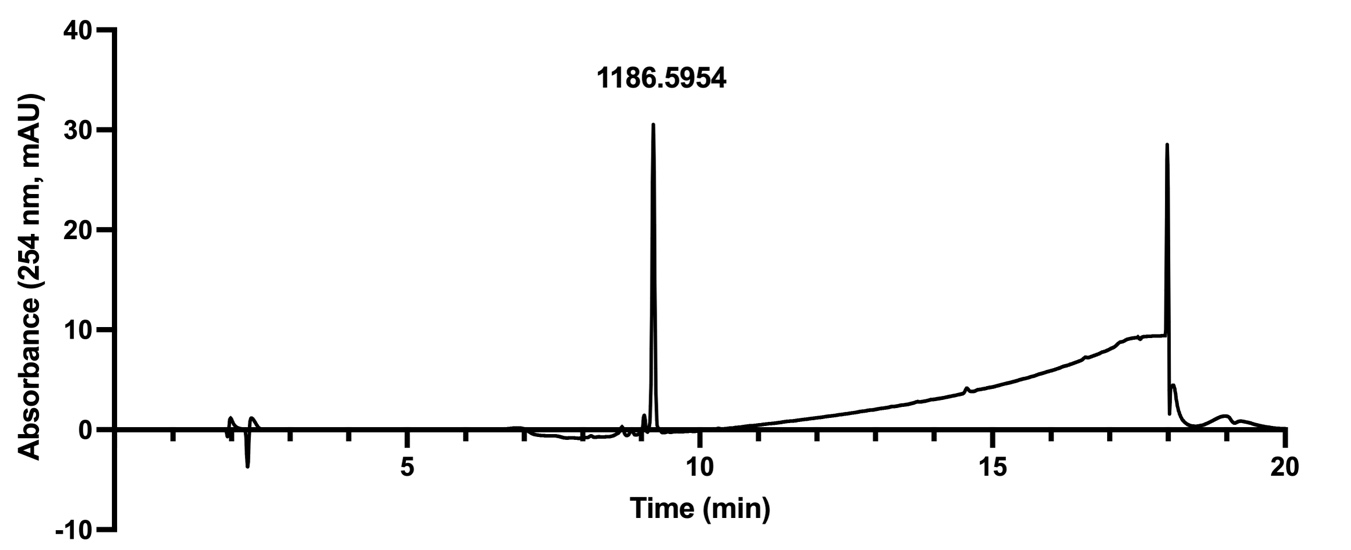
**Peptide 1**

**Chemical formula: [C_55_H_79_N_17_O_11_S]^+^**

**[M+H]^+^ calc:** **1186.5944**

**[M+H]^+^ obs:** **1186.5954**

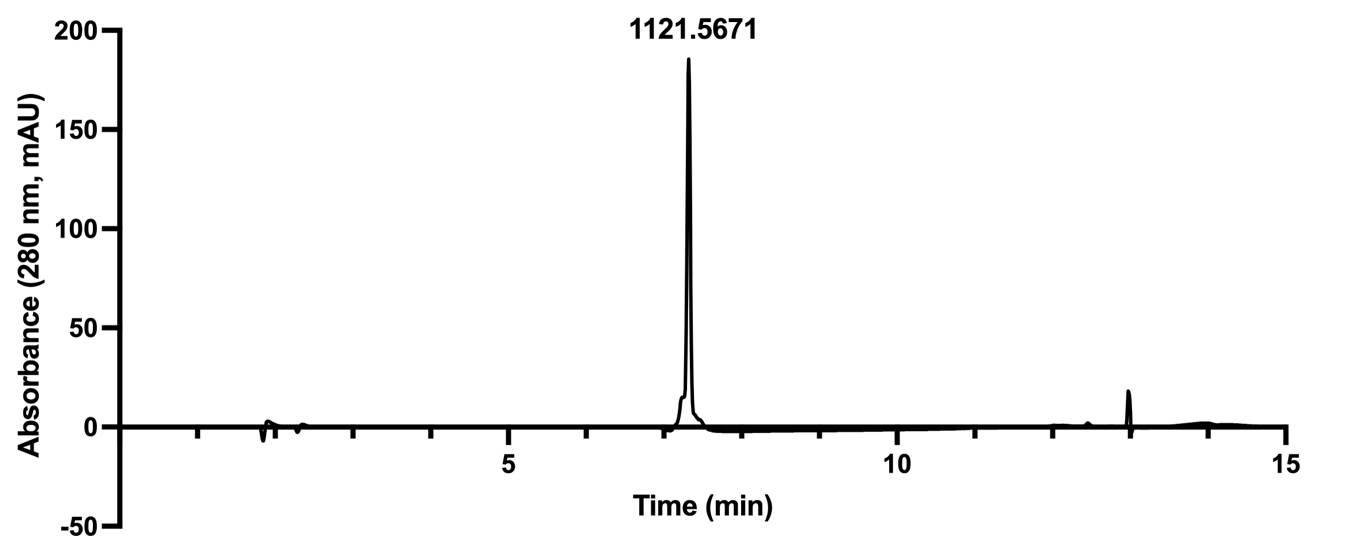
**Peptide 2a**

**Chemical formula: [C_51_H_77_N_16_O_11_S]^+^**

**[M+H]^+^ calc:** **1121.5678**

**[M+H]^+^ obs:** **1121.5671**

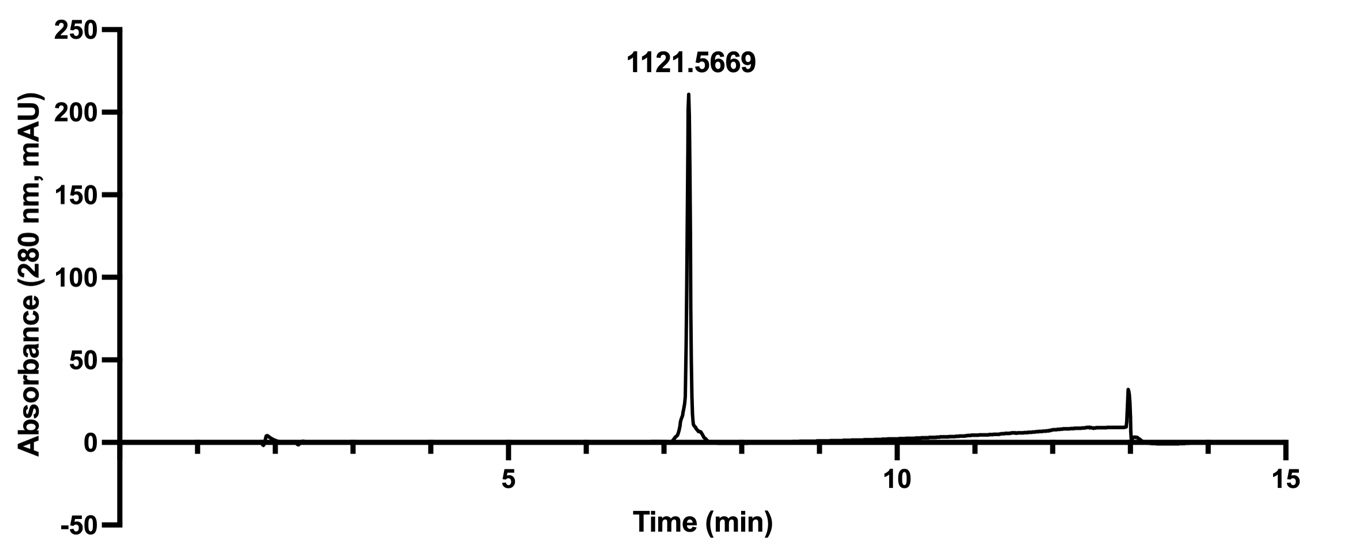

**Peptide 2b (R/S)**

**Chemical formula: [C_51_H_77_N_16_O_11_S]^+^**

**[M+H]^+^ calc:** **1121.5678**

**[M+H]^+^ obs: 1121.5669**

**Peptide 10**

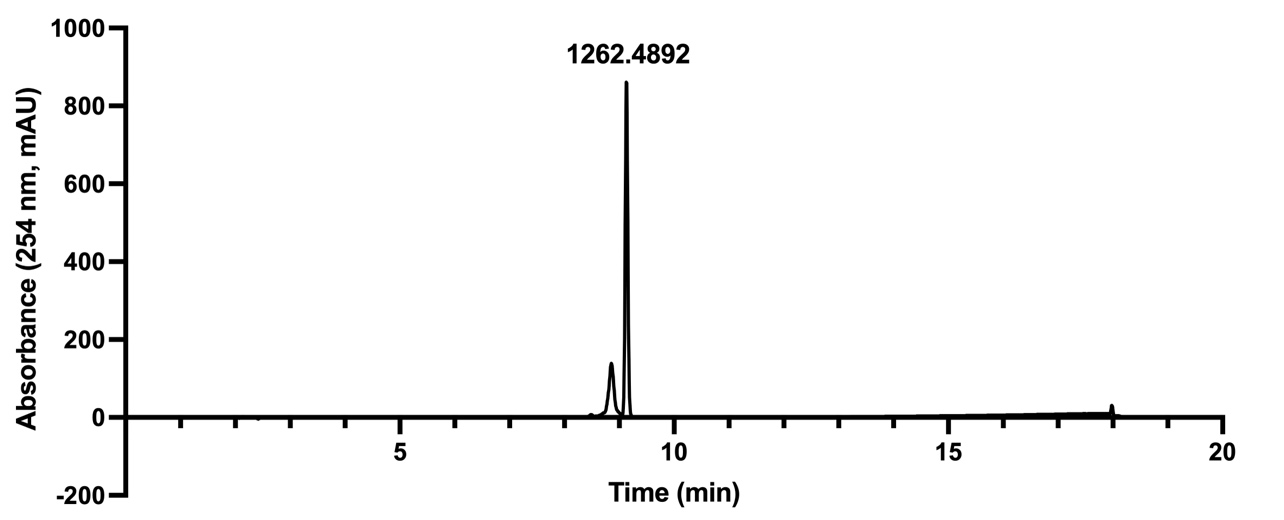

**Chemical formula: [C_58_H_83_N_18_O_10_S]^+^**

**[M+H]^+^ calc: 1223.6260**

**[M+H]^+^ obs: 1223.6269**

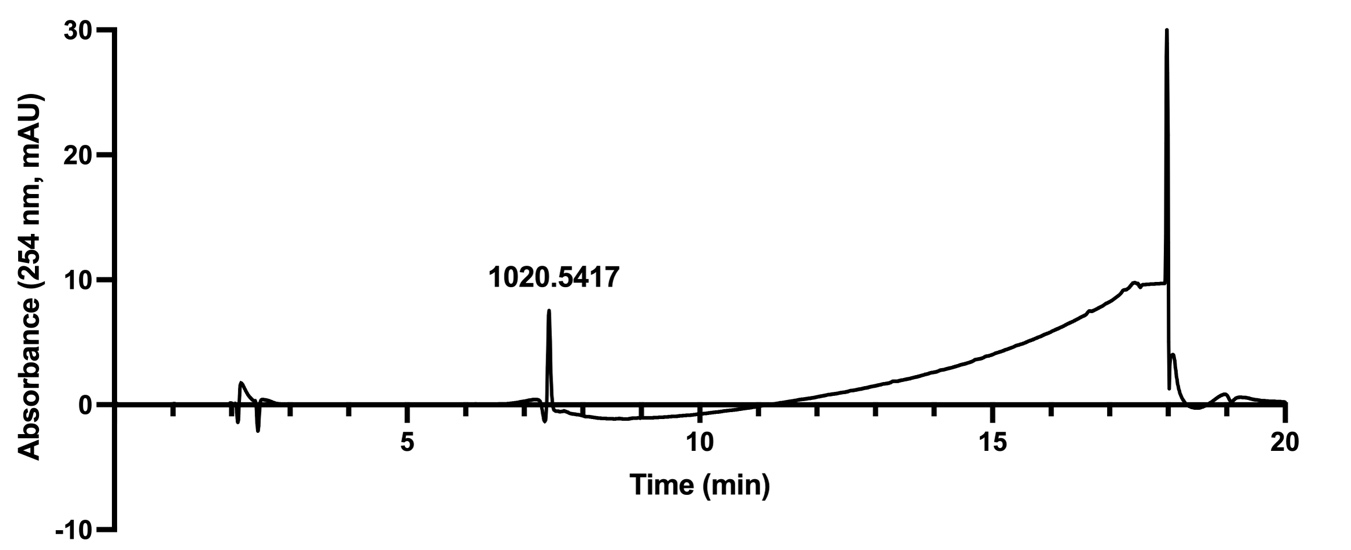
**Peptide 11**

**Chemical formula: [C_44_H_74_N_15_O_11_S]^+^**

**[M+H]^+^ calc: 1020.5413**

**[M+H]^+^ obs: 1020.5417**

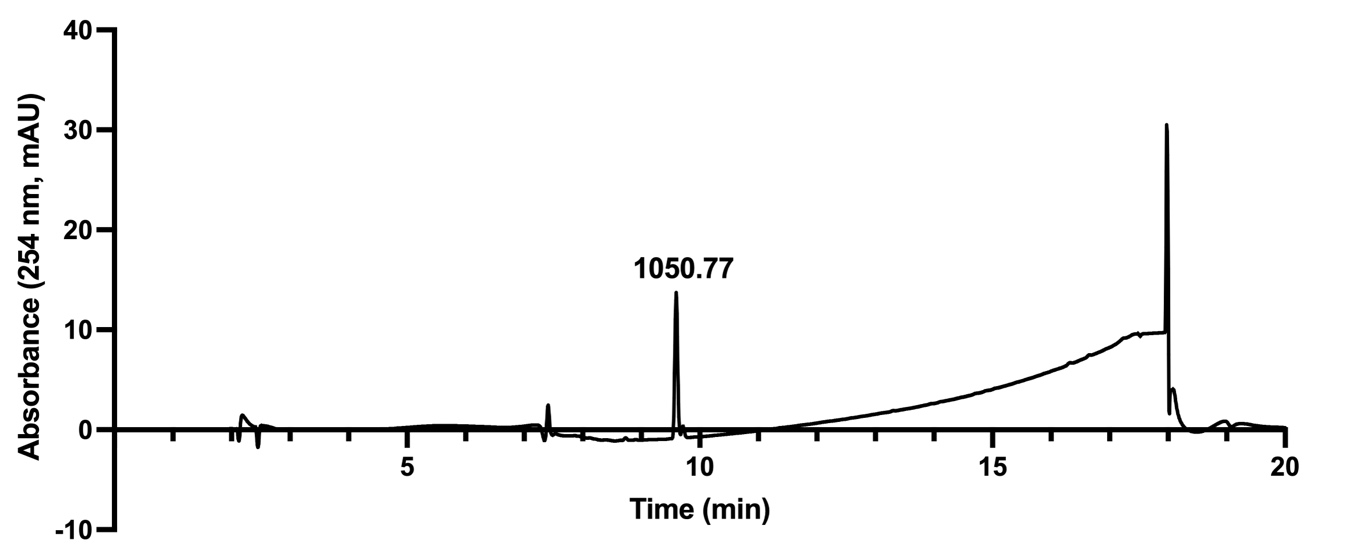
**Peptide 12**

**Chemical formula: [C_44_H_72_N_15_O_11_S_2_]^+^**

**[M+H]^+^ calc: 1050.4977**

**[M+H]^+^ obs: 1050.4977**

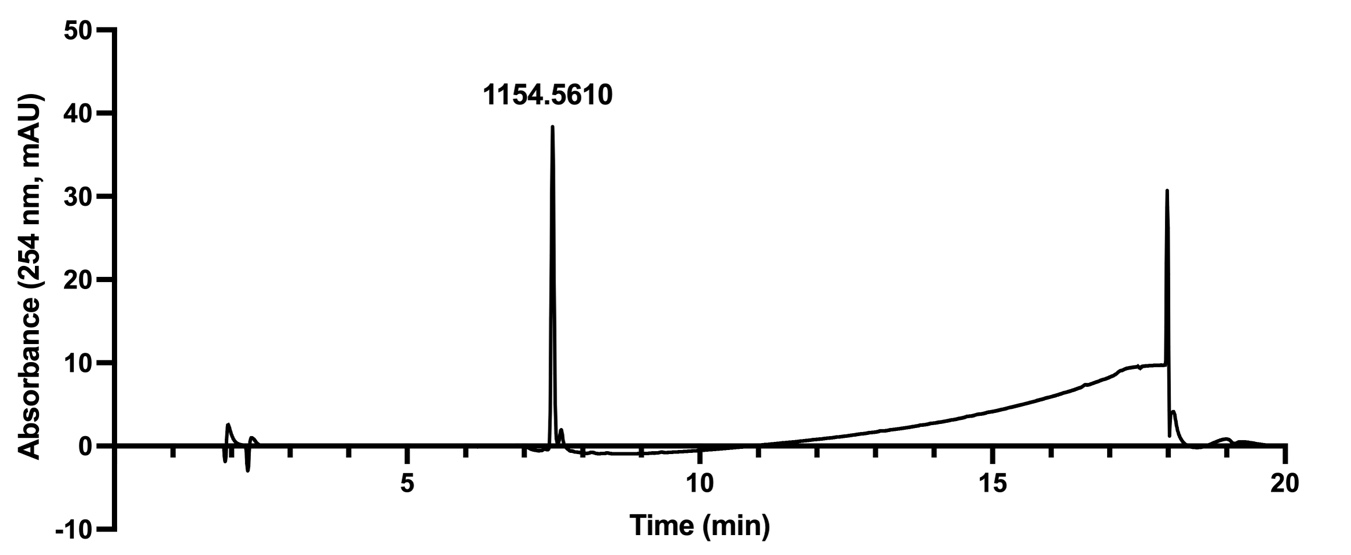
**Peptide 13**

**Chemical formula: [C_52_H_80_N_15_O_11_S_2_]^+^**

**[M+H]^+^ calc: 1154.5603**

**[M+H]^+^ obs: 1154.5610**

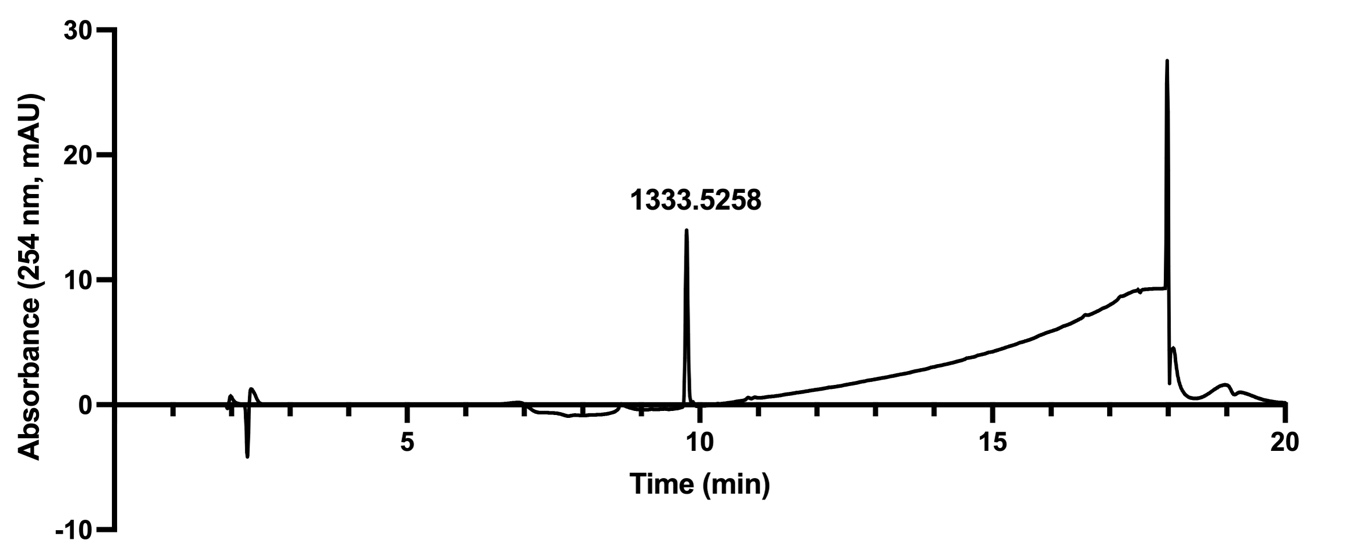
**Peptide 14**

**Chemical formula: [C_59_H_78_FN_16_O_15_S_2_]^+^**

**[M+H]^+^ calc: 1333.5258**

**[M+H]^+^ obs: 1333.5258**

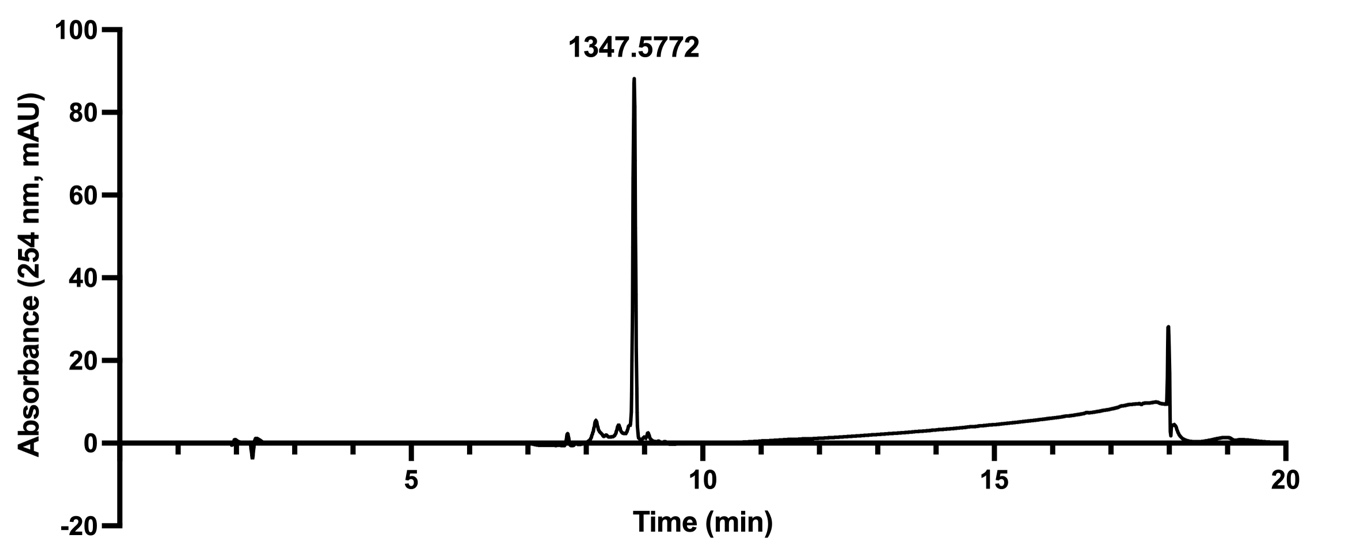
**Peptide 15**

**Chemical formula: [C_61_H_84_FN_16_O_14_S_2_]^+^**

**[M+H]^+^ calc: 1347.5778**

**[M+H]^+^ obs: 1347.5772**

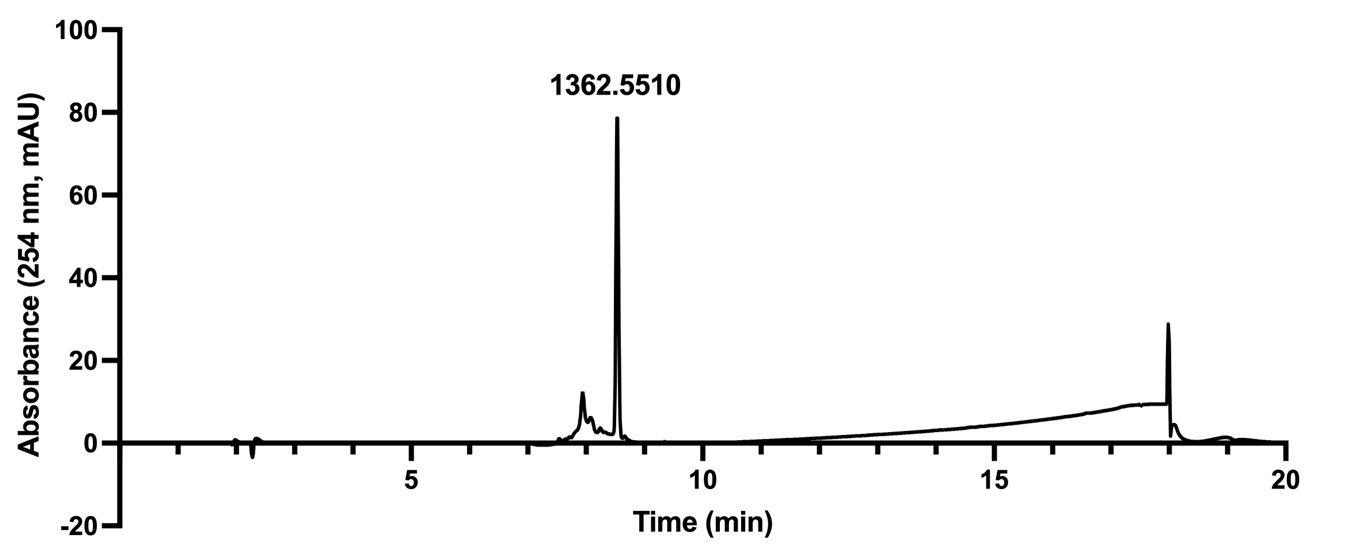
**Peptide 16**

**Chemical formula: [C_60_H_81_FN_17_O_15_S_2_]^+^**

**[M+H]^+^ calc: 1362.5524**

**[M+H]^+^ obs: 1362.5510**

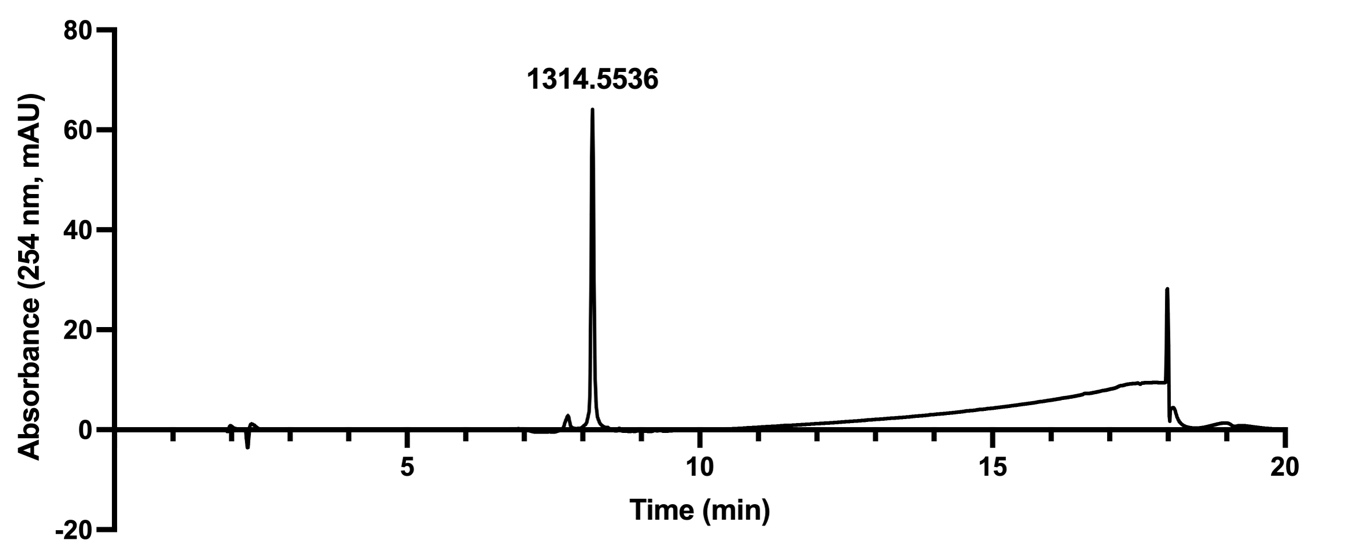
**Peptide 17**

**Chemical formula: [C_56_H_81_FN_17_O_15_S_2_]^+^**

**[M+H]^+^ calc: 1314.5523**

**[M+H]^+^ obs: 1314.5536**

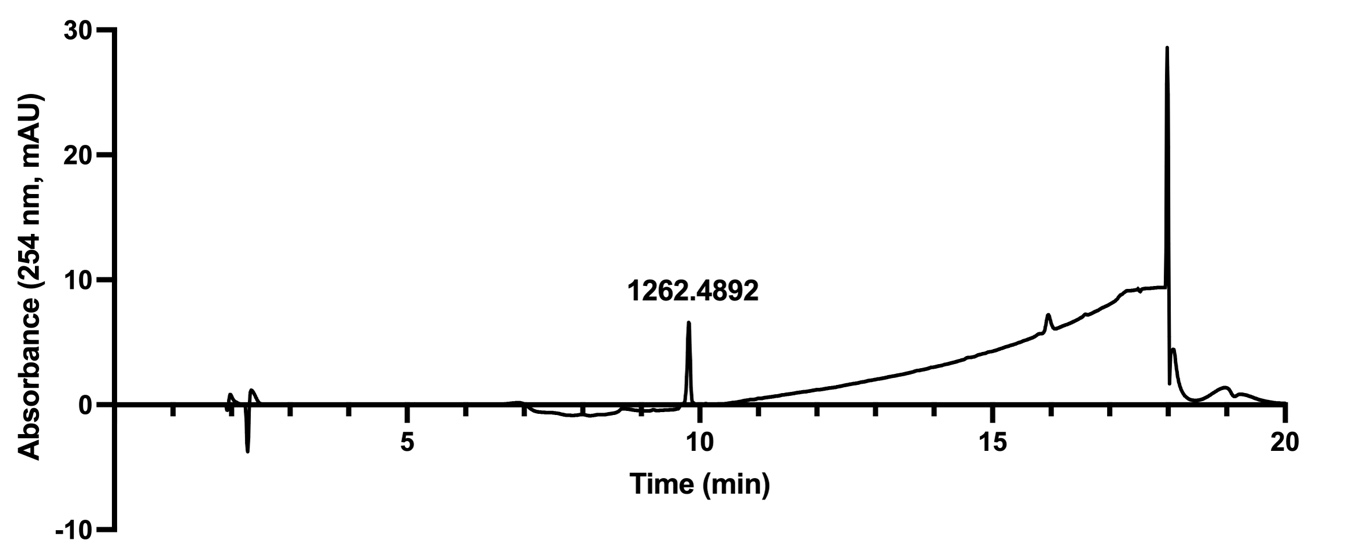
**Peptide 18**

**Chemical formula: [C_56_H_73_FN_15_O_14_S_2_]^+^**

**[M+H]^+^ calc: 1262.4887**

**[M+H]^+^ obs: 1262.4892**
